# What and When Local Predictors Drive Tadpole Diversity in Subtropical Temporary Ponds?

**DOI:** 10.1101/2020.03.27.978338

**Authors:** Diego Anderson Dalmolin, Tiago Gomes dos Santos, Alexandro Marques Tozetti, Maria João Ramos Pereira

**Affiliations:** Programa de Pós-Graduação em Biologia Animal, Universidade Federal do Rio Grande do Sul, Porto Alegre, Rio Grande do Sul, Brasil; Universidade Federal do Pampa, São Gabriel, Rio Grande do Sul, Brasil; Universidade do Vale do Rio dos Sinos, São Leopoldo, Rio Grande do Sul, Brazil; Centre for Environmental and Marine Studies, Universidade de Aveiro, Aveiro, Portugal

**Keywords:** Amphibians, community assembly, depth gradients, environmental filtering, seasonality, functional, phylogenetic diversity

## Abstract

We evaluated seasonal variation in taxonomic, functional and phylogenetic diversity and redundancy of tadpoles in 401 points of 10 ponds in southern Brazil. We predicted i) congruent patterns between all components of diversity and environmental descriptors; ii) stronger effects of environment in the diversity components in seasons when the water level in ponds is low; iii) diversity components to be influenced by distinct sets of environmental factors in different periods. Predictions were tested using Linear Mixed Models. We observed positive influence of water depth on taxonomic, functional and phylogenetic diversity, as well as on functional redundancy during periods when the water level in ponds is high. Phylogenetic redundancy was not explained by any of the selected environmental variables. When the water level in ponds is low none of the environmental descriptors affects any of the diversity components. Environmental filtering seems to strongly influence tadpole community structure in temporary ponds, at least in periods when water depth gradients create a variety of micro-habitats allowing diverse sets of species to settle and co-occur. These species sets are then filtered according to their swimming and foraging abilities along the depth gradient, where intermediate depths should contain the greatest tadpole diversity.

## INTRODUCTION

Different niche-based processes influence aquatic communities, suggesting that biological communities do not result exclusively from events of dispersion and stochasticity (Petchey et al., 2007; Villéger et al., 2010, Mason et al., 2011; de Bello et al., 2013), but are also the result of biotic interactions and, mostly, environmental filtering. In this process, environmental factors act as a filter, selecting the species of the regional pool (see the recent debate on the environmental filtering mechanism in Kraft et al., 2015, and Cadotte & Tucker, 2017), so that the local community composition results from the direct relation between the ecological characteristics of the species and the biotic and abiotic environment (Poff, 1997; Leibold et al., 2004; Hubbel, 2001). Thus, communities under equivalent environmental conditions should harbor species with similar ecological requirements and should be more similar to each other than expected at random (Parris, 2004; Heino et al., 2015).

Recent studies support the idea that research on community structure should not focus solely on taxonomic diversity, but also on a set of metrics encompassing different aspects of diversity that are complementary to each other (Díaz et al., 2007; Mouchet et al., 2010; Cavender-Bares et al., 2009). A more accurate evaluation of functional attributes associated with environmental descriptors and the cumulative evolutionary history in each community is achieved by evaluating functional and phylogenetic diversity (diversity of traits and diversity of evolutionary lineages, respectively; Villéger et al., 2012; de Bello et al., 2013; Corbelli et al., 2015). These aspects are crucial, as they tend to reflect the resilience of a system to environmental change (Petchey & Gaston, 2006; Mouchet et al., 2010; Meynard et al., 2011). Incorporating information on the evolutionary history of the species through phylogenetic diversity allows for the evaluation of the contribution of historical processes (e.g. extinction and speciation events) to the assembly of present-day communities, even when these processes act at larger spatial scales (Faith, 2016). Frequently patterns in the three components of diversity tend to be consistent (Erös et al., 2009; Meynard et al., 2011; Arnan et al., 2017), but these relationships between components vary according to the taxonomic group and degree of niche conservatism of the traits (Cavender-Bares et al., 2009; Arnan et al., 2015; Sobral & Cianciaruso, 2016). Thus, there are cases when the evaluated metrics may vary differentially along the same environmental gradients (Devictor et al., 2010; Safi et al., 2011; Corbelli et al., 2015).

Temporary ponds are part of the huge diversity of freshwater environments (Williams et al., 2004). The effect of environmental filtering on the community structure of these ponds is dependent on seasonal changes in the hydroperiod – period during which the pond retains water – and other local environmental conditions (Williams, 2006; Mendez et al., 2012; Ruhi et al., 2013). The filtering process is generally accentuated when the water level in ponds is low. Temperature and evapotranspiration rates, for example, tend to increase, constraining more severely the set of species able to persist in the ponds (Peltzer & Lajmanovich, 2004; Williams, 2006; Hamerlik et al., 2014). So, hydroperiod tends to define the composition of the community, with evolutionarily convergent and more functionally redundant species co-occurring when the water level in ponds is low (clustering pattern; Ruhi et al., 2013). On the other hand, functional and phylogenetically more divergent species co-occur when the water level in ponds is high, because temperature, dissolved oxygen and pH are not extreme (Schriever & Lytle, 2016; Ruhí et al., 2014; Martins et al., 2015; Strauβ et al., 2016).

Tadpoles select specific micro-habitats along environmental gradients (Altig & Johnston, 1989). This behavior seems to reflect the morphology and the evolutionary history of each species (Marques & Nomura, 2015) and may be influenced by several environmental factors (Haramuda, 2007). In fact, aquatic vegetation and depth were considered the main predictors of species richness and functional and phylogenetic diversity of tadpoles in ponds and streams (Both et al., 2011b; Queiroz, da Silva & Rossa-Feres, 2015; Melo et al., 2017; Escoriza & Ben Hassine, 2017). The presence and structuring of aquatic vegetation promotes micro-spatial heterogeneity, increasing the availability of sites for foraging and for seeking refuge from predators (Eterovick & Fernandes, 2001; Alford et al., 1999; Hero et al., 2001; Kopp, Wachlevski & Eterovick, 2006), while depth gradients allow for the exploration of different levels along the water column and the maintenance of species in distinct periods of larval development (Welborn, Skelly & Werner, 1996; Both et al., 2011a; Escoriza & Ben Hassine, 2017). Water chemistry (dissolved oxygen, pH) and temperature may also influence tadpole diversity, as they induce physiological and behavioral responses that are determinant in relationships with predators and competitors (Warner, Dunson & Travis, 1991). In drying pools pH levels generally increase due to the high concentration of ions and CO^2^ (Rowe, Sadinski & Dunson, 1992; Angélibert et al., 2004). Thus, tadpole survival rates should be expected to be different along the gradients of water chemistry (Moore & Towsend, 1998). Finally, tadpole distribution may also respond to pond morphology and its spatial distribution (Provete et al., 2014). Thus the effects of the physical and chemical descriptors of water can be irrelevant for the tadpole communitie assembly in ponds (Provete et al., 2014).

Few studies on tadpole community structuring have explored the influence of environmental filtering on the three components of diversity (taxonomic, functional, and phylogenetic) (Strauβ et al., 2016; Escoriza & Ben Hassine, 2017). Here, we aim to evaluate the relationships between local environmental descriptors and these metrics of diversity of tadpole communities in temporary ponds in southern Brazil. We evaluate if: i) local environmental factors act as predictors of tadpole diversity; ii) different components of diversity respond to the same environmental factors; iii) the influence of the environment on the diversity metrics changes seasonally. We expect congruent patterns between the three components of diversity and significant relationships with local environmental descriptors (Ribeiro et al., 2017; Escoriza & Ben Hassine, 2017). However, the patterns of diversity should be contrasting between seasons as a consequence of the reproductive phenology of the adults of some species that show seasonal peaks of activity related to some environmental conditions (e.g photoperiod – daylight length – and temperature; Both et al., 2008a). When the water level in ponds is high (Austral Spring and Winter), we expect a positive influence of aquatic vegetation, temperature and depth on the three components of diversity (Both et al., 2011a; Pujol-Buxó et al., 2017; Melo et al., 2017). Greater gradients of aquatic vegetation, temperature and depth favor the co-occurrence of benthic and nektonic species, which explore different resources according to their specific morphologies (Michel, 2011, Queiroz, da Silva & Rossa-Feres, 2015, Escoriza & Ben Hassine, 2017). However, when the water level in ponds is low (Austral Summer and Autumn) we expect a drastic change in pond environmental conditions, with shallow gradients in aquatic vegetation, temperature and depth, resulting in lower levels of diversity and higher levels of functional redundancy (Ruhí et al., 2014; Strauβ et al., 2016; Nunes et al. 2016). Diversities should be influenced by the environmental descriptors that affect the survival of tadpoles (Strauβ et al., 2016). Therefore, we expect that taxonomic, functional and phylogenetic diversities to be positively associated with dissolved oxygen levels and negatively associated with extreme temperature and pH values (Warner, Dunson & Travis, 1991).

## MATERIAL AND METHODS

### STUDY AREA

The study was done in the southern remnants of the Atlantic Forest Biome at the Reserva Biológica do Lami José Lutzemberger (30 ° 14’10.3 “S, 51 ° 05’51.7” W), an area with 204.04 hectares located on the banks of the Guaíba Lake, Porto Alegre, Rio Grande do Sul, Brazil (Figure 1). This is a conservation unit characterized by a low sandy plain landscape, formed by Quaternary sediments, and by a flat relief, which may present sandy elevations interspersed with depressions (Borges-Martins et al., 2013). Vegetation is typical of the ecotone between Semi-deciduous Seasonal Forest and Dense Ombrophylous Forest (Brack et al., 1998). Climate is mainly of the Cfa type (Wrege et al., 2011), characterized by average temperatures of 33°C in the warmest month and varying between 10 and 23°C in the coldest month (see the Table S1 of the Supplementary Material). Rainfall is well distributed throughout the year, with slightly higher rainfall volume between July (Austral Winter) and December (Austral Spring-Summer), a period of low water stress, while low levels of precipitation occur frequently between March (Austral Autumn-Summer) and May (Austral Autumn), a period of higher water stress, although there is no well-defined dry season. Annual average rainfall is 1500 mm (Wrege et al., 2011; Radin et al., 2017; Table S1 of the Supplementary Material).

**Figure 1:**
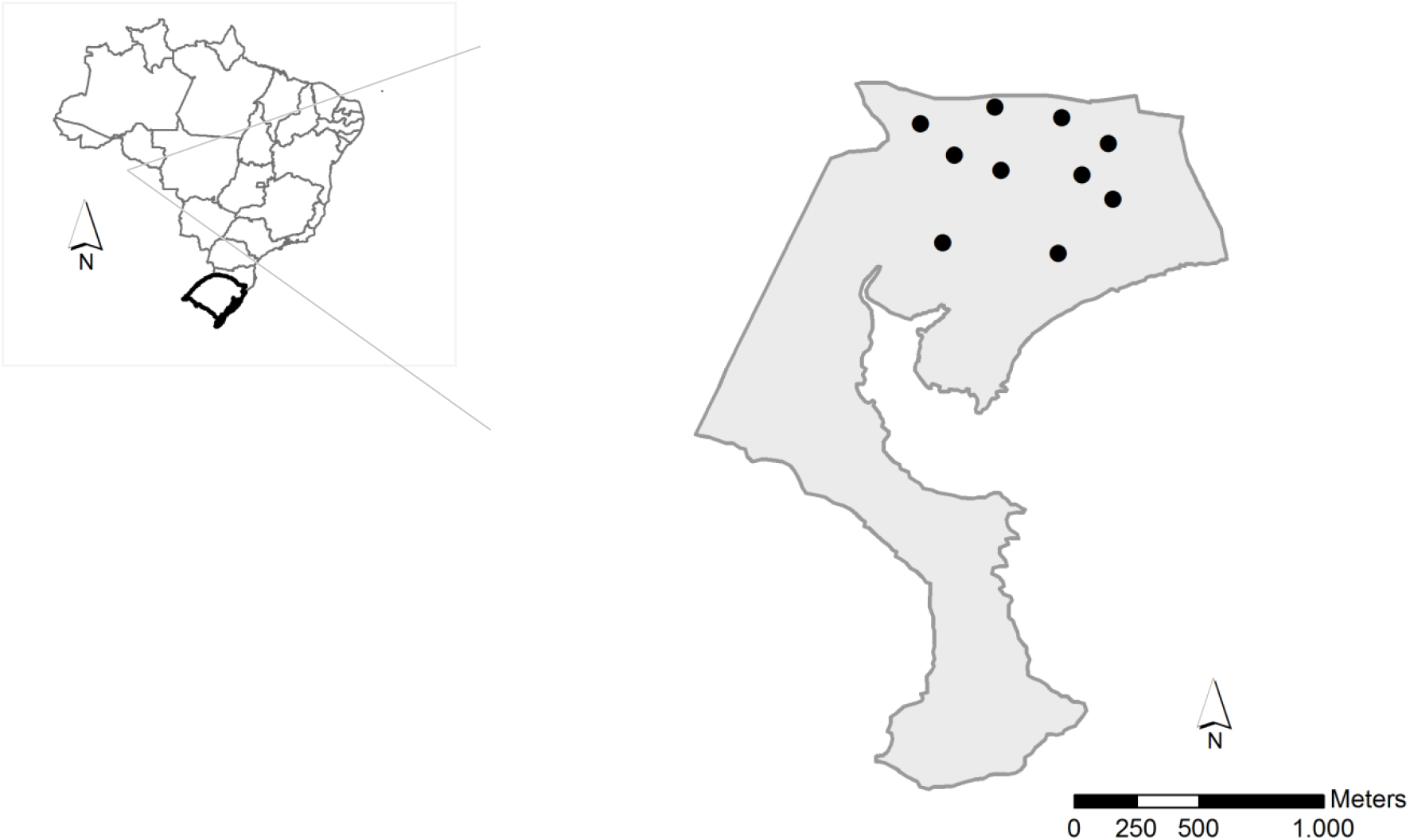
Location of the study area at Reserva Biológica do Lami in the Porto Alegre municipality, Rio Grande do Sul, Brazil. Dots indicate the 10 ponds sampled for tadpoles.

### DATA COLLECTION

The study was carried out in a group of small temporary water bodies (ponds) used as breeding sites by anurans. There are numerous ponds at the site with average depth of 50 centimeters when the water levels are high, surrounded by native vegetation cover composed mostly of grasslands associated with shrubs – predominantly of the Asteraceae and the Poaceae. Pond hydroperiod has two well-defined stages: low and high water levels, distributed in a non-predictive way throughout the year. We selected 10 ponds for sampling collection according to i) area accessibility and ii) distance between ponds of at least 250 meters. Data collection took place one time per month during anuran breeding season, between September 2013 and August 2014. Samples were collected in each pond at survey points at least three meters apart from each other (Prado et al., 2009). Delimitation of collection points and sampling methods of tadpoles followed Alford and Crump (1982), by marking each point to be sampled with a metal cylinder (70 centimeters long and 32 centimeters diameter), open in both ends. The metal cylinder was dropped to the chosen sampling point and, before tadpole collection, we checked if the lower end of the cylinder was well positioned on the pond substrate to prevent individuals from escaping. The individuals confined to the metal cylinder were collected with a wire mesh of 3 millimeters for a period of three minutes (more details on the sampling scheme in the Supplementary Material Figure S2). Each sampling point was resampled each month. Specimens collected in each sampling point were stored in separate flasks and identified in the laboratory to the species level with basis on literature and using specimens from scientific collections. The collected individuals were deposited in the Scientific Collection of the Zoology Department of the Universidade Federal do Rio Grande do Sul.

The characterization of the hydroperiod was done monthly by recording each pond’s individual total area and average depth. The surveyed ponds differed in relation to the environmental structure and the number of months they retained water (Table 1), as well as to the number of surveyed points (see Supplementary Material Table S3).

**Table 1.** Characterization of ponds (P1 – P10) sampled in the Reserva Biológica do Lami in relation to distribution of tadpoles in the period between September/2013 and August / 2014. Seasons: Spring (1); Summer (2); Autumn (3); Winter (4); Type of vegetation of the margin: Shrub (Sh.); underbrush (und.).

| Pond descriptors | P1 | P2 | P3 | P4 | P5 | P6 | P7 | P8 | P9 | P10 |
| --- | --- | --- | --- | --- | --- | --- | --- | --- | --- | --- |
| Total area (m²) | 50 | 205 | 20 | 810 | 85 | 54 | 15 | 64 | 112 | 84 |
| Average depth (cm) | 30 | 50 | 12 | 55 | 30 | 20 | 35 | 40 | 38 | 45 |
| Seasons of the year in which it retained water | 1; 4 | 1;2;3;4 | 1; 4 | 1;2;3;4 | 1;2;3;4 | 1;2;4 | 1. | 1;2;3;4 | 1; 4 | 1; 4 |
| Number of months of the year in which it retained water | 6 | 12 | 4 | 12 | 10 | 6 | 7 | 7 | 6 | 6 |
| Aquatic vegetation (%) | >50% | >50% | <50 % | >50% | >50% | >50% | >50% | >50% | 50% | >50 % |
| Types of margin vegetation | Sh.;Und . | Sh.;Und . | Und. | Sh.;Und . | Sh.;Und . | Sh.;Und . | Sh.;Und . | Sh.;Und . | Sh.;Und . | Und. |

For each collection point the following local descriptors were evaluated: percentage of vegetation cover in the internal area of the cylinder, categorized in: 1) (none), 2) (1-25%), 3) (26 - 50%), 4) (51-75%), 5) (> 76%); distance from the sampling point to the margin (cm); depth of the sampling point (cm); pH; dissolved oxygen; water temperature (°C); month of sampling. PH and water temperature were measured with a pHmeter, and dissolved oxygen of water was measured with an oximeter. The depth of the water column was measured with a ruler positioned in the center of the cylinder. Lastly, we divided the area of the cylinder into four parts, each representing approximately 25% of the covered area, to measure the percentage of aquatic vegetation in each section. The variation of each environmental descriptor between the seasons is presented in the Supplementary Material (S4).

### ETHICS STATEMENT

Collection permits were provided by the Brazilian government (ICMBio) (authorization 40180-2). Field studies did not involve endangered or protected species. Manipulation of animals in the field was restricted to a minimum, as sampling was restricted to specimens collected in the points delimited by the cylinder (see the section above). The collected specimens were immediately anesthetized with xylocaine and fixed in 10% formaldehyde. This study was approved by the Research and Ethics committee of the Universidade Federal do Rio Grande do Sul.

### DATA MATRICES

We built two matrices – a presence/absence matrix and an environmental descriptor matrix – containing the data of the species collected at the collection points of each of the sampled ponds during the two seasons: when the water level in ponds is high, September and October/2013 and May-August/2014, and when the water level in ponds is low, November and December/2013 and January-April/2014 (see Supplementary Material Table S1 for details). Environmental variables were standardized by subtracting the corresponding average and dividing by the standard deviation.

Based on the information contained in the presence/absence matrix, we created a phylogenetic tree by pruning the dated amphibian tree proposed by Jetz and Pyron (2018) to include only the species found in the communities using the function *prune.sample* of the R package picante (Kembel et al., 2010). We then built a matrix of phylogenetic distances composed of all the species occurring in the ponds throughout the seasons.

For the functional matrix, we measured 12 different traits (see Table S5 of the Supplementary Material) from a minimum of five tadpoles of each species collected during the study period in all ponds. The tadpoles were previously classified according to developmental stage (*sensu* Gosner, 1960), and only those between stages 33 and 39 were measured (Queiroz, da Silva & Rossa-Feres, 2015; Jordani et al., 2019). Restriction to this range of development reduces the influence of intraspecific variation, thus excluding possible allometric differences related to ontogenetic development (Grosjean, 2005). The functional traits we used were selected on the basis of their well-known relations with feeding and swimming behavior, habitat use, or tadpole life history (Inger et al., 1987; Altig & Johnston, 1989; Rossa-Feres, Jim, & Fonseca, 2004; Lajmanovich, 2000; Eterovick & Barros, 2003; Strauβ et al., 2010).

We used Variance Inflation Factor Analysis (VIF, Lin, Foster, & Ungar, 2011) to assess multicollinearity between the descriptors of the environmental matrix and between the traits of the functional matrix. The value of VIF for an explanatory variable is obtained using the R^2^ value of the regression of this variable against all other explanatory variables. We considered variables with VIF values above five as correlated. Results were not indicative of significant correlation between any of the traits (Supplementary Material Table S6) or between any of the environmental descriptors (Supplementary Material Table S7).

All analyses were performed using the ‘SYNCSA’, ‘ade4’, ‘ape’ and ‘vegan’ packages in R (R Development Core Team, 2013).

### STATISTICAL ANALYSIS

The tadpole diversity at each of the sampling points was evaluated using Rao’s Quadratic Entropy Index (Rao, 1982). This index is based on the proportion of species present in a community and some measure of dissimilarity, ranging from 0 to 1. One of the advantages of this method is that it allows for the division of biodiversity into alpha, beta and gamma components (Pavoine, Dufour, & Chessel, 2004), providing a flexible framework that can be adapted to quantify and compare different components of diversity such as taxonomic, functional, and phylogenetic diversity of the communities (de Bello et al., 2009, Meynard et al., 2011, Bernard-Verdier, Flores, Navas & Garnier, 2013; Arnan, Cerdá & Retana, 2014; 2017). Taxonomic distances between species within each sampled point *k* were obtained from the presence/absence matrix using the formula:

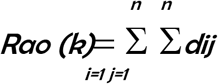

d*ij* = 1, where d*ij* is the distance between species *i* and *j*. Phylogenetic distances between species were measured using the co-phenetic distances of the phylogenetic matrix. We used the functional matrix to calculate the Euclidean distances between species based on the Gower distance. Finally, we evaluated the functional and phylogenetic redundancies that measure the resilience of a community to ensure the provision of ecosystem processes against any type of disturbance (Pillar et al., 2013). This maintenance is ensured by the presence of functional and phylogenetically similar species, differing in their responses to those disturbances. Redundancy metrics were obtained by calculating the difference between species diversity and the Rao quadratic entropy based on their functional and phylogenetic dissimilarity, respectively (de Bello et al., 2006). All Rao’s quadratic entropy calculations were performed using the ‘SYNCSA’ package in the R environment (R Development Core Team, 2013).

We used Linear Mixed-Effect Models (LMM) with Gaussian distribution to model the relation between diversity indexes (taxonomic, functional and phylogenetic diversity and functional and phylogenetic redundancies) and local environmental descriptors. This approach explicitly models the relation within the data set using random effects or latent random variables (Breslow & Clayton, 1993; Zhang et al., 2012). We built several models with different sets of environmental descriptors, so that all possible combinations would be evaluated. Pond was included as random effect. Model selection was carried out using the correted Akaike Information Criteria (AICc) to select the model containing the most information between all candidate hypotheses (Buham & Anderson, 2002). We also took into account the AICc weights (w), indicative of the empirical support for each model relative to the others in the candidate set. Finally, we applied a threshold of AICc 2 units to define model support (in other words, we considered models with ΔAIC<2 as equivalent; Zuur et al., 2009). Models were built using the ‘lme4’ package in R (R Development Core Team, 2013).

## RESULTS

We collected a total of 2,390 tadpoles from 18 species of three families: Hylidae (seven species), Leptodactylidae (10 species) and Odontoprhynidae (one species) (Table 2). The highest richness and abundance of tadpoles occurred when the water level was higher (Austral Spring and Winter) showed (Table 2).

**Table 2.**
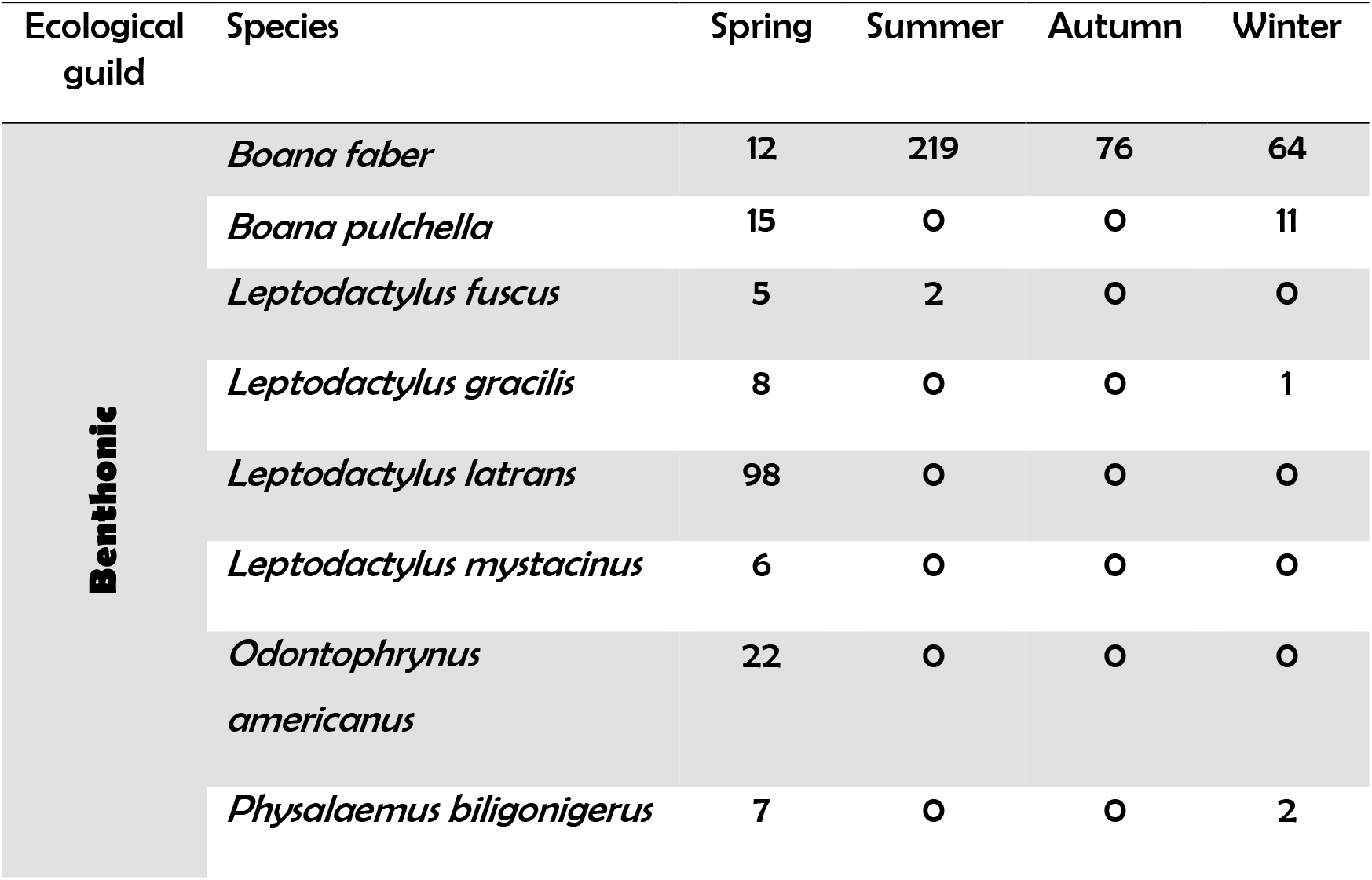
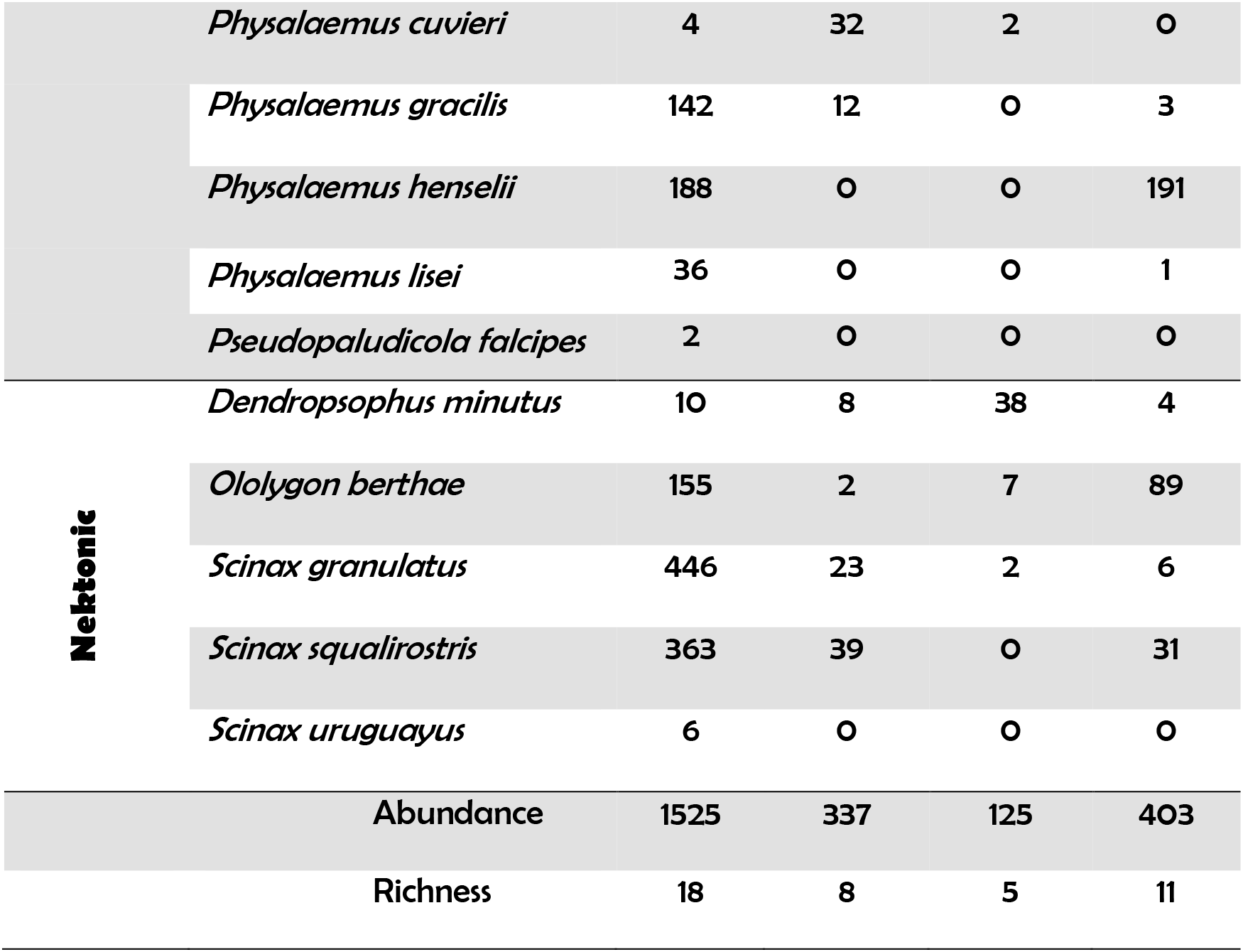
Abundance of tadpoles of anuran species registered at the Reserve Biológica do Lami, southern Brazil, between September/2013 and August/2014. Seasons with high water levels in ponds - Austral Spring and Winter; Seasons with low water levels in ponds Austral Summer and Autumn.

Models showed that the three components of diversity generally responded to the same set of local environmental descriptors (Tables 3 and 4). However, that relationship varied throughout the seasons. In the Austral Winter (when the water level is high) water depth positively affected the components of diversity. However, phylogenetic redundancy was not explained by any of the local environmental descriptors. When the water level is low – Austral Summer and Autumn – none of local environmental descriptors explained the observed patterns of diversity. The table including all the models for all the evaluated diversity metrics is presented as Supplementary Material (Tables S8-S27).

**Table 3.**
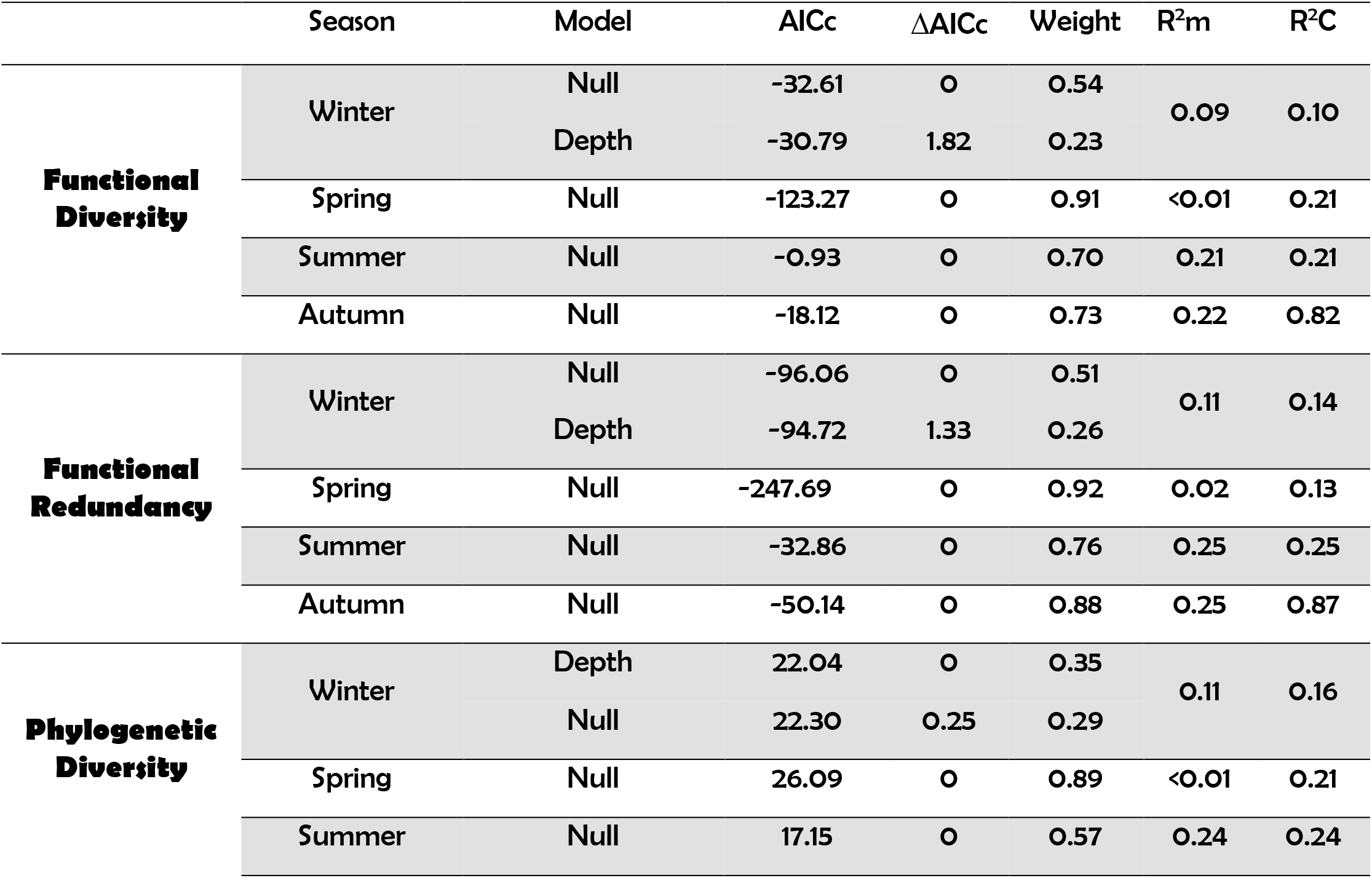
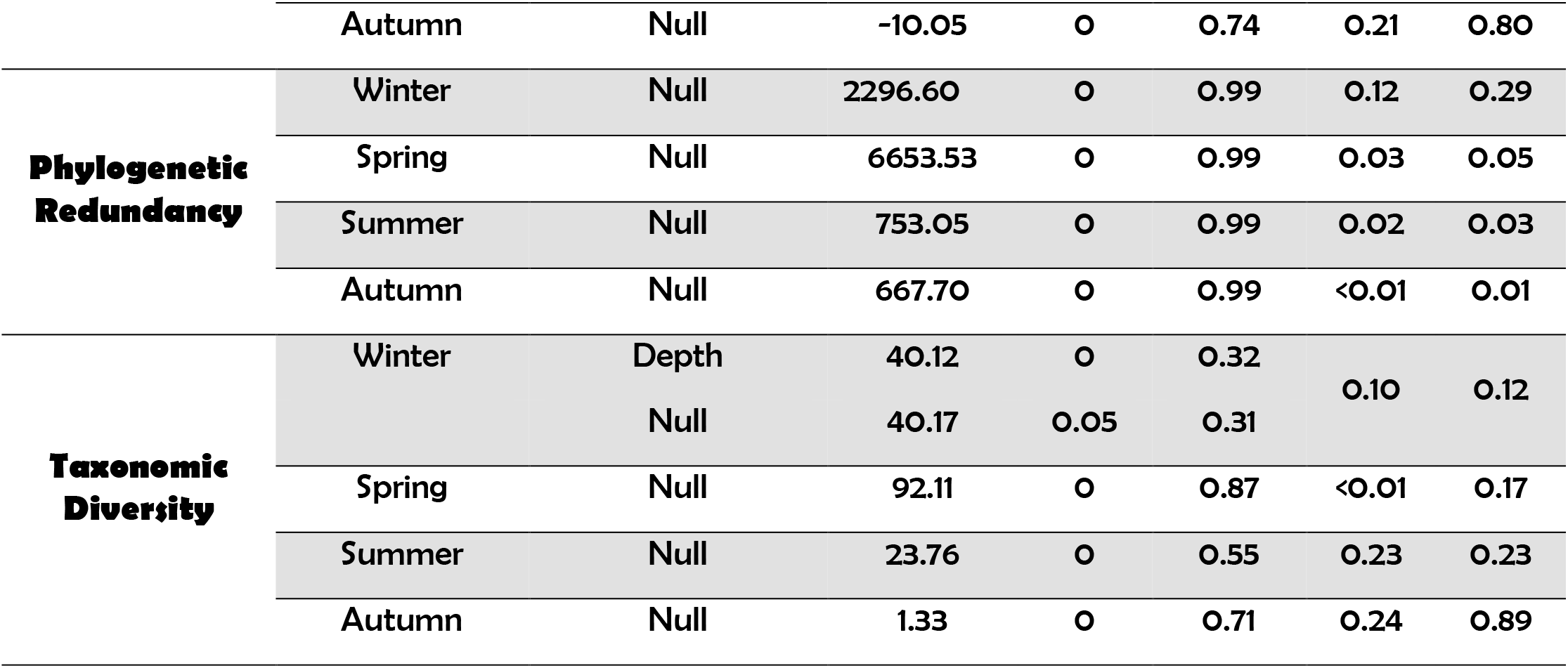
Summary of the best-adjusted LMM for each of the diversity components analyzed for tadpole communities. R^2^m (fixed effects); R^2^C (fixed + random effects).

**Table 4:** Results of ANOVA tests showing the effects of water depth on tadpole diversity components in the Winter.

| Diversity Components | Effect | F | p |
| --- | --- | --- | --- |
| Functional diversity | 0.05 | 6.95 | 0.010 |
| Functional redundancy | 0.04 | 9.06 | 0.003 |
| Phylogenetic diversity | 0.08 | 8.55 | 0.004 |
| Phylogenetic redundancy | NS | NS | NS |
| Taxonomic diversity | 0.09 | 8.08 | 0.006 |

During the Austral Winter, intermediary (16-45 centimeters) and deep (46-70 centimeters) water depth micro-habitats showed the highest richness and diversity of tadpoles (Figure 2; Figure 3 a-d), while shallow depth micro-habitats (0-15 cm) presented the lowest values. We also observed that benthic tadpoles – those with vertically flattened bodies and low fins – were more abundant at shallow and intermediary depths, whereas nektonic tadpoles – those with horizontally flattened bodies and high fins – were more abundant at intermediary and deep depths (Figure 4).

**Figure 2:**
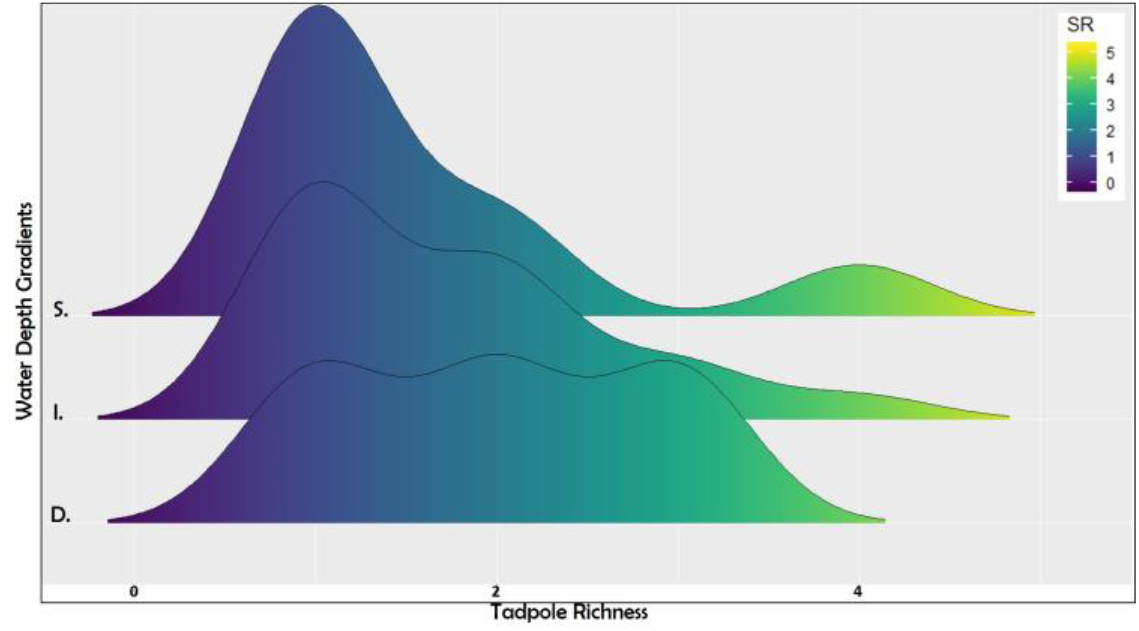
Tadpole richness distribution in microhabitats along the depth gradient during the winter. SR: species richness; S: shallow (0-15 cm); I: intermediary (16-45 cm); D: deep (46-70 cm).

**Figure 3:**
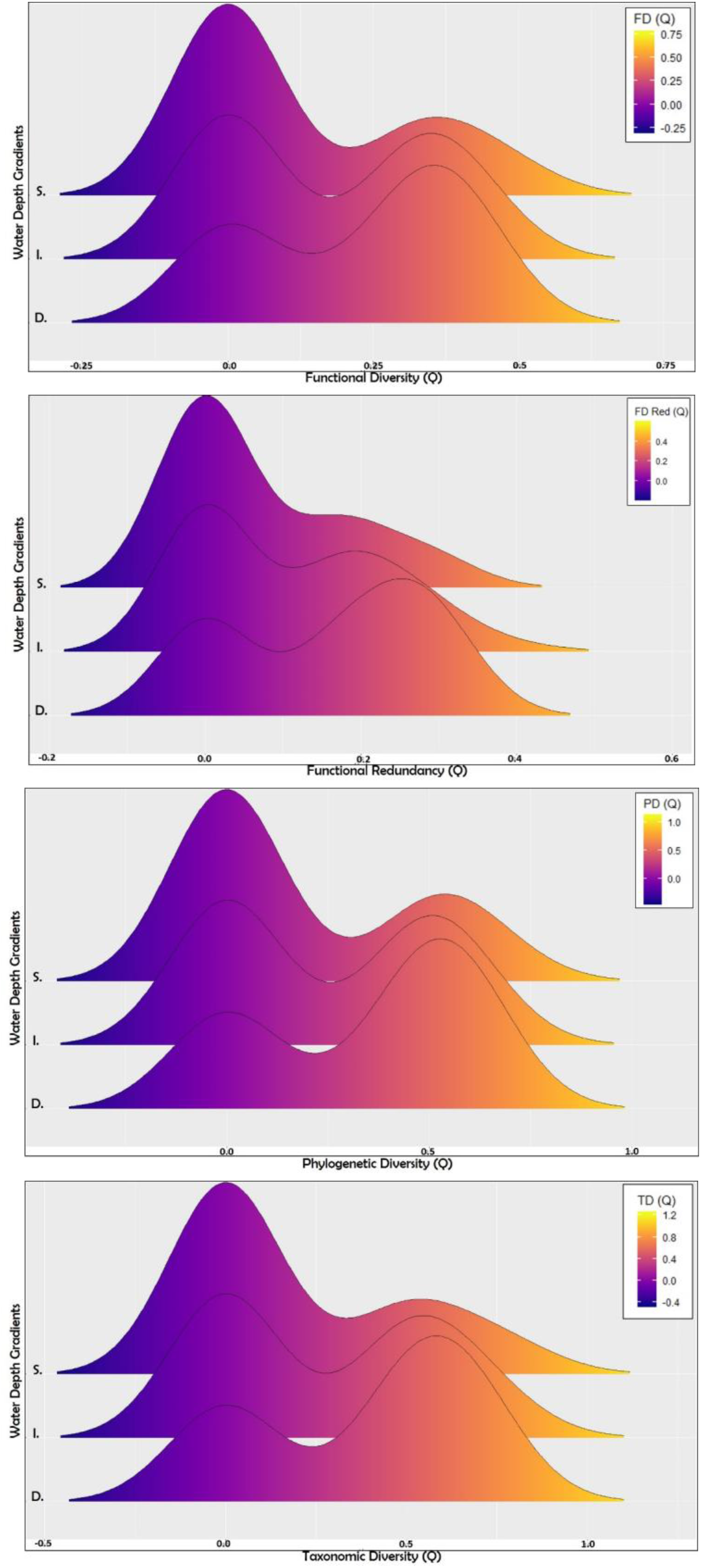
Distribution of the components of tadpole diversity along the depth gradient during the period of high water levels in ponds (winter). S: shallow (0-15 cm); I: intermediary (16-45 cm); D: deep (46-70 cm).

**Figure 4:**
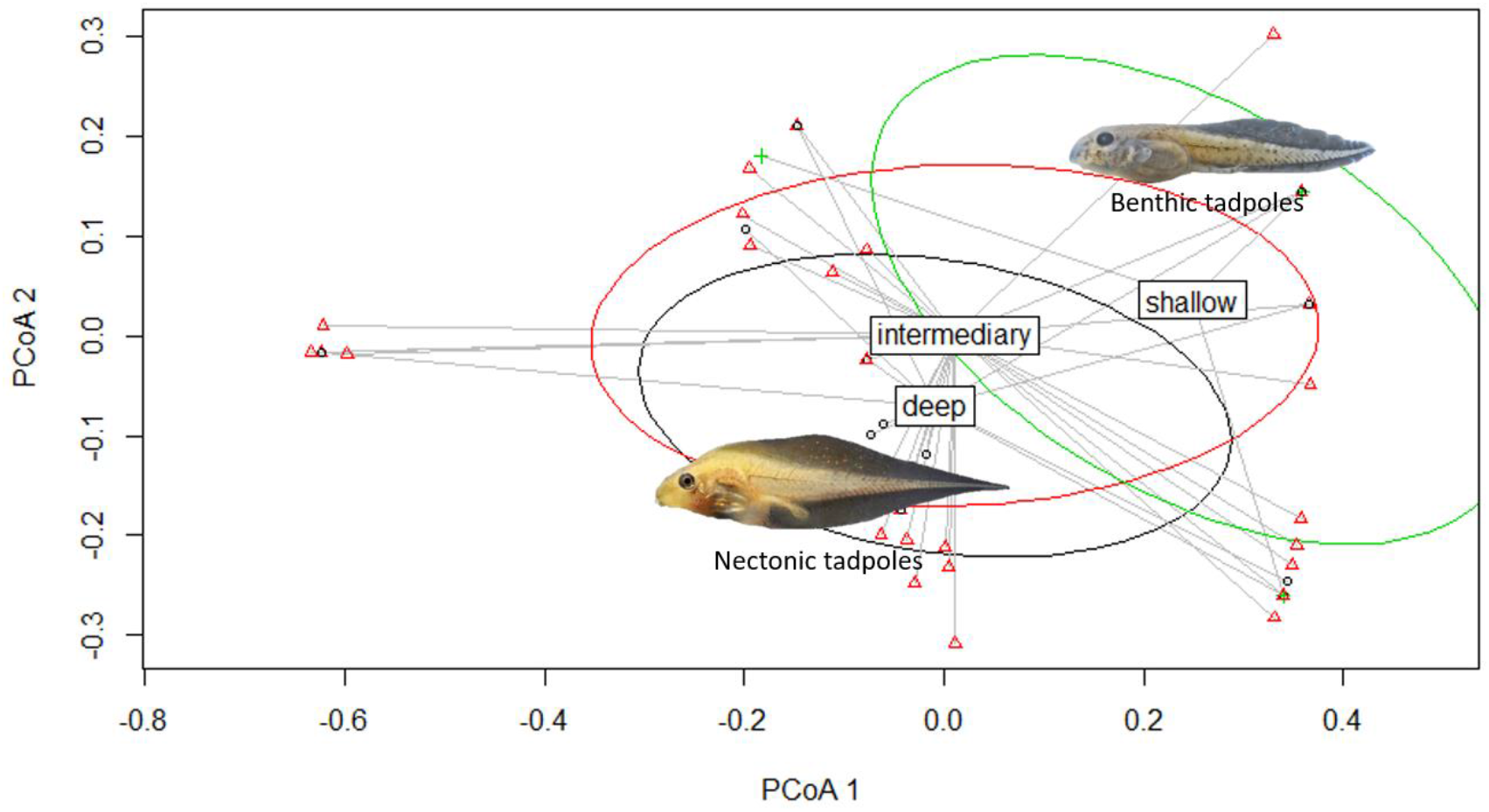
Distribution of tadpole morphological guilds along depth gradient during the period of high water levels in ponds (winter): shallow (0-15 cm); intermediary (16-45 cm); and deep (46-70 cm).

## DISCUSSION

### Tadpole community assembly is not completely random

Results showed local environmental factors influence tadpole diversity in ponds. Although the null model was frequently among the best set of models, all diversity metrics evaluated here were associated with water depth in the Austral Winter, when the water level is higher. Therefore, environmental descriptors drive the diversity of tadpoles in lentic systems, at least under such conditions. This is in agreement with non-random patterns of organization in tadpole communities found in other studies (Both et al., 2011b, Strauβ et al., 2016; Knauth, Moreira & Maltchik, 2018).

This not with standing, environmental filtering does not seem to be the only mechanism driving tadpole assembly. Recent studies have reported that tadpole diversity patterns in ponds are the result of a complex balance between environmental and spatial factors, the latter of which seemingly having the largest fraction of total variance observed in communities (Leão-Pires, Luiz & Sawaya, 2018; Marques et al., 2018; Dalmolin et al., 2019). Furthermore, other niche-based processes (e.g. predation and competition) have been identified as the main agents in the modification of the structure of aquatic communities, to the extent of modifying the relation between environmental and spatial factors and the components of diversity (Livingston et al., 2017).

The multifaceted approach we used made it possible to bring to the foreground the role of local environmental descriptors as predictors of tadpole diversity in temporary ponds. Some studies suggested that the environment governs the relation between taxonomic, functional and phylogenetic diversity and that these may covariate differentially along the environmental gradients (Devictor et al., 2010, Safi et al., 2011; Bernard-Verdier, Flores, Navas & Garnier et al., 2013). However, our results revealed congruent responses of the three components of diversity to the environmental gradients, corroborating our initial prediction, and patterns described for other anuran communities (Ribeiro et al., 2017, Escoriza & Ben Hassein, 2017).

### The relation between the environment and tadpole diversity changes seasonally

Results are consistent with the theory that environmental filtering is an important agent driving the assembly of communities in seasonal systems, as evidenced by several groups of aquatic organisms (e.g. Poff, 1997;; Florencio et al., 2014; Datry, Bonada & Heino, 2016), including anurans (Escoriza & Ben Hassine, 2017; Couto et al., 2017; Knauth, Moreira & Maltchik, 2018). The effect of local environmental factors on components of diversity was discrepant between seasons, and significantly when the water level is high. These results do not support our initial prediction and are discrepant with the findings of previous studies in similar systems, where there was evidence of environmental filtering when the water level is low (e.g. Chase, 2007; Ruhí et al., 2013).

Richness and abundance of tadpoles was highest in the end of the period when the water level is high and declined gradually towards periods when the water level is low, following the general pattern found in tropical and subtropical regions (e.g. Kopp, Wachlevski, & Eterovick, 2006, Both et al., 2008b; Both et al. 2011b; Strauβ et al., 2016). The period when water level is high coincides with the peak of the reproductive activity for most of the registered species (see Supplementary Material Table S28). During this period the ponds are full of water, and the gradual increase of photoperiod and air temperature may serve as stimulus for the beginning of reproduction of several species; this may even occur by the end of the Austral Winter (Both et al., 2008a). As environmental seasonality strongly influences anuran reproductive phenology and may affect functional and phylogenetic diversities (da Silva et al., 2012; Martins et al., 2015), it is not surprising that tadpole community assembly, specifically species spatial and temporal distribution, to be closely related to the reproductive activity of the adults (Resetaritis & Wilbur, 1991; Alford, 1999).

In the area of our study, the increase in water volume seems to create more favorable conditions for the species present (Lake, 2003), including greater diversity and availability of foraging and roosting resources (e.g. food and micro-habitats). Different micro-habitats combining for a wide array of environmental conditions are necessary for the co-occurrence of a larger set of species (Eterovick & Barata, 2006; Both et al., 2011b). Environmental heterogeneity (e.g. depth, temperature and vegetation) usually presents a linear relation with the diversity of tadpoles in ponds, as it maximizes the occupancy of the functional space and, consequently, should also add to the phylogenetic diversity (Escoriza & Ben Hassine, 2017).

Periods of low water were characterized by the reduction of tadpole richness and abundance. In fact, most tadpoles probably complete larval development and leave the ponds before these dry completely (Strauβ et al., 2016). During these periods, water depth may reach extremely low levels and this is a limiting factor for the occurrence of some groups of species that have limited swimming ability at shallow depths (e.g. genus *Scinax*; Queiroz, Silva & Rossa-Feres, 2015). Accordingly, the remaining sets of species were composed, for the most part, of species that present flat bodies, low fins and ventral oral discs (e.g. genus *Physalaemus*). These functional traits are shared by species belonging to different lineages; they constitute adaptations for exploring the bottom of the ponds and may also be important for resisting extreme temperatures and low levels of dissolved oxygen (Babbitt & Tarner, 2000; Martins et al., 2015; Queiroz, Silva & Rossa-Feres, 2015; Strauβ et al., 2016). Under these conditions the influence of local predictors may be irrelevant, and community patterns may simply result from stochasticity (Chase, 2007; Delatorre et al., 2015).

The interchange between the deterministic and stochastic processes in the assembly of tadpole communities along the disturbance cycle associated to the level of water corroborates the patterns evidenced for other organisms in ponds subjected to desiccation (Zhou et al., 2014; Llames et al., 2017; Daniel et al., 2019). These cycles influence the processes of ecological succession, and the disturbance intensity will condition communities to distinct structuring agents (Zhou et al., 2014; Måren et al., 2018). Although we have observed more severe local environmental conditions in seasons when the water volume in ponds is low, it is possible that environment filters the species in the late Spring, i.e. when evapotranspiration increases and local environmental conditions begin to modify. The end of the desiccation period resulting from the increase in water depth, occurring in the Winter, expose species to a new selection process, allowing for a modified community to establish and for species to segregate along depth gradients (Figure; O’Neill, 2016). In summary, the effect of deterministic processes on the ecological succession and on the community assembly seems to dominate in the early and late periods of the disturbance cycle, while stochastic processes apparently dominate in the intermediate periods (Zhou et al., 2014; Daniel et al., 2019).

### Tadpole diversity responds to water depth

When the water level in ponds is high, taxonomic, functional, and phylogenetic tadpole diversity seem to be influenced by water depth, corroborating our initial prediction. Many studies have reported a strong influence of water depth in the assembly process of aquatic organisms, including on functional and phylogenetic alpha and beta diversity patterns (e.g., corals: Doxa et al., 2016; fishes: Langer et al., 2017; anurans: Werner et al., 2007; Both et al., 2011a; Semlitsch et al., 2015; Queiroz, da Silva & Rossa-Feres, 2015; Péntek et al., 2016; Dalmolin et al., 2019) and also an influence on the recolonization of ponds that undergo some type of environmental stress (Lesbarrères et al., 2009).

Water depth is, obviously, related to the variation in the water volume of the ponds and may be used as proxy for the hydroperiod (Vanschoenwinkel et al., 2009). Tadpoles exhibit strong and direct responses to the variation in the water volume of the ponds (e.g. phenotypic plasticy and adjustments in the levels of activity, movement and time of metamorphosis), especially under extreme environmental situations (Welborn, Skelly & Werner, 1996; Werner et al., 2007). There seems to be a trend for peaks of richness, density and diversity to be reached at intermediate depths and hydroperiods (Welborn, Skelly & Werner, 1996; Snodgrass, Bryan Jr. & Burger, 2000; Semlitsch et al., 2015), and our results are consistent with this trend.

The relation between tadpole diversity and water depth became even clearer when traits were used to explain the patterns of functional dissimilarity between pond communities (Queiroz, da Silva & Rossa-Feres, 2015). Indeed, species seem to be filtered according to their swimming and foraging abilities (Escoriza & Ben Hassine, 2017), and ponds in the extreme of the depth gradient will mostly contain tadpoles of the Leptodactylidae (in shallow depths) or of the Hylidae (in deep depths), while intermediate depths should harbor more functionally and phylogenetically diverse clusters (Queiroz, da Silva & Rossa-Feres, 2015).

Water depth can also affect the resources used by tadpoles. In fact, pond productivity, represented by abundance and diversity of aquatic vegetation, algae (which are food resources for tadpoles), and other organic matter, are all influenced by depth (Altig, 2007; Shulse et al., 2010, 2012; Langer et al., 2017). Foraging and roosting resources decline in extreme ranges of the depth gradient and are less variable in seasons when the water persists for longer times in the ponds (Wetzel, 2001; Anusa, Ndagurwa & Magadza, 2012; Doxa et al., 2016; Langer et al., 2017). So, the availability and variation of these resources should drive diversity patterns and promote dissimilarities at the local scale (Levin et al., 2001; Lins et al., 2017).

## CONCLUSIONS

Tadpole components of diversity respond to environmental factors. However, in our study system this relation was only significant when the water level in ponds is high, because the environmental conditions are milder and tend to allow for the coexistence of phylogenetically and functionally diverse taxa. Under these conditions diversity metrics were significantly associated with water depth. Also, the diversity metrics used here were consistent in their response to local environmental factors, demonstrating that the results produced for one metric may be extended to the others. Our results show the impact of environment on the anuran life cycle and reinforce the thesis that the assembly of tadpole communities is not random. Our results contribute to the understanding of the ecology of the anuran larval phase, a less known stage of the life of a group of organisms participating in important and diverse ecosystem functions.

## ACKNOWLEDGMENTS

This study was done under licenses from the ICMBIO (Ministério do Meio Ambiente, Brazil) and the Research and Ethics committee of the Universidade Federal do Rio Grande do Sul (UFRGS). We thank the Conselho Nacional de Desenvolvimento Científico e Tecnológico (CNPq) for scholarships and grants provided to DAD, TGS and MJRP. We thank the Coordenação de Aperfeiçoamento de Pessoal de Nível Superior (CAPES) for the financial support to the Post-Graduation Program in Animal Biology - UFRGS. The continuity and universality of the Brazilian research grant system is fundamental for the country’s scientific development, and we vehemently oppose to the political attacks perpetrated by the current federal government to public education, research and biodiversity conservation in Brazil. We are grateful to the staff of the Reserva Biológica do Lami for the support during the field activities.. We also thank all the colleagues who helped during field and lab work, and Márcio Borges Martins (UFRGS), Leandro Duarte (UFRGS), Luis Maurício Bini (Universidade Federal de Goiás), Eduardo Ferreira (Universidade de Aveiro), Vinícius Bastazini (Centre National de la Recherche Scientifique), and two anonymous reviewers for their comments on previous versions of the manuscript.

## SUPPLEMENTARY MATERIAL

### 1) Supplementary Material relating to the material and methods section

**S1.** Monthly average of the climate descriptors recorded for the city of Porto Alegre, Rio Grande do Sul, Brazil, in the period between September 2013 and August 2014.

| Month | Season | Photoperiod/day length (hours) | Total Precipitation (cm<sup>3</sup>) | Maximum Temperature (°C) | Minimum Temperature (°C) | Air humidity (average) | Evaporation (average) |
| --- | --- | --- | --- | --- | --- | --- | --- |
| September/2013 | Wet | 137,7 | 135,3 | 21,2 | 13,6 | 78,3 | 55,9 |
| October/2013 | Wet | 207,8 | 131,8 | 22,9 | 15 | 74,7 | 75,3 |
| November/2013 | Dry | 258,1 | 184,6 | 26,5 | 17,47 | 71,1 | 107,1 |
| December/2013 | Dry | 282,8 | 76,6 | 33,1 | 20,28 | 70,9 | 123,9 |
| January/2014 | Dry | 244,9 | 76,8 | 33,8 | 21,9 | 71,2 | 106,7 |
| February/2014 | Dry | 222,3 | 147,7 | 32,4 | 21,8 | 74,3 | 130,5 |
| March/2014 | Dry | 191,3 | 126,4 | 29,8 | 19,41 | 77,7 | 106,9 |
| April/2014 | Dry | 181,4 | 83,5 | 27,4 | 17,4 | 76,9 | 78,8 |
| May/2014 | Wet | 134,7 | 71 | 21,6 | 13,67 | 83,8 | 47,1 |
| June/2014 | Wet | 102,8 | 228,6 | 19,8 | 12,11 | 87,1 | 36,1 |
| July/2014 | Wet | 154 | 253,5 | 21,16 | 11,29 | 82,8 | 38,3 |
| August/2014 | Wet | 164,3 | 129,1 | 22,8 | 10,5 | 78,1 | 43,6 |

**S2.**
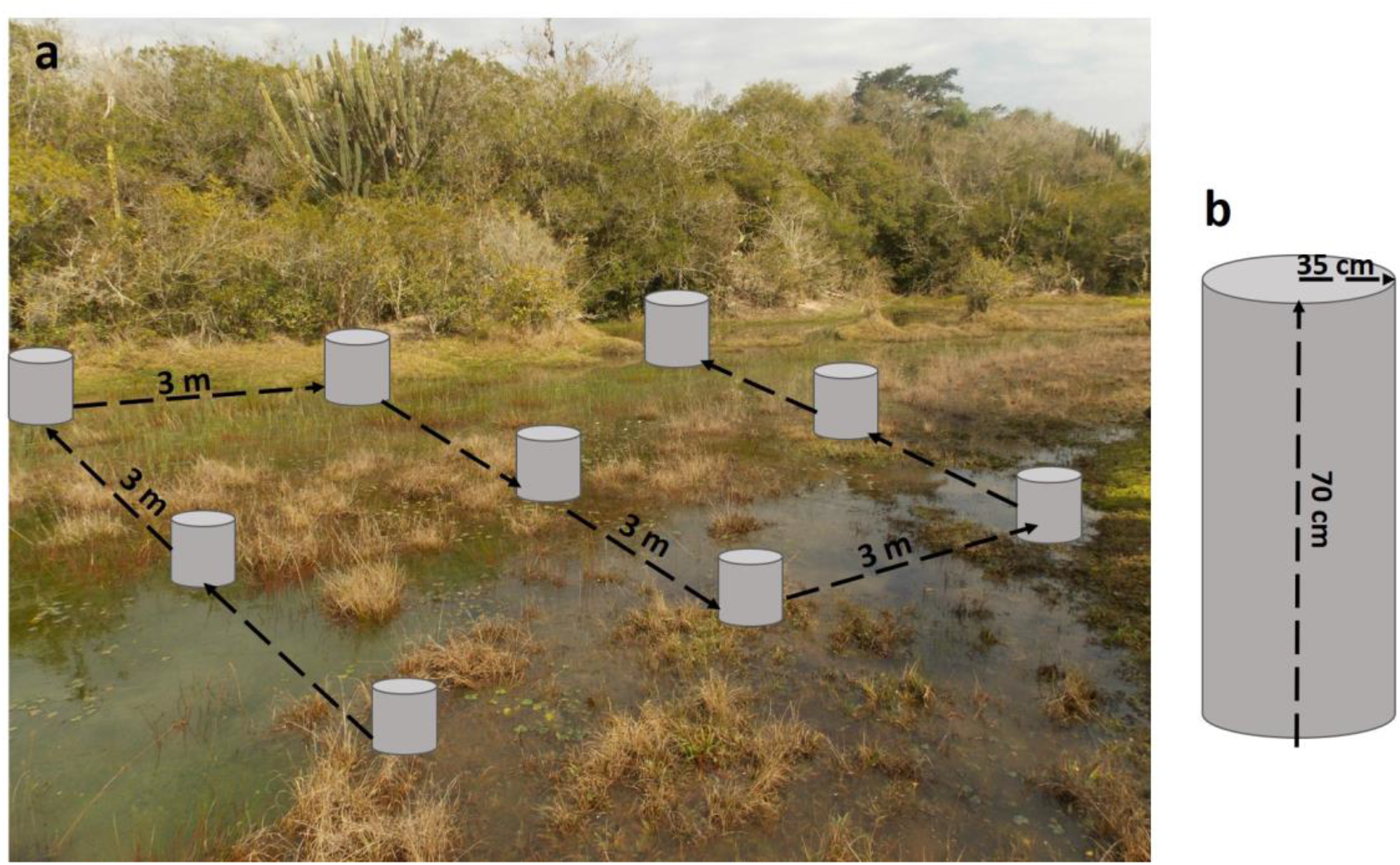
Scheme showing (a) the sampling procedure in transects and (b) the metal cylinder that was used to delimit the collection points and confine the tadpoles.

**S3.** Number of points sampled in each pond across seasons

| Pond | Spring | Summer | Autumn | Winter |
| --- | --- | --- | --- | --- |
| Pond 1 | 12 | 0 | 0 | 3 |
| Pond 2 | 49 | 17 | 9 | 17 |
| Pond 3 | 10 | 0 | 0 | 6 |
| Pond 4 | 41 | 7 | 16 | 9 |
| Pond 5 | 45 | 8 | 5 | 21 |
| Pond 6 | 23 | 4 | 0 | 7 |
| Pond 7 | 6 | 0 | 0 | 0 |
| Pond 8 | 29 | 6 | 3 | 14 |
| Pond 9 | 9 | 0 | 0 | 6 |
| Pond 10 | 11 | 0 | 0 | 8 |
| **Total** | 235 | 42 | 33 | 91 |

**S4.**
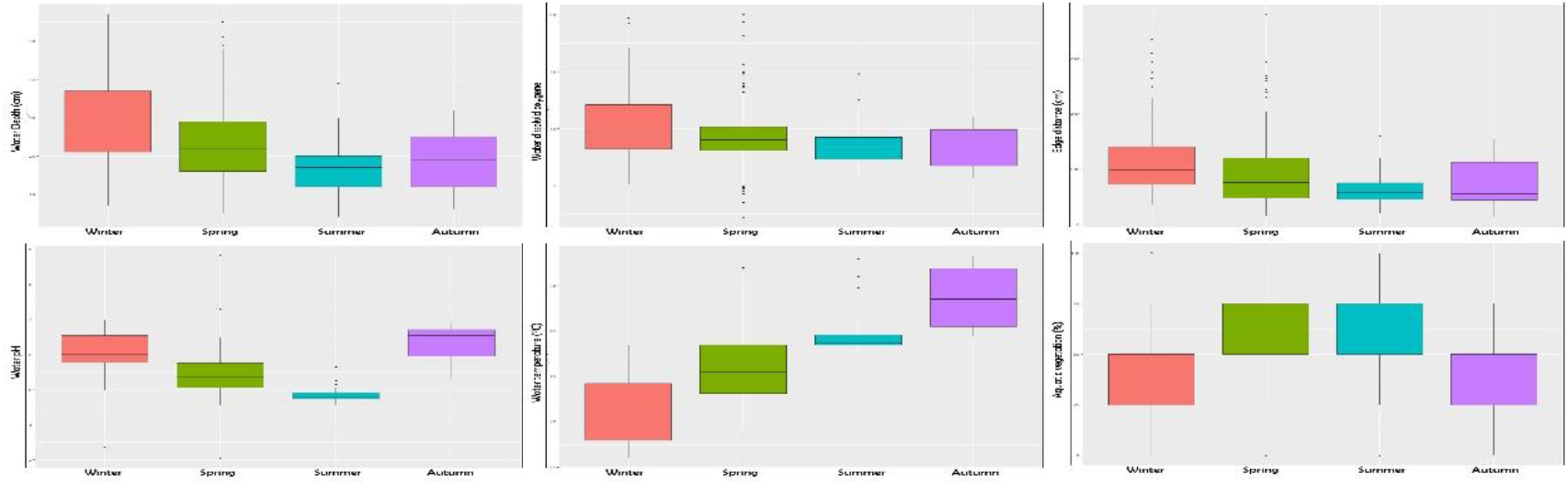
Boxplots showing the total variation in micro-habitats within each season of: a) water depth; b) dissolved oxygen; c) of distance from micro-habitats to the ponds’ edge; d) pH of water; e) temperature of water; f) % aquatic vegetation.

**S5.** Functional traits measured from different functional characteristics of tadpoles present in the studied ponds.

| Functional trait | Characteristics measured | Variable |
| --- | --- | --- |
| Body Height | body height/total length | Continuous |
| Body length | Body length/total length | Continuous |
| Body Width | Body Width/Total Length | Continuous |
| Caudal muscles width | maximum width of the caudal musculature/body length | Continuous |
| Dorsal fin height | maximum height of dorsal fin/ maximum height of the caudal musculature | Continuous |
| Ventral fin height | maximum height of ventral fin/ maximum height of the caudal musculature | Continuous |
| Number of teeth rows | Sum of number of forward and back rows | Continuous |
| Eye size | eye diameter / body length | Continuous |
| Oral disc position | Anteroventral; ventral | Categorical |
| Position in the water column | Benthic; nectonic | Categorical |
| Protection of eggs and early stages of life | protected in a foam nest; protected in natural or constructed basins; none/absent | Categorical |
| Presence of scourge | Present or absent | Binary |

**S6.** Variation Inflation Factor Analysis of tadpole traits.

36 – Variation Inflation Factor Analysis of tadpole traits.
| Functional traits | VIF value |
| --- | --- |
| Body length | 1.62 |
| Body width | 2.06 |
| Body height | 1.93 |
| Width of caudal muscles | 1.45 |
| Dorsal fin height | 1.52 |
| Ventral fin height | 1.22 |
| Nostril size | 1.03 |
| Nostril position | 2.06 |
| Eye size | 1.49 |
| Spiracle length | 1.34 |
| Opening position of the spiracle | 1.29 |
| Presence of flagellum | 1.17 |
| Number of teeth rows | 1.05 |
| Oral disc position | 2.76 |
| Eye position | 1.47 |
| Eggs | 1.21 |

**§7** – Variation Inflation Factor Analysis of Environmental descriptors of micro-habitats.

**S7.** Variation Inflation Factor Analysis of Environmental descriptors of micro-habitats.

| Environmental descriptors | VIF value |
| --- | --- |
| Aquatic Vegetation | 1.08 |
| Depth | 1.32 |
| Dissolved Oxygen | 1.02 |
| Edge Distance | 1.13 |
| pH | 1.10 |
| Temperature | 1.21 |

### 2) Supplementary Material relating to Results section

**S8:**
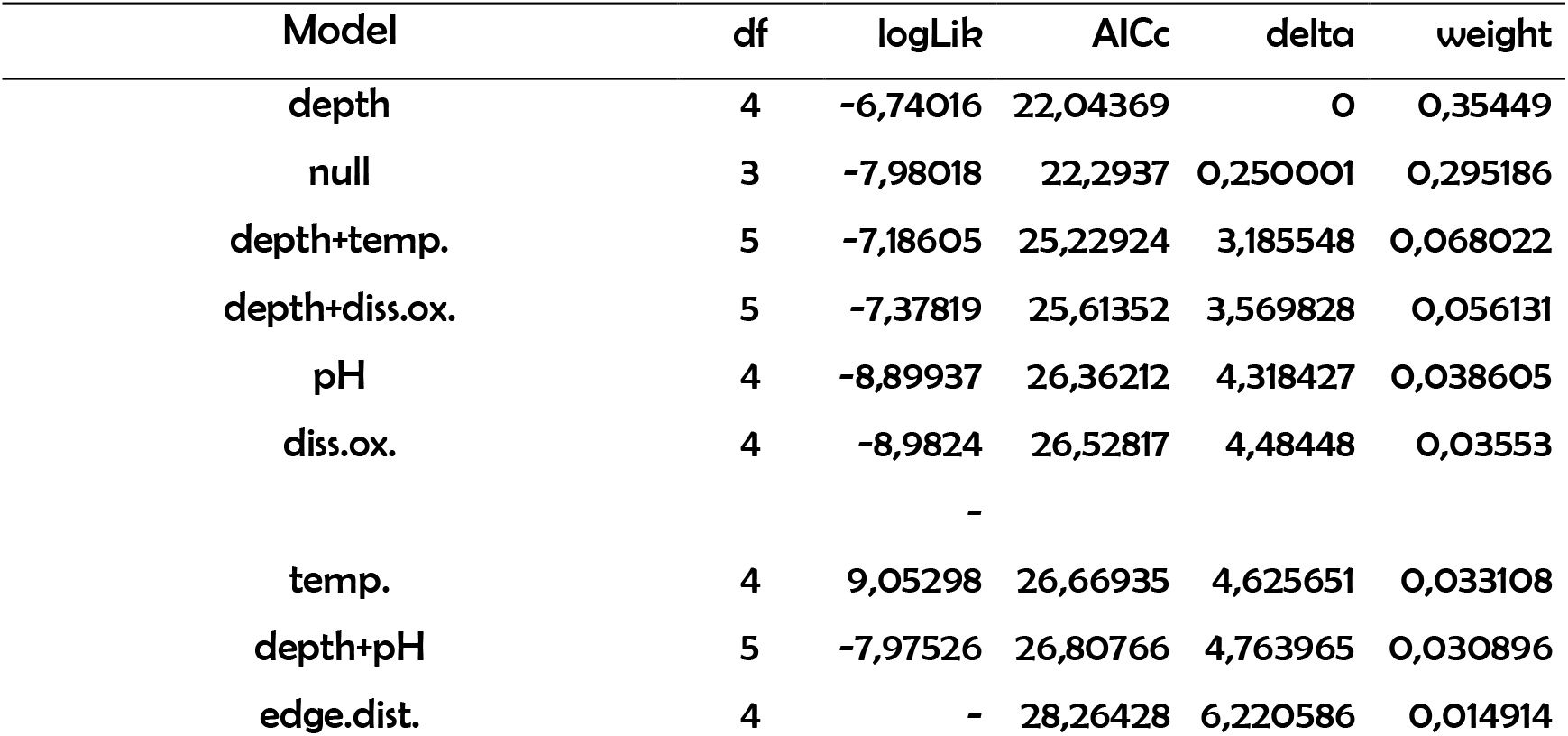
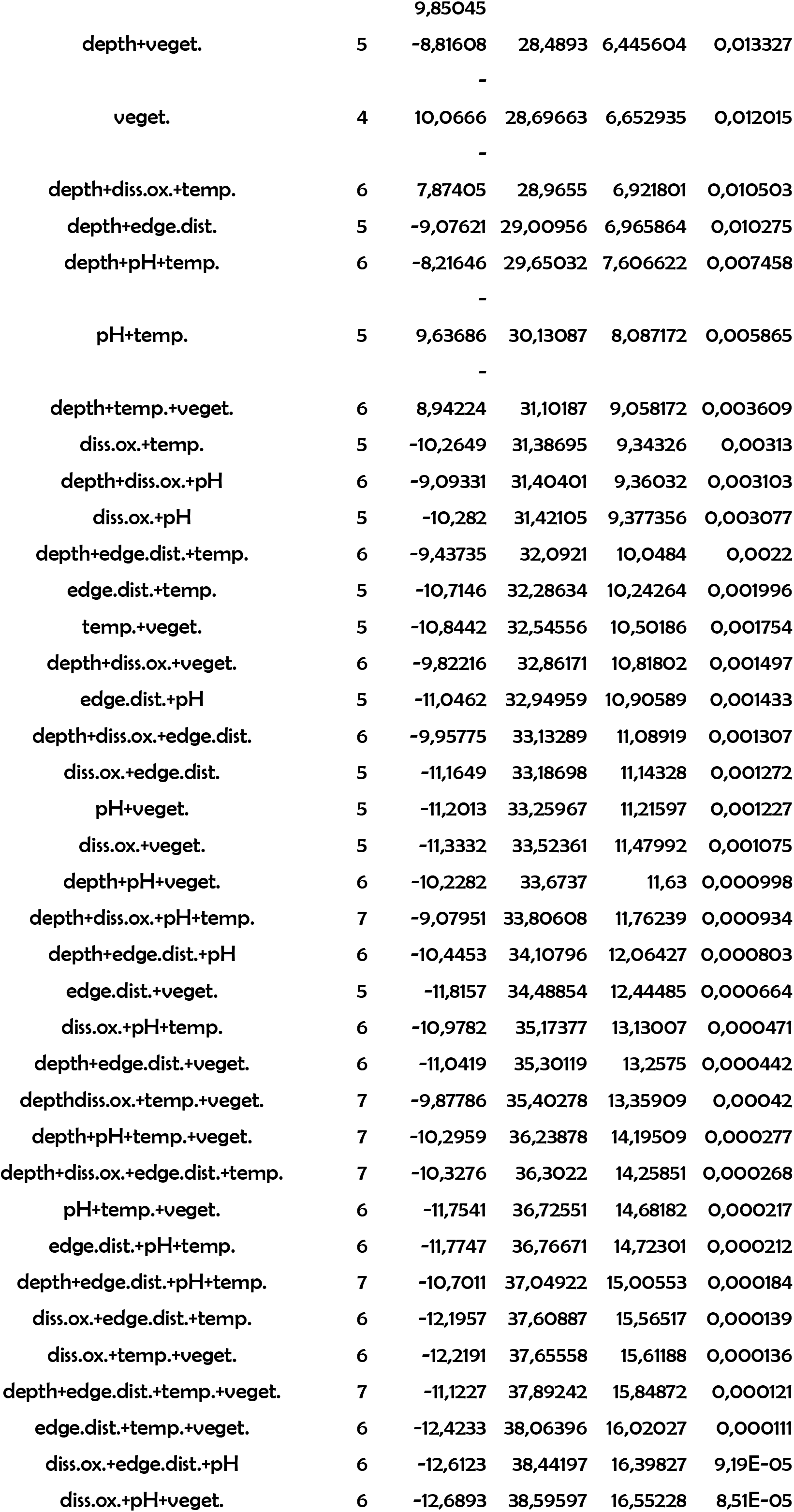
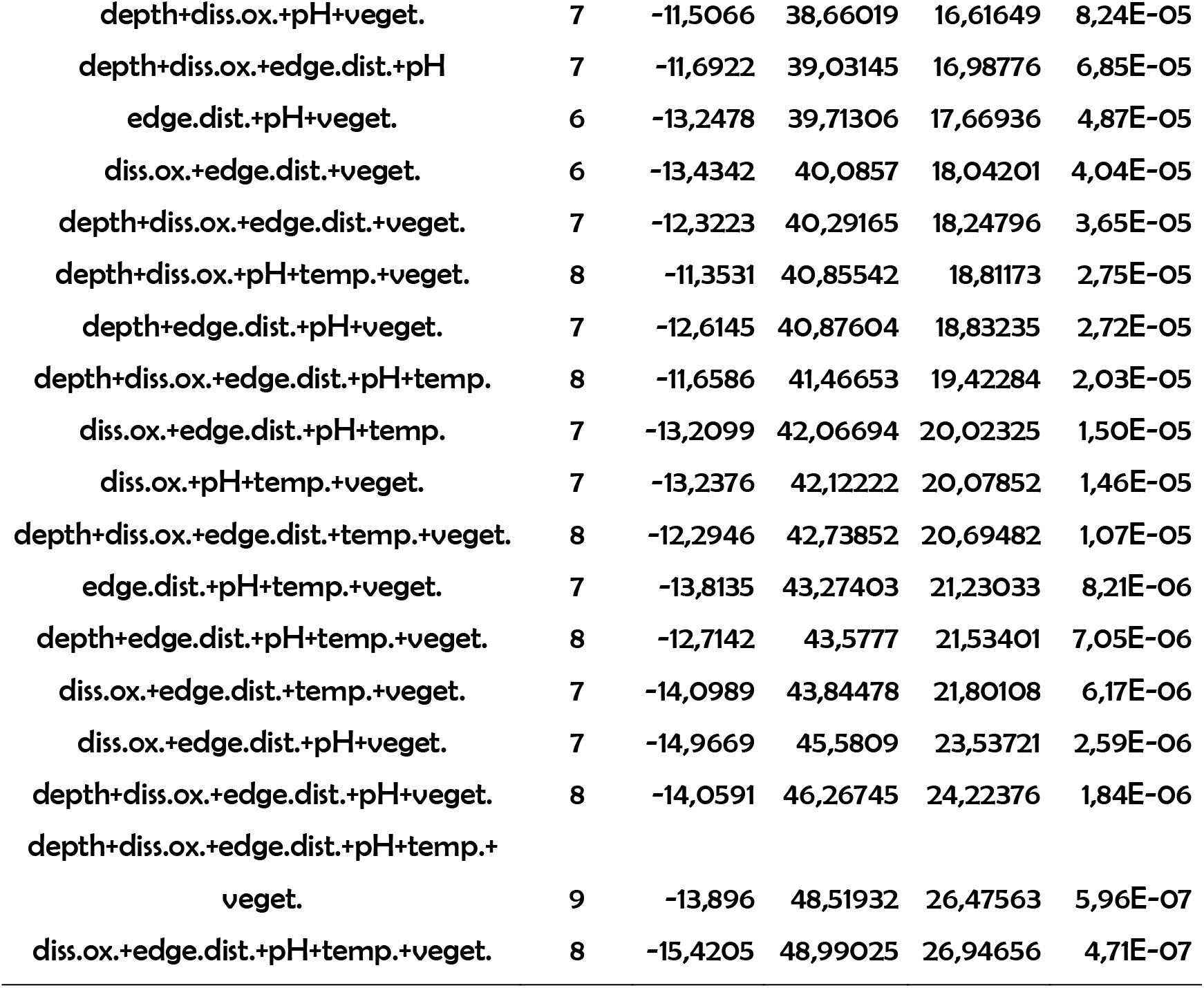
Summary of adjusted models produced by GLMM analysis relating local descriptors and phylogenetic diversity in tadpole communities in ponds at Winter season (season with low hydric stress). Local descriptors: Depth (water column depth); Diss. ox. (dissolved oxygen); edge. dist. (edge distance); temp (water temperature); veget (percentage of aquatic vegetation cover).

**S9:**
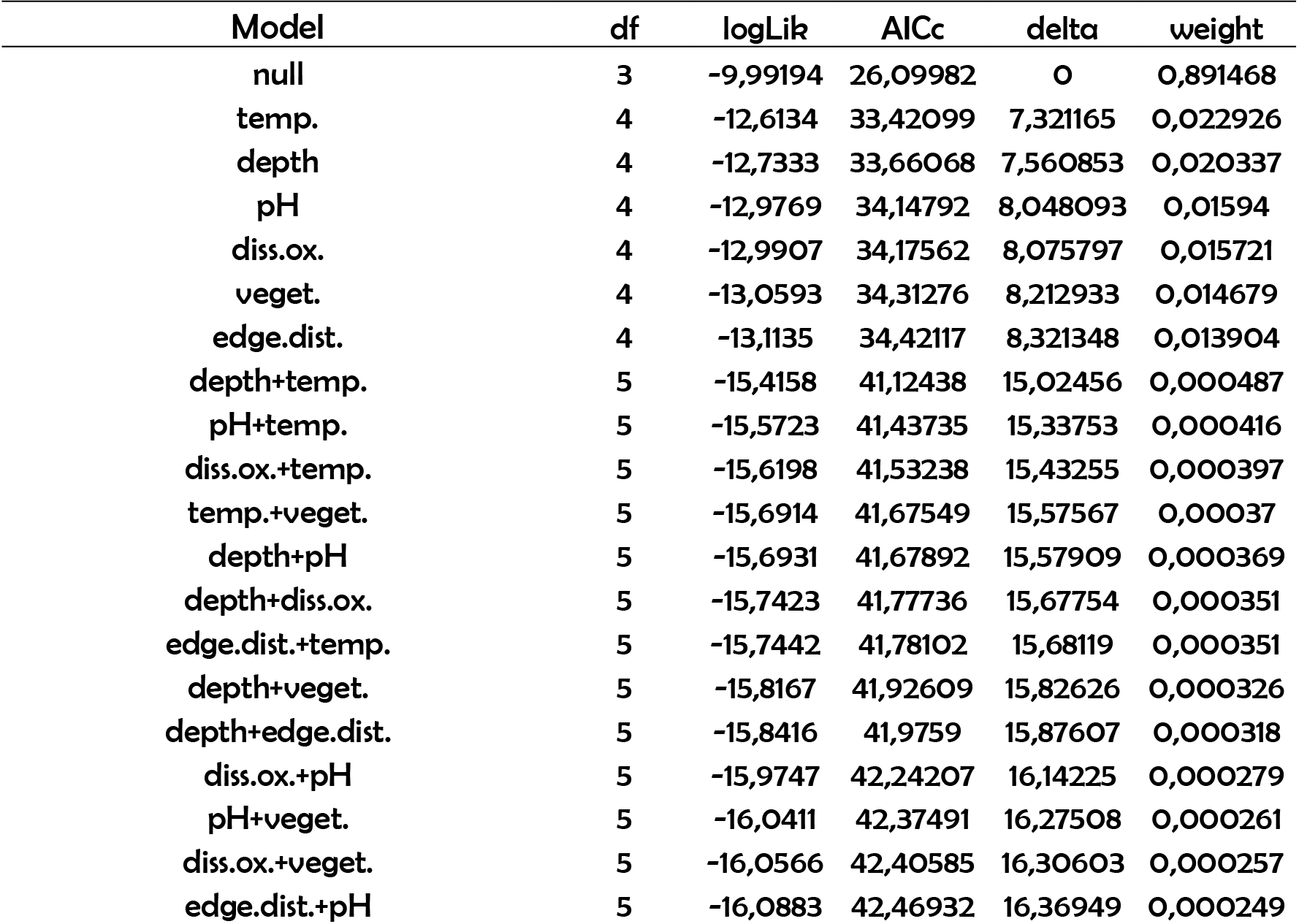
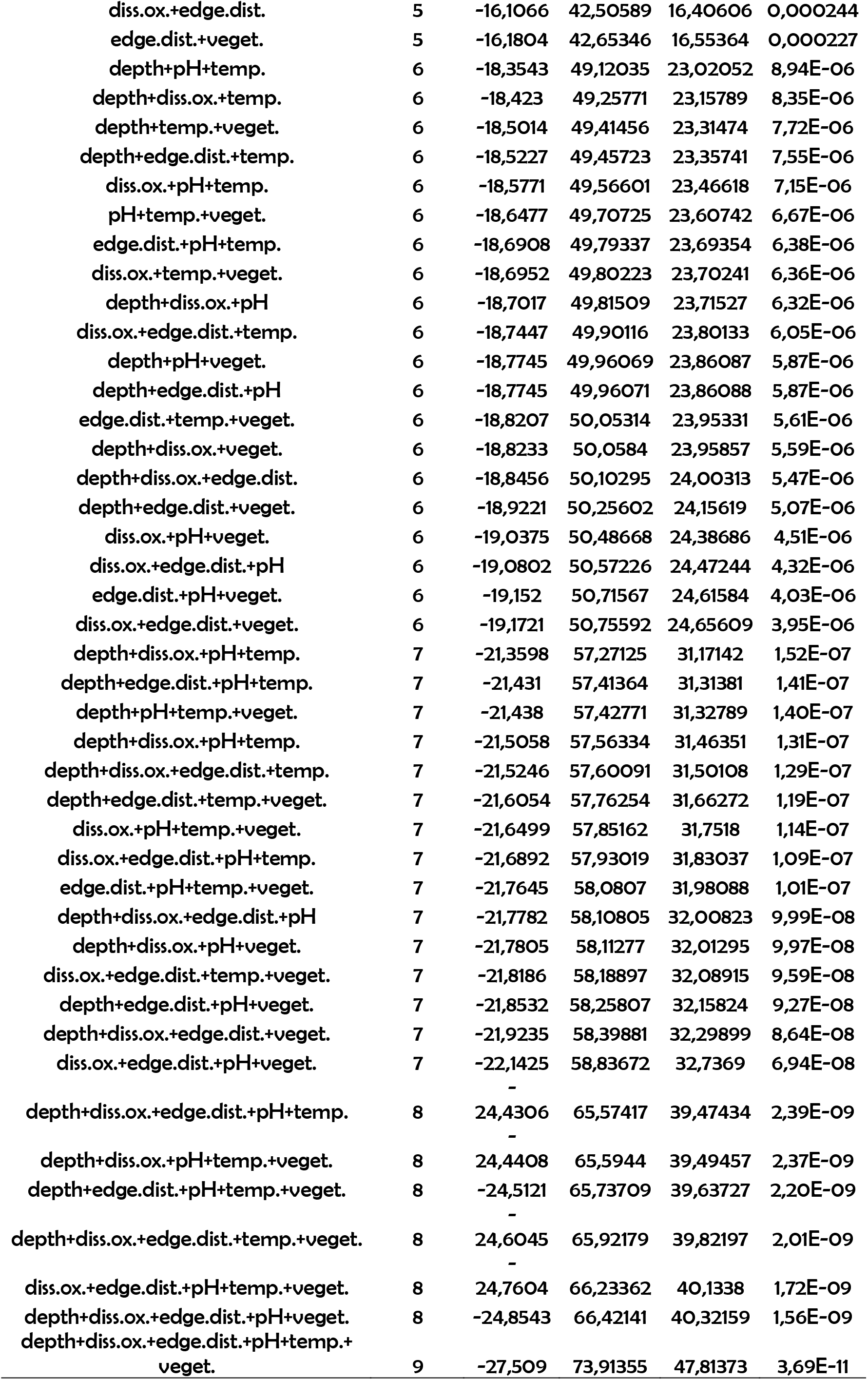
Summary of adjusted models produced by GLMM analysis relating local descriptors and phylogenetic diversity in tadpole communities in ponds in the Spring (season with low hydric stress). Local descriptors: Depth (water column depth); Diss.ox. (dissolved oxygen); edge.dist. (edge distance); temp (water temperature); <u>veget</u> (percentage of aquatic vegetation cover).

**S10:**
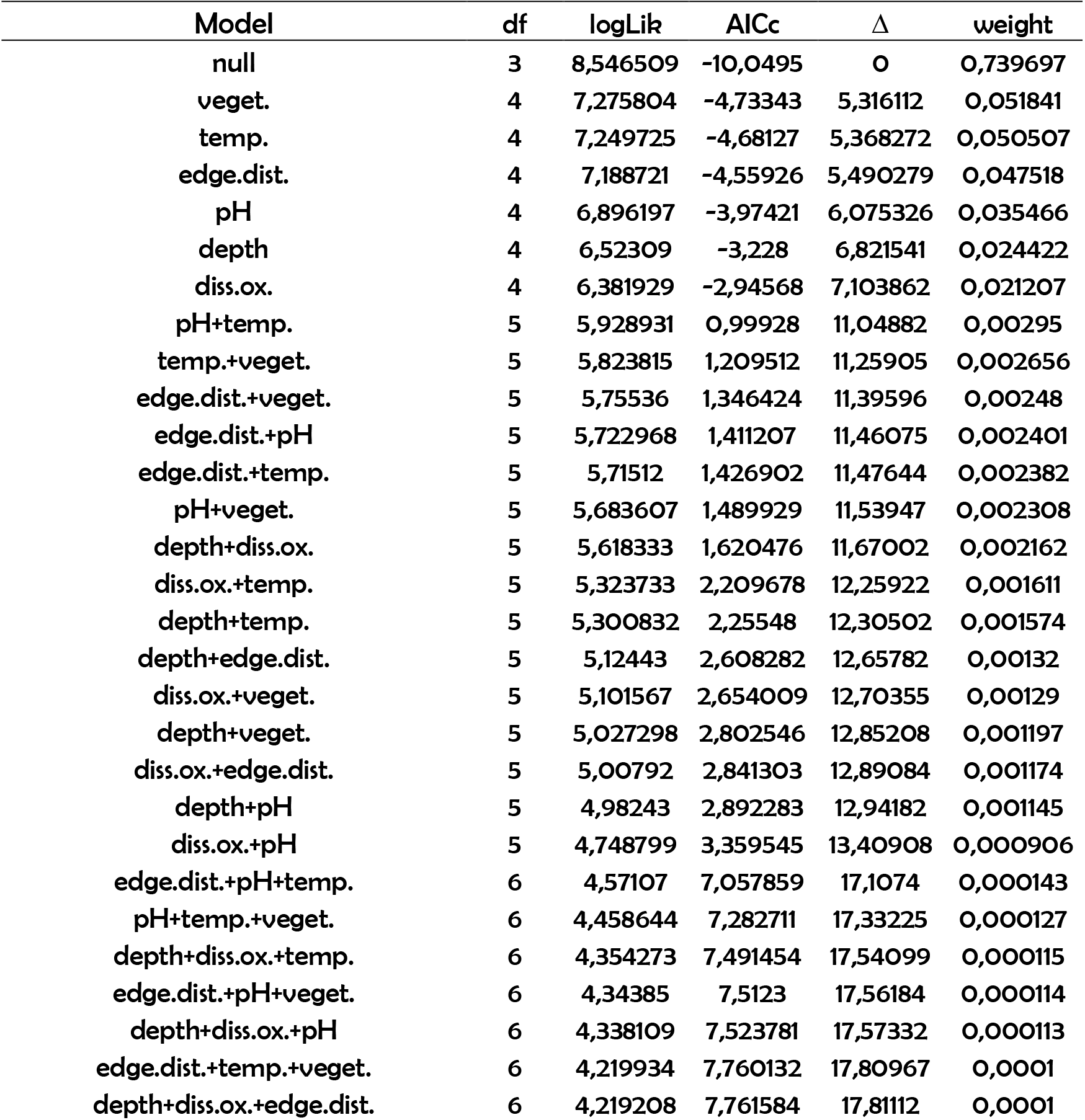
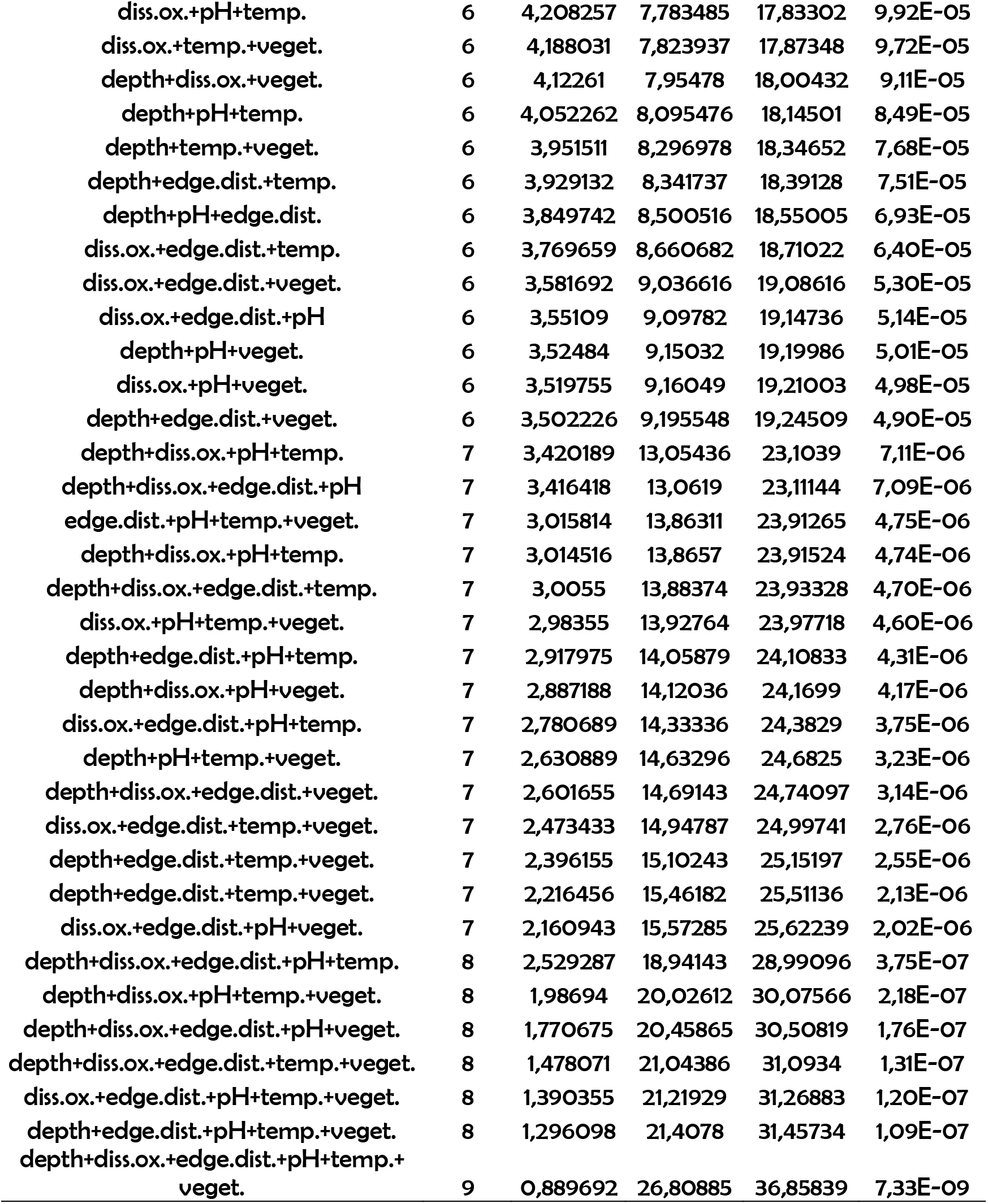
Summary of adjusted models produced by GLMM analysis relating local descriptors and phylogenetic diversity in tadpole communities in ponds in the Autumn (season with high hydric stress). Local descriptors: Depth (water column depth); Diss.ox. (dissolved oxygen); edge.dist. (edge distance); temp (water temperature); veget (percentage of aquatic vegetation cover).

**S11:** Summary of adjusted models produced by GLMM analysis relating local descriptors and phylogenetic redundancy in tadpole communities in ponds in the Winter (season with low hydric stress). Local descriptors: Depth (water column depth); Diss.ox. (dissolved oxygen); edge.dist. (edge distance); temp (water temperature); veget (percentage of aquatic vegetation cover).

| Model | df | logLik | AICc | $\Delta$ | weight |
| --- | --- | --- | --- | --- | --- |
| null | 9 | -1137,94 | 2296,598 | 0 | 0,999988 |
| depth+diss.ox.+edge.dist.+pH+temp.+veget. | 8 | -1151,79 | 2321,725 | 25,12764 | 3,50E-06 |
| depth+diss.ox.+pH+temp.+veget. | 8 | -1151,82 | 2321,789 | 25,19087 | 3,39E-06 |
| depth+diss.ox.+edge.dist.+temp.+veget. | 8 | -1151,95 | 2322,049 | 25,45131 | 2,97E-06 |
| depth+diss.ox.+edge.dist.+pH+temp. | 8 | -1152,65 | 2323,449 | 26,85073 | 1,48E-06 |
| diss.ox.+edge.dist.+pH+temp.+veget. | 8 | -1153,52 | 2325,193 | 28,59477 | 6,18E-07 |
| depth+edge.dist.+pH+temp.+veget. | 8 | -1156,32 | 2330,791 | 34,19341 | 3,76E-08 |
| depth+diss.ox.+edge.dist.+pH+veget. | 7 | -1165,68 | 2347,016 | 50,41842 | 1,13E-11 |
| depth+diss.ox.+temp.+veget. | 7 | -1165,8 | 2347,244 | 50,64595 | 1,01E-11 |
| depth+diss.ox.+edge.dist.+temp. | 7 | -1165,9 | 2347,438 | 50,84069 | 9,12E-12 |
| depth+diss.ox.+pH+temp. | 7 | -1166,36 | 2348,36 | 51,76227 | 5,75E-12 |
| diss.ox.+pH+temp.+veget. | 7 | -1166,54 | 2348,724 | 52,12617 | 4,80E-12 |
| diss.ox.+edge.dist.+temp.+veget. | 7 | -1166,63 | 2348,911 | 52,31293 | 4,37E-12 |
| diss.ox.+edge.dist.+pH+temp. | 7 | -1167,25 | 2350,148 | 53,55027 | 2,35E-12 |
| depth+pH+temp.+veget. | 7 | -1167,43 | 2350,497 | 53,89961 | 1,98E-12 |
| depth+edge.dist.+temp.+veget. | 7 | -1167,78 | 2351,209 | 54,61116 | 1,38E-12 |
| depth+edge.dist.+pH+temp. | 7 | -1168,02 | 2351,692 | 55,09424 | 1,09E-12 |
| edge.dist.+pH+temp.+veget. | 7 | -1170,12 | 2355,894 | 59,29589 | 1,33E-13 |
| depth+diss.ox.+pH+veget. | 7 | -1170,25 | 2356,143 | 59,54534 | 1,17E-13 |
| depth+diss.ox.+edge.dist.+veget. | 7 | -1170,36 | 2356,376 | 59,77868 | 1,05E-13 |
| depth+diss.ox.+edge.dist.+pH | 7 | -1170,74 | 2357,133 | 60,53482 | 7,16E-14 |
| diss.ox.+edge.dist.+pH+veget. | 7 | -1171,63 | 2358,898 | 62,30019 | 2,96E-14 |
| depth+edge.dist.+temp.+veget. | 6 | -1179,73 | 2372,685 | 76,08765 | 3,00E-17 |
| depth+diss.ox.+temp. | 6 | -1180,23 | 2373,677 | 77,07952 | 1,83E-17 |
| diss.ox.+temp.+veget. | 6 | -1180,42 | 2374,062 | 77,46412 | 1,51E-17 |
| diss.ox.+pH+temp. | 6 | -1180,51 | 2374,246 | 77,64868 | 1,38E-17 |
| diss.ox.+edge.dist.+temp. | 6 | -1181,15 | 2375,509 | 78,91092 | 7,32E-18 |
| depth+temp.+veget. | 6 | -1181,59 | 2376,39 | 79,79265 | 4,71E-18 |
| depth+pH+temp. | 6 | -1181,65 | 2376,52 | 79,92255 | 4,42E-18 |
| pH+temp.+veget. | 6 | -1181,68 | 2376,586 | 79,98785 | 4,27E-18 |
| depth+edge.dist.+temp. | 6 | -1181,93 | 2377,084 | 80,48637 | 3,33E-18 |
| edge.dist.+temp.+veget. | 6 | -1182,25 | 2377,713 | 81,11551 | 2,43E-18 |
| edge.dist.+pH+temp. | 6 | -1184,03 | 2381,286 | 84,68791 | 4,08E-19 |
| depth+diss.ox.+veget. | 6 | -1184,24 | 2381,695 | 85,0971 | 3,32E-19 |
| depth+diss.ox.+pH | 6 | -1184,27 | 2381,764 | 85,16662 | 3,21E-19 |
| depth+diss.ox.+edge.dist. | 6 | -1184,42 | 2382,065 | 85,46723 | 2,76E-19 |
| diss.ox.+pH+veget. | 6 | -1184,68 | 2382,569 | 85,97156 | 2,15E-19 |
| diss.ox.+edge.dist.+veget. | 6 | -1184,77 | 2382,748 | 86,15072 | 1,96E-19 |
| diss.ox.+edge.dist.+pH | 6 | -1185,33 | 2383,875 | 87,27713 | 1,12E-19 |
| depth+pH+veget. | 6 | -1185,58 | 2384,368 | 87,77031 | 8,73E-20 |
| depth+edge.dist.+veget. | 6 | -1185,85 | 2384,924 | 88,32609 | 6,61E-20 |
| depth+edge.dist.+pH | 6 | -1185,89 | 2385,002 | 88,40443 | 6,36E-20 |
| edge.dist.+pH+veget. | 5 | -1194,27 | 2399,389 | 102,7908 | 4,78E-23 |
| diss.ox.+temp. | 5 | -1195,45 | 2401,766 | 105,1682 | 1,46E-23 |
| depth+temp. | 5 | -1195,54 | 2401,935 | 105,3369 | 1,34E-23 |
| temp.+veget. | 5 | -1195,94 | 2402,73 | 106,1322 | 8,99E-24 |
| pH+temp. | 5 | -1196,19 | 2403,233 | 106,635 | 6,99E-24 |
| edge.dist.+temp. | 5 | -1198,13 | 2407,122 | 110,5241 | 1,00E-24 |
| depth+diss.ox. | 5 | -1198,34 | 2407,527 | 110,9297 | 8,16E-25 |
| diss.ox.+veget. | 5 | -1198,5 | 2407,849 | 111,2509 | 6,95E-25 |
| diss.ox.+pH | 5 | -1198,68 | 2408,218 | 111,6199 | 5,78E-25 |
| diss.ox.+edge.dist. | 5 | -1199,25 | 2409,364 | 112,766 | 3,26E-25 |
| depth+veget. | 5 | -1199,52 | 2409,894 | 113,2963 | 2,50E-25 |
| pH+veget. | 5 | -1199,62 | 2410,103 | 113,5049 | 2,25E-25 |
| depth+pH | 5 | -1199,81 | 2410,479 | 113,8807 | 1,87E-25 |
| depth+edge.dist. | 5 | -1199,85 | 2410,555 | 113,9575 | 1,80E-25 |
| edge.dist.+veget. | 5 | -1200,09 | 2411,036 | 114,4387 | 1,41E-25 |
| edge.dist.+pH | 4 | -1209,83 | 2428,214 | 131,6161 | 2,63E-29 |
| temp. | 4 | -1212,39 | 2433,346 | 136,748 | 2,02E-30 |
| diss.ox. | 4 | -1213,45 | 2435,462 | 138,8638 | 7,02E-31 |
| veget. | 4 | -1213,55 | 2435,666 | 139,068 | 6,34E-31 |
| depth | 4 | -1213,76 | 2436,081 | 139,4828 | 5,15E-31 |
| pH | 4 | -1214,06 | 2436,675 | 140,0769 | 3,83E-31 |
| edge.dist. | 3 | -1227,7 | 2461,726 | 165,1277 | 1,39E-36 |

**§12:** Summary of adjusted models produced by GLMM analysis relating local descriptors and phylogenetic redundancy in tadpole communities in ponds in the Spring (season with low hydric stress). Local descriptors: Depth (water column depth); Diss.ox. (dissolved oxygen); edge.dist. (edge distance); temp (water temperature); veget (percentage of aquatic vegetation cover).

**S12:** Summary of adjusted models produced by GLMM analysis relating local descriptors and phylogenetic redundancy in tadpole communities in ponds in the Spring (season with low hydric stress). Local descriptors: Depth (water column depth); Diss.ox. (dissolved oxygen); edge.dist. (edge distance); temp (water temperature); veget (percentage of aquatic vegetation cover).

| Model | df | logLik | AICc | $\Delta$ | weight |
| --- | --- | --- | --- | --- | --- |
| null | 9 | -3317,32 | 6653,532 | 0 | 0,999975 |
| depth+diss.ox.+edge.dist.+pH+temp.+veget. | 8 | -3330,42 | 6677,549 | 24,01641 | 6,09E-06 |
| depth+edge.dist.+pH+temp.+veget. | 8 | -3330,61 | 6677,938 | 24,40516 | 5,02E-06 |
| depth+diss.ox.+pH+temp.+veget. | 8 | -3330,81 | 6678,342 | 24,81008 | 4,10E-06 |
| depth+diss.ox.+edge.dist.+pH+veget. | 8 | -3330,85 | 6678,405 | 24,87251 | 3,97E-06 |
| depth+diss.ox.+edge.dist.+temp.+veget. | 8 | -3330,88 | 6678,481 | 24,94844 | 3,82E-06 |
| diss.ox.+edge.dist.+pH+temp.+veget. | 8 | -3331,52 | 6679,755 | 26,22215 | 2,02E-06 |
| depth+diss.ox.+edge.dist.+pH+temp. | 7 | -3343,71 | 6701,974 | 48,4421 | 3,03E-11 |
| depth+pH+temp.+veget. | 7 | -3343,9 | 6702,347 | 48,81488 | 2,51E-11 |
| depth+edge.dist.+pH+veget. | 7 | -3343,94 | 6702,441 | 48,90878 | 2,40E-11 |
| depth+edge.dist.+temp.+veget. | 7 | -3343,99 | 6702,529 | 48,99692 | 2,29E-11 |
| edge.dist.+pH+temp.+veget. | 7 | -3344,07 | 6702,696 | 49,16357 | 2,11E-11 |
| depth+diss.ox.+pH+veget. | 7 | -3344,27 | 6703,084 | 49,55195 | 1,74E-11 |
| diss.ox.+edge.dist.+temp.+veget. | 7 | -3344,28 | 6703,114 | 49,58141 | 1,71E-11 |
| depth+diss.ox.+temp.+veget. | 7 | -3344,31 | 6703,162 | 49,62995 | 1,67E-11 |
| diss.ox.+edge.dist.+pH+veget. | 7 | -3344,41 | 6703,366 | 49,8339 | 1,51E-11 |
| depth+diss.ox.+edge.dist.+veget. | 7 | -3344,42 | 6703,394 | 49,86141 | 1,49E-11 |
| diss.ox.+pH+temp.+veget. | 7 | -3344,63 | 6703,812 | 50,27984 | 1,21E-11 |
| depth+edge.dist.+pH+temp. | 7 | -3344,83 | 6704,204 | 50,67168 | 9,93E-12 |
| depth+diss.ox.+pH+temp. | 7 | -3344,99 | 6704,539 | 51,00673 | 8,39E-12 |
| depth+diss.ox.+edge.dist.+temp. | 7 | -3345,03 | 6704,618 | 51,08536 | 8,07E-12 |
| diss.ox.+edge.dist.+pH+temp. | 7 | -3345,05 | 6704,652 | 51,11997 | 7,93E-12 |
| depth+diss.ox.+edge.dist.+pH | 6 | -3357,16 | 6726,732 | 73,19955 | 1,27E-16 |
| depth+pH+veget. | 6 | -3357,36 | 6727,138 | 73,60566 | 1,04E-16 |
| edge.dist.+pH+veget. | 6 | -3357,37 | 6727,156 | 73,62323 | 1,03E-16 |
| edge.dist.+temp.+veget. | 6 | -3357,38 | 6727,17 | 73,63719 | 1,02E-16 |
| depth+temp.+veget. | 6 | -3357,49 | 6727,388 | 73,8558 | 9,17E-17 |
| depth+edge.dist.+veget. | 6 | -3357,52 | 6727,457 | 73,9242 | 8,86E-17 |
| pH+temp.+veget. | 6 | -3357,74 | 6727,896 | 74,36325 | 7,12E-17 |
| diss.ox.+edge.dist.+veget. | 6 | -3357,77 | 6727,946 | 74,41364 | 6,94E-17 |
| diss.ox.+pH+veget. | 6 | -3357,8 | 6728,019 | 74,48689 | 6,69E-17 |
| depth+diss.ox.+veget. | 6 | -3357,94 | 6728,286 | 74,7539 | 5,85E-17 |
| depth+pH+temp. | 6 | -3357,94 | 6728,289 | 74,75631 | 5,85E-17 |
| diss.ox.+temp.+veget. | 6 | -3358,1 | 6728,611 | 75,07815 | 4,98E-17 |
| depth+edge.dist.+temp. | 6 | -3358,14 | 6728,687 | 75,15455 | 4,79E-17 |
| edge.dist.+pH+temp. | 6 | -3358,15 | 6728,714 | 75,18142 | 4,73E-17 |
| depth+edge.dist.+pH | 6 | -3358,32 | 6729,047 | 75,51427 | 4,00E-17 |
| depth+diss.ox.+pH | 6 | -3358,38 | 6729,176 | 75,64349 | 3,75E-17 |
| diss.ox.+edge.dist.+temp. | 6 | -3358,43 | 6729,275 | 75,74257 | 3,57E-17 |
| depth+diss.ox.+temp. | 6 | -3358,48 | 6729,374 | 75,84146 | 3,40E-17 |
| diss.ox.+edge.dist.+pH | 6 | -3358,55 | 6729,518 | 75,98611 | 3,16E-17 |
| diss.ox.+pH+temp. | 6 | -3358,59 | 6729,583 | 76,05024 | 3,06E-17 |
| depth+diss.ox.+edge.dist. | 5 | -3370,8 | 6751,892 | 98,35993 | 4,38E-22 |
| edge.dist.+veget. | 5 | -3370,82 | 6751,942 | 98,40985 | 4,27E-22 |
| pH+veget. | 5 | -3370,89 | 6752,077 | 98,54483 | 3,99E-22 |
| depth+veget. | 5 | -3371,04 | 6752,375 | 98,8427 | 3,44E-22 |
| temp.+veget. | 5 | -3371,33 | 6752,958 | 99,42592 | 2,57E-22 |
| diss.ox.+veget. | 5 | -3371,43 | 6753,143 | 99,61106 | 2,34E-22 |
| depth+pH | 5 | -3371,49 | 6753,266 | 99,73386 | 2,20E-22 |
| edge.dist.+temp. | 5 | -3371,54 | 6753,368 | 99,836 | 2,09E-22 |
| depth+temp. | 5 | -3371,55 | 6753,4 | 99,86732 | 2,06E-22 |
| edge.dist.+pH | 5 | -3371,66 | 6753,603 | 100,0705 | 1,86E-22 |
| pH+temp. | 5 | -3371,68 | 6753,651 | 100,119 | 1,82E-22 |
| depth+edge.dist. | 5 | -3371,88 | 6754,056 | 100,5241 | 1,48E-22 |
| diss.ox.+edge.dist. | 5 | -3371,92 | 6754,142 | 100,6099 | 1,42E-22 |
| diss.ox.+pH | 5 | -3371,99 | 6754,263 | 100,7309 | 1,34E-22 |
| depth+diss.ox. | 5 | -3372,02 | 6754,342 | 100,8093 | 1,29E-22 |
| diss.ox.+temp. | 4 | -3384,39 | 6776,975 | 123,4427 | 1,57E-27 |
| veget. | 4 | -3384,95 | 6778,101 | 124,5685 | 8,92E-28 |
| edge.dist. | 4 | -3385 | 6778,187 | 124,6548 | 8,54E-28 |
| pH | 4 | -3385,09 | 6778,37 | 124,8374 | 7,80E-28 |
| depth | 4 | -3385,13 | 6778,447 | 124,9148 | 7,50E-28 |
| temp. | 4 | -3385,44 | 6779,078 | 125,5452 | 5,47E-28 |
| diss.ox. | 3 | -3398,51 | 6803,141 | 149,6089 | 3,26E-33 |

**§13:** Summary of adjusted models produced by GLMM analysis relating local descriptors and phylogenetic redundancy in tadpole communities in ponds in the Summer (season with high hydric stress). Local descriptors: Depth (water column depth); Diss.ox. (dissolved oxygen); edge.dist. (edge distance); temp (water temperature); veget (percentage of aquatic vegetation cover).

**S13:**
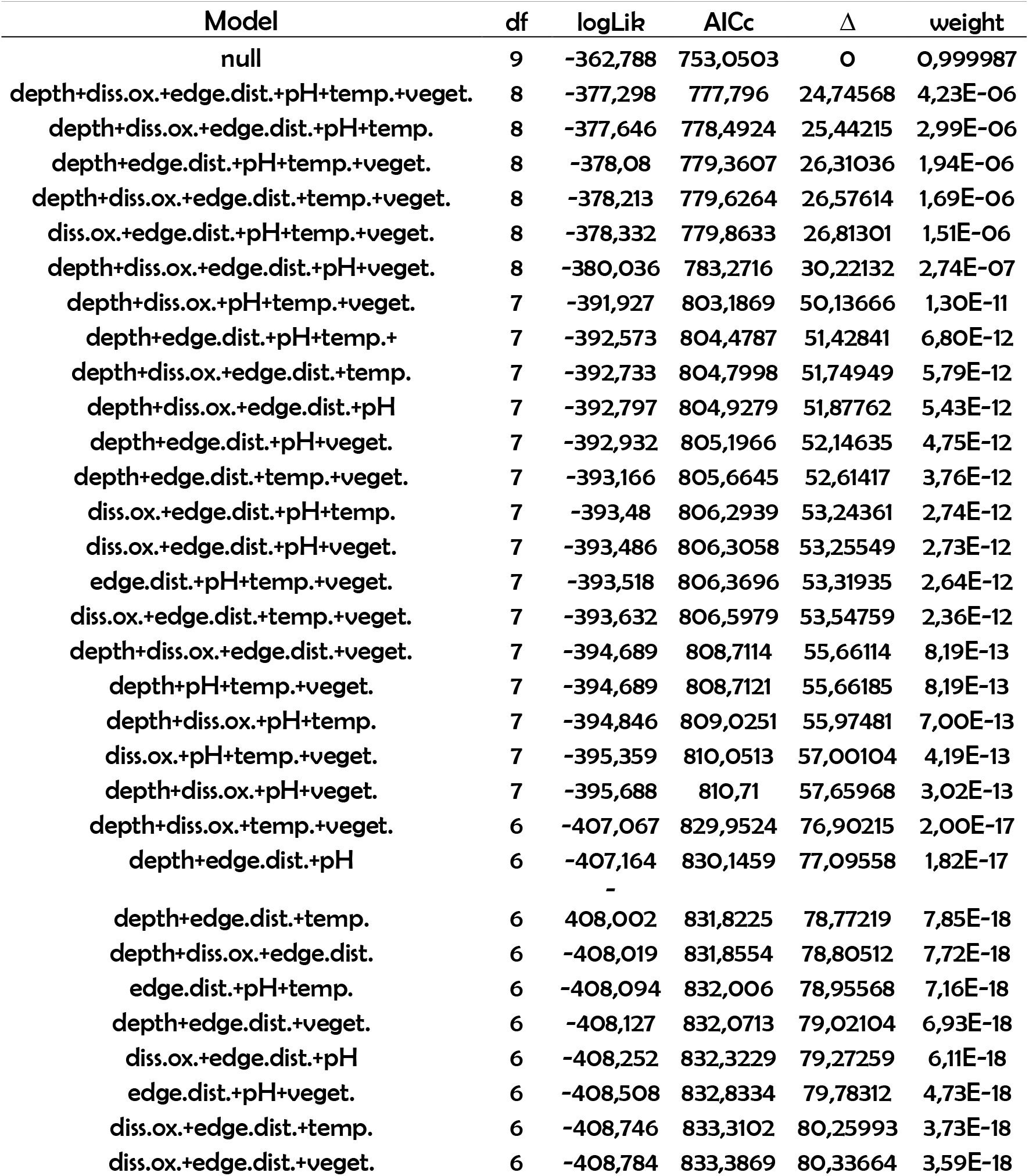
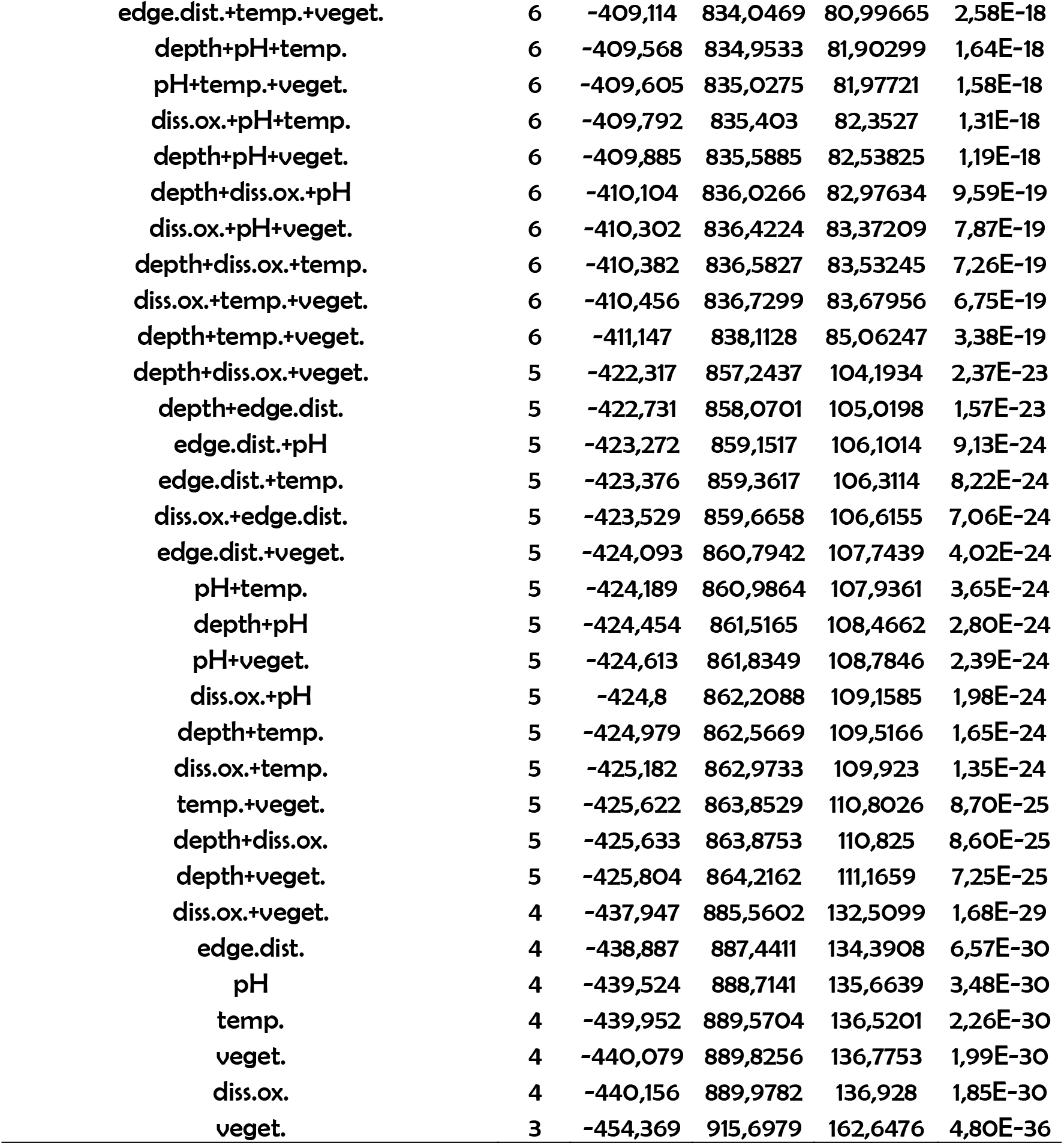
Summary of adjusted models produced by GLMM analysis relating local descriptors and phylogenetic redundancy in tadpole communities in ponds in the Summer (season with high hydric stress). Local descriptors: Depth (water column depth); Diss.ox. (dissolved oxygen); edge.dist. (edge distance); temp (water temperature); veget (percentage of aquatic vegetation cover).

**S14:** Summary of adjusted models produced by GLMM analysis relating local descriptors and phylogenetic redundancy in tadpole communities in ponds in the Autumn (season with high hydric stress). Local descriptors: Depth (water column depth); Diss.ox. (dissolved oxygen); edge.dist. (edge distance); temp (water temperature); veget (percentage of aquatic vegetation cover).

| Model | df | logLik | AICc | $\Delta$ | weight |
| --- | --- | --- | --- | --- | --- |
| null | 9 | -319,557 | 667,7015 | 0 | 0,999976 |
| depth+diss.ox.+edge.dist.+pH+temp.+veget. | 8 | -334,026 | 692,0525 | 24,35106 | 5,15E-06 |
| depth+diss.ox.+edge.dist.+pH+temp. | 8 | -334,137 | 692,2743 | 24,57287 | 4,61E-06 |
| depth+diss.ox.+edge.dist.+temp.+veget. | 8 | -334,192 | 692,3835 | 24,68209 | 4,37E-06 |
| diss.ox.+edge.dist.+pH+temp.+veget. | 8 | -334,329 | 692,6579 | 24,95649 | 3,81E-06 |
| depth+edge.dist.+pH+temp.+veget. | 8 | -334,345 | 692,6906 | 24,98917 | 3,75E-06 |
| depth+diss.ox.+edge.dist.+pH+veget. | 8 | -334,726 | 693,451 | 25,74955 | 2,56E-06 |
| depth+diss.ox.+pH+temp.+veget. | 7 | -348,424 | 716,7418 | 49,04038 | 2,24E-11 |
| depth+diss.ox.+edge.dist.+veget. | 7 | -348,475 | 716,845 | 49,14353 | 2,13E-11 |
| edge.dist.+pH+temp.+veget. | 7 | -348,603 | 717,1016 | 49,40019 | 1,87E-11 |
| depth+diss.ox.+edge.dist.+temp. | 7 | -348,657 | 717,2078 | 49,50633 | 1,78E-11 |
| diss.ox.+edge.dist.+pH+temp. | 7 | -348,744 | 717,3836 | 49,68216 | 1,63E-11 |
| diss.ox.+edge.dist.+temp.+veget. | 7 | -348,79 | 717,4743 | 49,77281 | 1,56E-11 |
| depth+edge.dist.+pH+temp. | 7 | -348,836 | 717,5665 | 49,86503 | 1,49E-11 |
| depth+edge.dist.+temp.+veget. | 7 | -348,918 | 717,7317 | 50,0302 | 1,37E-11 |
| depth+diss.ox.+edge.dist.+pH | 7 | -348,936 | 717,767 | 50,06558 | 1,34E-11 |
| diss.ox.+edge.dist.+pH+veget. | 7 | -349,08 | 718,0551 | 50,35367 | 1,16E-11 |
| depth+edge.dist.+pH+veget. | 7 | -349,358 | 718,6109 | 50,90942 | 8,81E-12 |
| depth+diss.ox.+temp.+veget. | 7 | -349,423 | 718,74 | 51,03854 | 8,26E-12 |
| diss.ox.+pH+temp.+veget. | 7 | -349,509 | 718,9118 | 51,21033 | 7,58E-12 |
| depth+diss.ox.+temp.+veget. | 7 | -349,527 | 718,9494 | 51,24795 | 7,44E-12 |
| depth+diss.ox.+pH+temp. | 7 | -349,564 | 719,0218 | 51,32036 | 7,18E-12 |
| depth+pH+temp.+veget. | 6 | -362,888 | 741,9768 | 74,27532 | 7,44E-17 |
| diss.ox.+edge.dist.+veget. | 6 | -362,93 | 742,0591 | 74,35769 | 7,14E-17 |
| edge.dist.+temp.+veget. | 6 | -362,947 | 742,0931 | 74,39159 | 7,02E-17 |
| edge.dist.+pH+veget. | 6 | -363,114 | 742,4278 | 74,72636 | 5,93E-17 |
| depth+edge.dist.+veget. | 6 | -363,21 | 742,6206 | 74,91912 | 5,39E-17 |
| diss.ox.+edge.dist.+temp. | 6 | -363,301 | 742,8024 | 75,10099 | 4,92E-17 |
| edge.dist.+pH+temp. | 6 | -363,311 | 742,8216 | 75,12017 | 4,87E-17 |
| depth+diss.ox.+edge.dist. | 6 | -363,316 | 742,8311 | 75,12961 | 4,85E-17 |
| depth+edge.dist.+temp. | 6 | -363,494 | 743,1884 | 75,48698 | 4,06E-17 |
| diss.ox.+edge.dist.+pH | 6 | -363,633 | 743,4653 | 75,76382 | 3,53E-17 |
| depth+diss.ox.+veget. | 6 | -363,633 | 743,4664 | 75,76491 | 3,53E-17 |
| depth+edge.dist.+pH | 6 | -363,657 | 743,5145 | 75,81308 | 3,45E-17 |
| pH+temp.+veget. | 6 | -364,028 | 744,2551 | 76,55361 | 2,38E-17 |
| diss.ox.+veget.+temp. | 6 | -364,108 | 744,4154 | 76,71391 | 2,20E-17 |
| depth+temp.+veget. | 6 | -364,15 | 744,4998 | 76,79834 | 2,11E-17 |
| depth+diss.ox.+temp. | 6 | -364,162 | 744,523 | 76,82157 | 2,08E-17 |
| diss.ox.+pH+veget. | 6 | -364,179 | 744,5588 | 76,85731 | 2,04E-17 |
| diss.ox.+pH+temp. | 6 | -364,321 | 744,8418 | 77,14034 | 1,77E-17 |
| depth+pH+veget. | 6 | -364,366 | 744,9322 | 77,23077 | 1,70E-17 |
| depth+pH+temp. | 6 | -364,379 | 744,9571 | 77,2556 | 1,68E-17 |
| depth+diss.ox.+pH | 5 | -376,839 | 766,5358 | 98,83439 | 3,45E-22 |
| edge.dist.+veget. | 5 | -377,809 | 768,4754 | 100,7739 | 1,31E-22 |
| edge.dist.+pH | 5 | -377,831 | 768,5193 | 100,8178 | 1,28E-22 |
| diss.ox.+edge.dist. | 5 | -377,935 | 768,727 | 101,0255 | 1,15E-22 |
| edge.dist.+temp. | 5 | -378,108 | 769,073 | 101,3715 | 9,72E-23 |
| diss.ox.+veget. | 5 | -378,124 | 769,1049 | 101,4035 | 9,56E-23 |
| temp.+veget. | 5 | -378,145 | 769,1463 | 101,4449 | 9,37E-23 |
| depth+edge.dist. | 5 | -378,178 | 769,2129 | 101,5114 | 9,06E-23 |
| pH+veget. | 5 | -378,327 | 769,5113 | 101,8099 | 7,80E-23 |
| depth+veget. | 5 | -378,435 | 769,727 | 102,0255 | 7,01E-23 |
| depth+diss.ox. | 5 | -378,766 | 770,389 | 102,6875 | 5,03E-23 |
| pH+temp. | 5 | -378,78 | 770,4162 | 102,7147 | 4,96E-23 |
| diss.ox.+temp. | 5 | -378,927 | 770,7111 | 103,0096 | 4,28E-23 |
| depth+temp. | 5 | -378,989 | 770,836 | 103,1345 | 4,02E-23 |
| diss.ox.+pH | 5 | -379,171 | 771,1996 | 103,4982 | 3,35E-23 |
| depth+pH | 4 | -392,058 | 793,9341 | 126,2326 | 3,88E-28 |
| edge.dist. | 4 | -392,129 | 794,0763 | 126,3748 | 3,61E-28 |
| veget. | 4 | -392,956 | 795,7309 | 128,0294 | 1,58E-28 |
| diss.ox. | 4 | -393,267 | 796,3519 | 128,6505 | 1,16E-28 |
| depth | 4 | -393,339 | 796,4972 | 128,7957 | 1,08E-28 |
| pH | 4 | -393,346 | 796,5108 | 128,8093 | 1,07E-28 |
| temp. | 3 | -407,251 | 821,5445 | 153,843 | 3,92E-34 |

**§15:** Summary of adjusted models produced by GLMM analysis relating local descriptors and functional diversity in tadpole communities in ponds in the Winter (season with low hydric stress). Local descriptors: Depth (water column depth); Diss.ox. (dissolved oxygen); edge.dist. (edge distance); temp (water temperature); veget (percentage of aquatic vegetation cover).

**S15:** Summary of adjusted models produced by GLMM analysis relating local descriptors and functional diversity in tadpole communities in ponds in the Winter (season with low hydric stress). Local descriptors: Depth (water column depth); Diss.ox. (dissolved oxygen); edge.dist. (edge distance); temp (water temperature); veget (percentage of aquatic vegetation cover).

| Model | df | logLik | AICc | $\Delta$ | weight |
| --- | --- | --- | --- | --- | --- |
| null | 3 | 19,47235 | -32,6114 | 0 | 0,540167 |
| depth | 4 | 19,67615 | -30,7889 | 1,822444 | 0,230381 |
| pH | 4 | 18,03736 | -27,5113 | 5,100023 | 0,042801 |
| temp. | 4 | 17,81358 | -27,0638 | 5,54759 | 0,034219 |
| diss.ox. | 4 | 17,78358 | -27,0038 | 5,60759 | 0,033208 |
| depth+temp. | 5 | 18,5956 | -26,3341 | 6,27731 | 0,023758 |
| depth+diss.ox. | 5 | 18,3008 | -25,7445 | 6,866907 | 0,017692 |
| veget. | 4 | 17,11535 | -25,6673 | 6,944041 | 0,017023 |
| edge.dist. | 4 | 17,07823 | -25,5931 | 7,018286 | 0,016403 |
| depth+pH | 5 | 17,91449 | -24,9718 | 7,639518 | 0,012023 |
| depth+veget. | 5 | 17,33034 | -23,8035 | 8,807818 | 0,006704 |
| depth+edge.dist. | 5 | 16,90686 | -22,9566 | 9,654784 | 0,004389 |
| pH+temp. | 5 | 16,86829 | -22,8794 | 9,731919 | 0,004223 |
| depth+pH+temp. | 6 | 17,23906 | -21,2607 | 11,35062 | 0,00188 |
| diss.ox.+pH | 5 | 16,03882 | -21,2205 | 11,39087 | 0,001843 |
| diss.ox.+temp. | 5 | 16,01317 | -21,1692 | 11,44216 | 0,001796 |
| depth+diss.ox.+temp. | 6 | 17,17021 | -21,123 | 11,48834 | 0,001755 |
| temp.+veget. | 5 | 15,93205 | -21,007 | 11,6044 | 0,001656 |
| edge.dist.+temp. | 5 | 15,66839 | -20,4796 | 12,13173 | 0,001272 |
| depth+temp.+veget. | 6 | 16,75658 | -20,2958 | 12,3156 | 0,00116 |
| pH+veget. | 5 | 15,42347 | -19,9898 | 12,62157 | 0,000996 |
| edge.dist.+pH | 5 | 15,42343 | -19,9897 | 12,62165 | 0,000996 |
| diss.ox.+edge.dist. | 5 | 15,22067 | -19,5842 | 13,02717 | 0,000813 |
| diss.ox.+veget. | 5 | 15,18125 | -19,5054 | 13,106 | 0,000782 |
| depth+diss.ox.+pH | 6 | 16,12662 | -19,0358 | 13,57552 | 0,000618 |
| edge.dist.+veget. | 5 | 14,87267 | -18,8882 | 13,72317 | 0,000574 |
| depth+edge.dist.+temp. | 6 | 15,951 | -18,6846 | 13,92675 | 0,000519 |
| depth+diss.ox.+veget. | 6 | 15,58109 | -17,9448 | 14,66657 | 0,000358 |
| depth+pH+veget. | 6 | 15,38777 | -17,5581 | 15,05321 | 0,000295 |
| depth+diss.ox.+edge.dist. | 6 | 15,3811 | -17,5448 | 15,06655 | 0,000293 |
| depth+edge.dist.+pH | 6 | 15,05095 | -16,8845 | 15,72684 | 0,000211 |
| diss.ox.+pH+temp. | 6 | 14,79322 | -16,3691 | 16,24231 | 0,000163 |
| depth+edge.dist.+veget. | 6 | 14,70132 | -16,1852 | 16,42612 | 0,000149 |
| pH+temp.+veget. | 6 | 14,52758 | -15,8378 | 16,77359 | 0,000125 |
| edge.dist.+pH+temp. | 6 | 14,31518 | -15,413 | 17,19839 | 0,000101 |
| depth+diss.ox.+pH+temp. | 7 | 15,50482 | -15,3626 | 17,24878 | 9,85E-05 |
| diss.ox.+temp.+veget. | 6 | 13,88112 | -14,5448 | 18,06651 | 6,54E-05 |
| edge.dist.+temp.+veget. | 6 | 13,85969 | -14,502 | 18,10938 | 6,40E-05 |
| depth+diss.ox.+temp.+veget. | 7 | 15,00503 | -14,363 | 18,24836 | 5,97E-05 |
| depth+pH+temp.+veget. | 7 | 14,95643 | -14,2658 | 18,34556 | 5,69E-05 |
| diss.ox.+edge.dist.+temp. | 6 | 13,64041 | -14,0634 | 18,54793 | 5,14E-05 |
| diss.ox.+edge.dist.+pH | 6 | 13,29169 | -13,366 | 19,24537 | 3,63E-05 |
| diss.ox.+pH+veget. | 6 | 13,28852 | -13,3597 | 19,25171 | 3,62E-05 |
| depth+edge.dist.+pH+temp. | 7 | 14,38047 | -13,1139 | 19,49747 | 3,20E-05 |
| depth+diss.ox.+edge.dist.+temp. | 7 | 14,37055 | -13,094 | 19,51733 | 3,17E-05 |
| depth+edge.dist.+temp.+veget. | 7 | 14,17706 | -12,7071 | 19,9043 | 2,61E-05 |
| edge.dist.+pH+veget. | 6 | 12,94 | -12,6626 | 19,94874 | 2,55E-05 |
| diss.ox.+edge.dist.+veget. | 6 | 12,63961 | -12,0618 | 20,54952 | 1,89E-05 |
| depth+diss.ox.+pH+veget. | 7 | 13,38355 | -11,12 | 21,49132 | 1,18E-05 |
| depth+diss.ox.+edge.dist.+pH | 7 | 13,15523 | -10,6634 | 21,94795 | 9,40E-06 |
| depth+diss.ox.+edge.dist.+veget. | 7 | 12,7026 | -9,75814 | 22,85322 | 5,98E-06 |
| depth+edge.dist.+pH+veget. | 7 | 12,63028 | -9,61351 | 22,99785 | 5,56E-06 |
| diss.ox.+pH+temp.+veget. | 7 | 12,30073 | -8,9544 | 23,65696 | 4,00E-06 |
| diss.ox.+edge.dist.+pH+temp. | 7 | 12,18229 | -8,71752 | 23,89384 | 3,55E-06 |
| edge.dist.+pH+temp.+veget. | 7 | 12,05135 | -8,45565 | 24,15571 | 3,12E-06 |
| depth+diss.ox.+pH+temp.+veget. | 8 | 13,02451 | -7,89976 | 24,7116 | 2,36E-06 |
| diss.ox.+edge.dist.+temp.+veget. | 7 | 11,5723 | -7,49754 | 25,11382 | 1,93E-06 |
| depth+diss.ox.+edge.dist.+pH+temp. | 8 | 12,58544 | -7,02162 | 25,58974 | 1,52E-06 |
| depth+diss.ox.+edge.dist.+temp.+veget. | 8 | 12,24789 | -6,34653 | 26,26483 | 1,09E-06 |
| depth+edge.dist.+pH+temp.+veget. | 8 | 12,15362 | -6,15799 | 26,45337 | 9,88E-07 |
| diss.ox.+edge.dist.+pH+veget. | 7 | 10,60374 | -5,56043 | 27,05093 | 7,33E-07 |
| depth+diss.ox.+edge.dist.+pH+veget. | 8 | 10,47944 | -2,80963 | 29,80173 | 1,85E-07 |
| diss.ox.+edge.dist.+pH+temp.+veget. | 8 | 9,754287 | -1,35932 | 31,25204 | 8,97E-08 |
| depth+diss.ox.+edge.dist.+pH+temp.+veget. | 9 | 10,15102 | 0,425225 | 33,03659 | 3,67E-08 |

**§16:** Summary of adjusted models produced by GLMM analysis relating local descriptors and functional diversity in tadpole communities in ponds in the Spring (season with low hydric stress). Local descriptors: Depth (water column depth); Diss.ox. (dissolved oxygen); edge.dist. (edge distance); temp (water temperature); veget (percentage of aquatic vegetation cover).

**S16:**
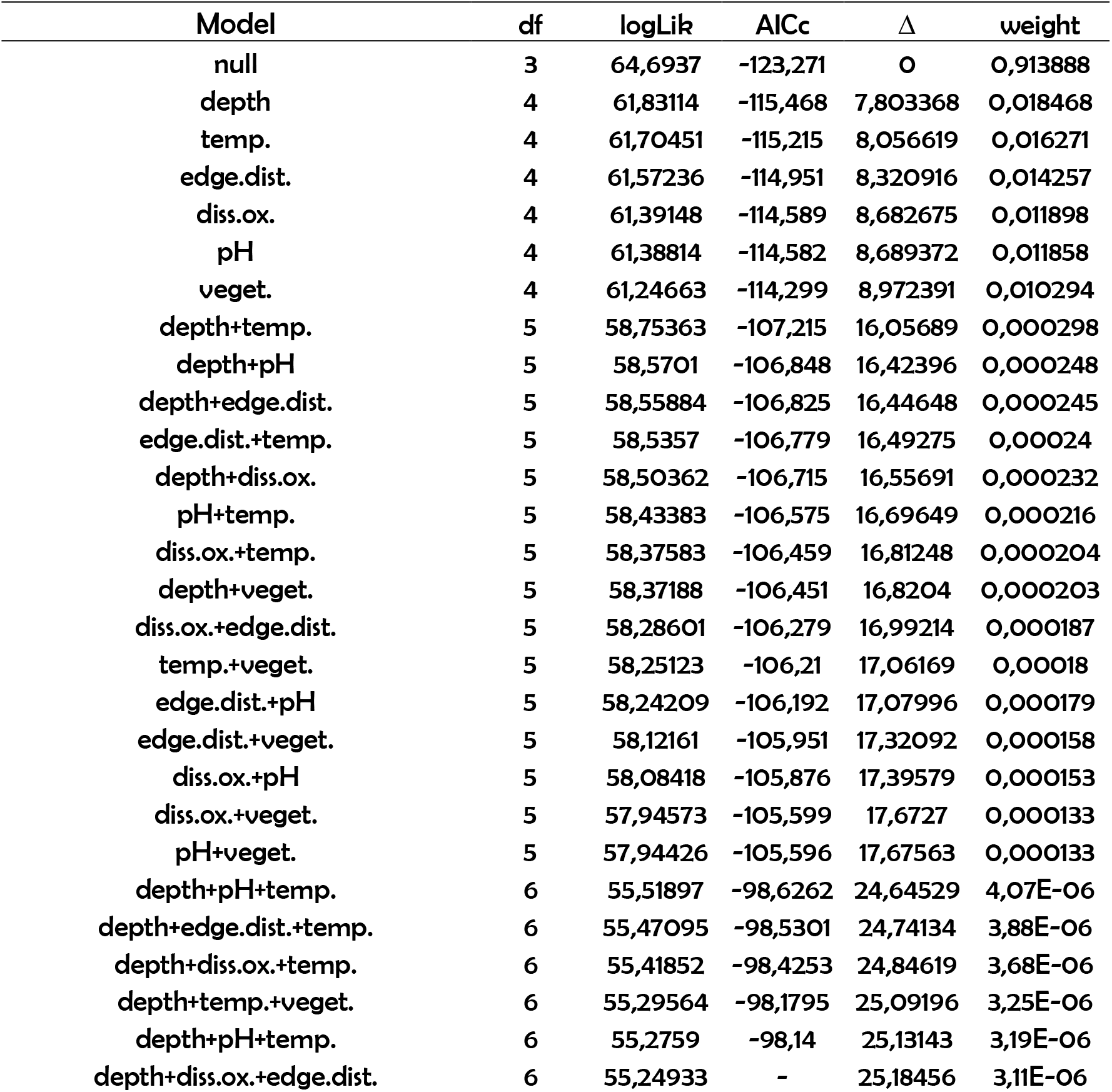
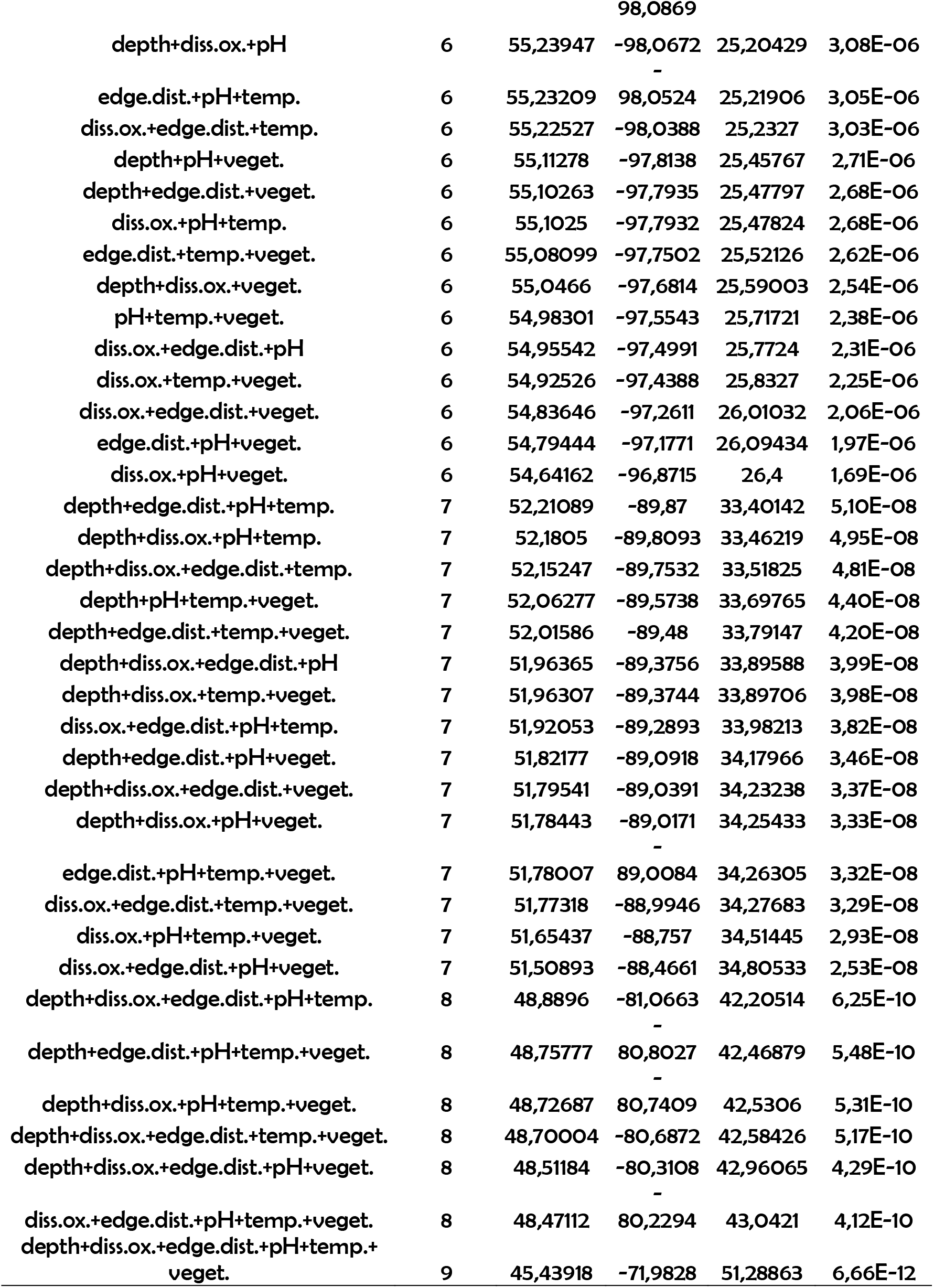
Summary of adjusted models produced by GLMM analysis relating local descriptors and functional diversity in tadpole communities in ponds in the Spring (season with low hydric stress). Local descriptors: Depth (water column depth); Diss.ox. (dissolved oxygen); edge.dist. (edge distance); temp (water temperature); veget (percentage of aquatic vegetation cover).

**S17:**
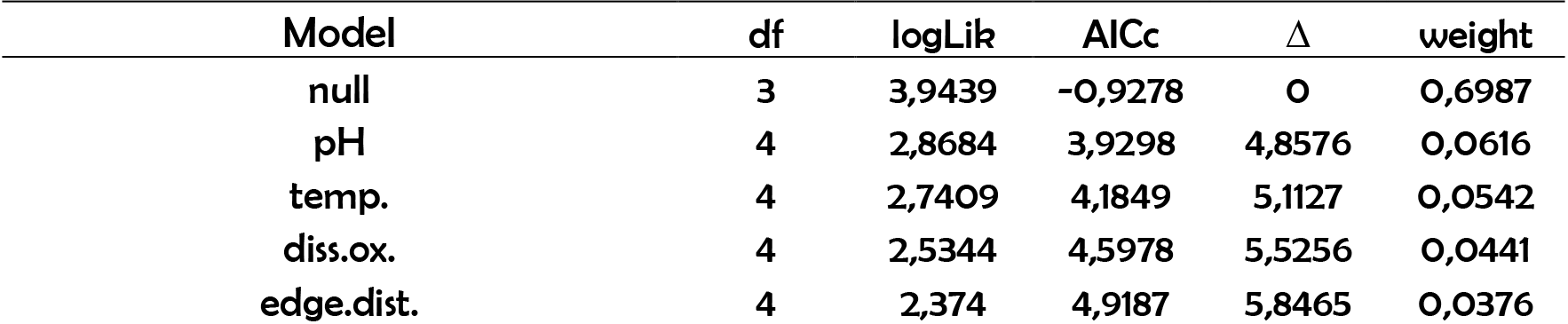
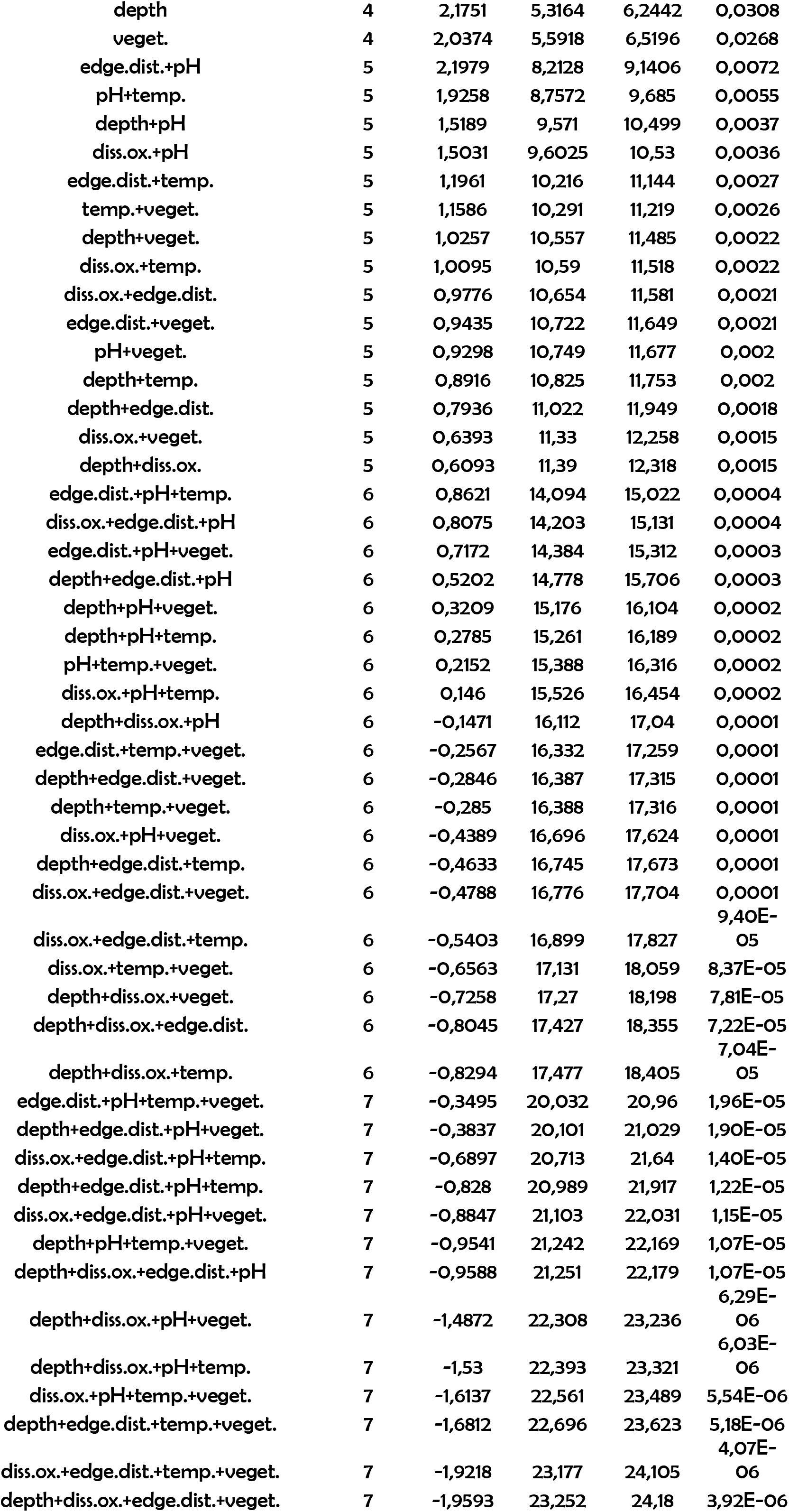
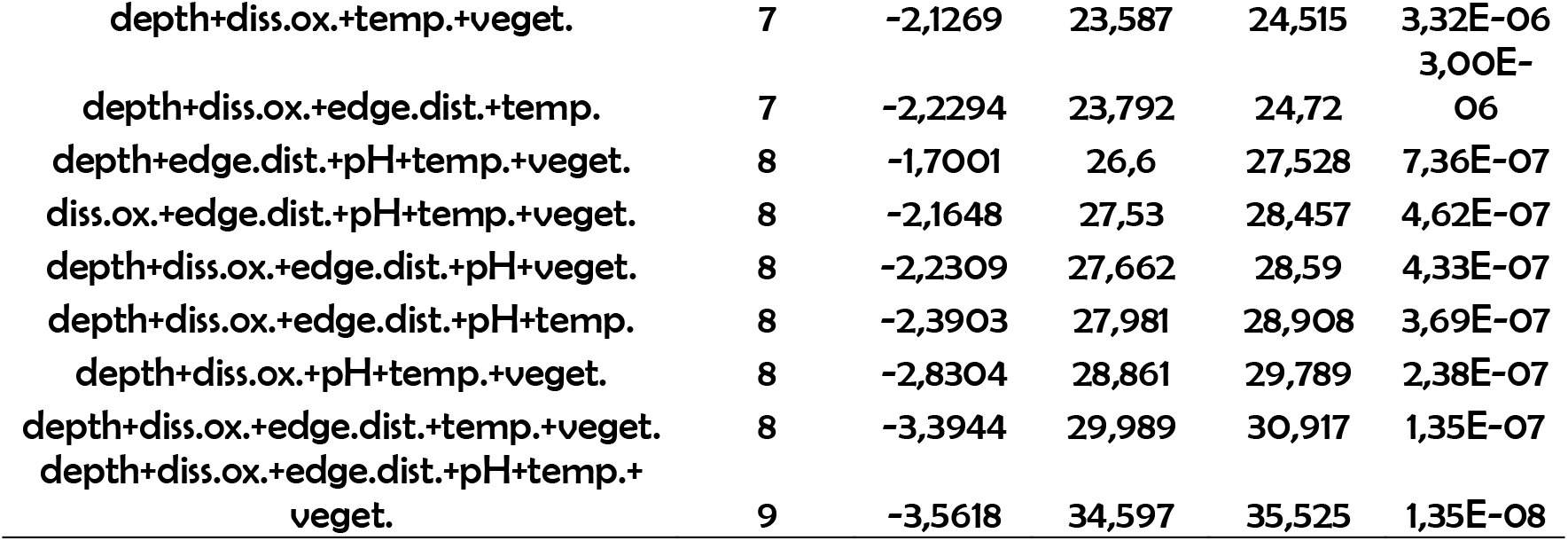
Summary of adjusted models produced by GLMM analysis relating local descriptors and functional diversity in tadpole communities in ponds in the Summer (season with high hydric stress). Local descriptors: Depth (water column depth); Diss.ox. (dissolved oxygen); edge.dist. (edge distance); temp (water temperature); veget (percentage of aquatic vegetation cover).

**S18:**
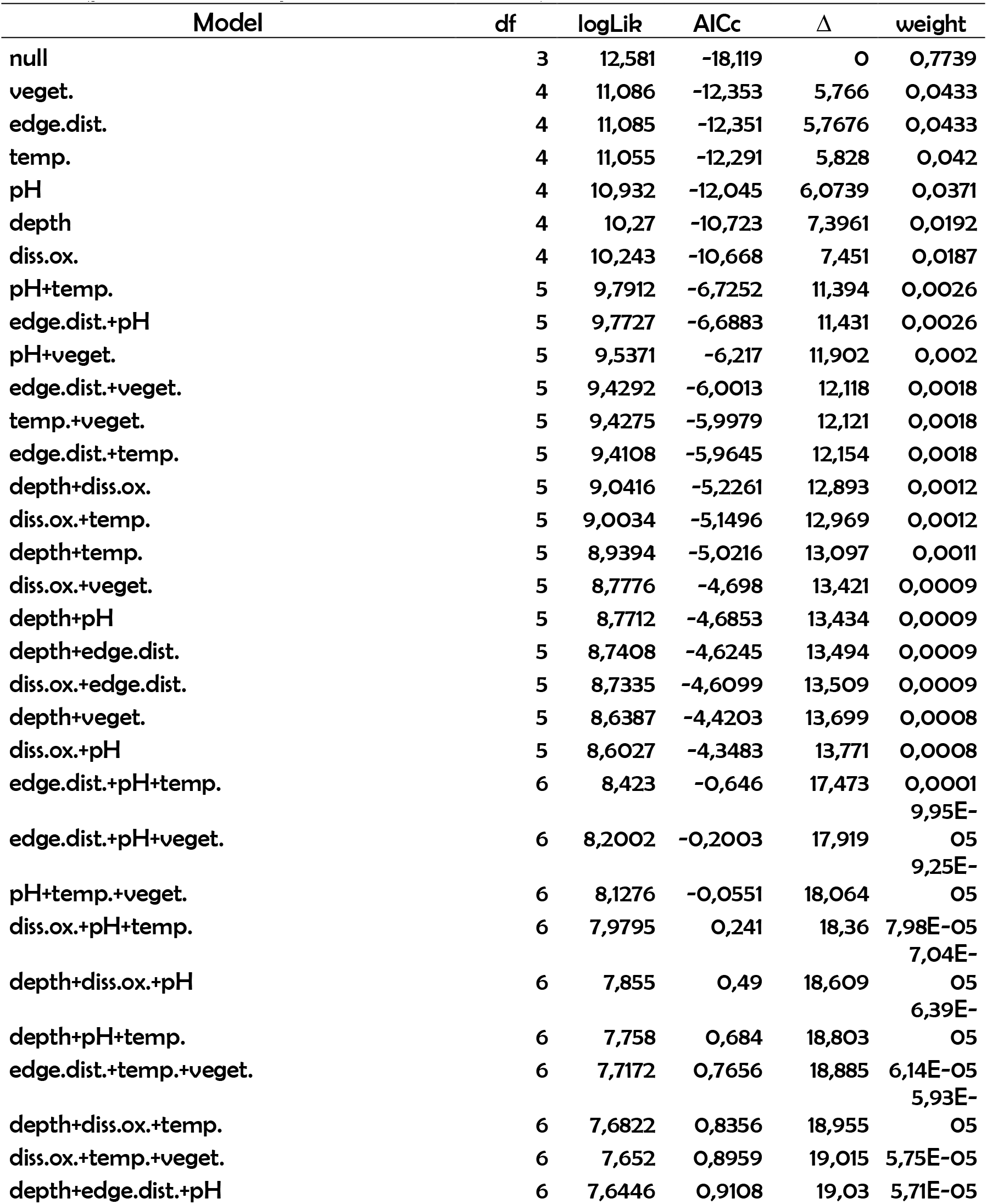
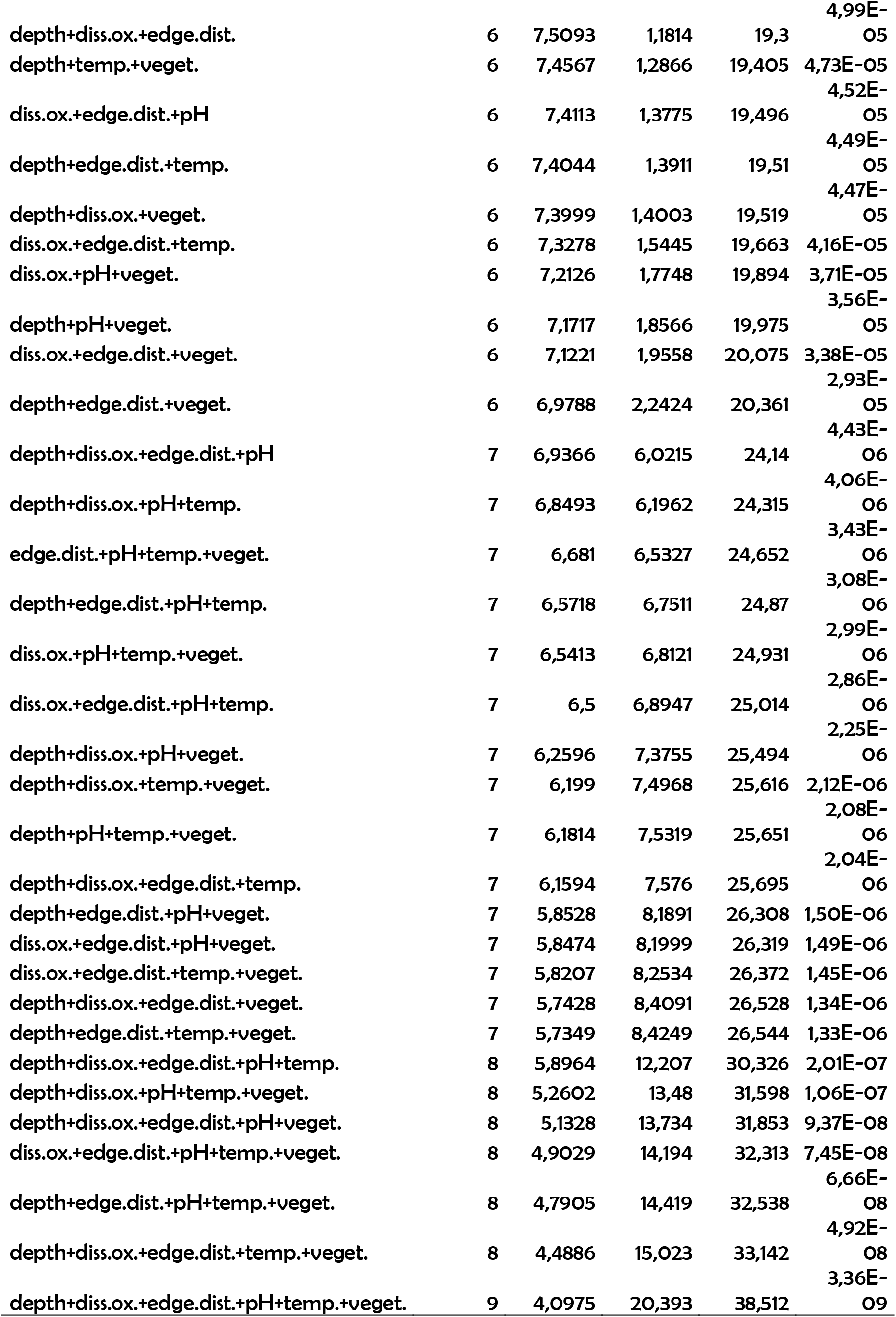
Summary of adjusted models produced by GLMM analysis relating local descriptors and functional diversity in tadpole communities in ponds in the Autumn (season with high hydric stress). Local descriptors: Depth (water column depth); Diss.ox. (dissolved oxygen); edge.dist. (edge distance); temp (water temperature); veget (percentage of aquatic vegetation cover).

**S19:** Summary of adjusted models produced by GLMM analysis relating local descriptors and functional redundancy in tadpole communities in ponds in the Winter (season with low hydric stress). Local descriptors: Depth (water column depth); Diss.ox. (dissolved oxygen); edge.dist. (edge distance); temp (water temperature); veget (percentage of aquatic vegetation cover).

| Model | df | logLik | AICc | $\Delta$ | weight |
| --- | --- | --- | --- | --- | --- |
| null | 3 | 51,198 | -96,062 | 0 | 0,5129 |
| depth | 4 | 51,645 | -94,727 | 1,3351 | 0,2631 |
| pH | 4 | 50,584 | -92,605 | 3,4572 | 0,0911 |
| depth+pH | 5 | 50,615 | -90,373 | 5,6887 | 0,0298 |
| edge.dist. | 4 | 49,005 | -89,447 | 6,6153 | 0,0188 |
| diss.ox. | 4 | 48,962 | -89,36 | 6,7017 | 0,018 |
| depth+diss.ox. | 5 | 50,074 | -89,291 | 6,7705 | 0,0174 |
| temp. | 4 | 48,546 | -88,529 | 7,5331 | 0,0119 |
| veget. | 4 | 48,168 | -87,772 | 8,2895 | 0,0081 |
| depth+temp. | 5 | 49,048 | -87,24 | 8,8221 | 0,0062 |
| depth+edge.dist. | 5 | 48,804 | -86,751 | 9,3114 | 0,0049 |
| depth+veget. | 5 | 48,58 | -86,303 | 9,7586 | 0,0039 |
| diss.ox.+pH | 5 | 48,228 | -85,599 | 10,463 | 0,0027 |
| edge.dist.+pH | 5 | 47,866 | -84,875 | 11,187 | 0,0019 |
| pH+temp. | 5 | 47,842 | -84,826 | 11,236 | 0,0019 |
| depth+diss.ox.+pH | 6 | 48,686 | -84,154 | 11,908 | 0,0013 |
| pH+veget. | 5 | 47,389 | -83,921 | 12,141 | 0,0012 |
| depth+pH+temp. | 6 | 48,067 | -82,916 | 13,146 | 0,0007 |
| diss.ox.+edge.dist. | 5 | 46,563 | -82,27 | 13,792 | 0,0005 |
| edge.dist.+temp. | 5 | 46,434 | -82,011 | 14,051 | 0,0005 |
| depth+edge.dist.+pH | 6 | 47,472 | -81,726 | 14,336 | 0,0004 |
| depth+pH+veget. | 6 | 47,379 | -81,54 | 14,522 | 0,0004 |
| diss.ox.+temp. | 5 | 46,126 | -81,394 | 14,668 | 0,0003 |
| edge.dist.+veget. | 5 | 45,995 | -81,133 | 14,929 | 0,0003 |
| depth+diss.ox.+temp. | 6 | 47,174 | -81,131 | 14,931 | 0,0003 |
| diss.ox.+veget. | 5 | 45,88 | -80,903 | 15,159 | 0,0003 |
| depth+diss.ox.+edge.dist. | 6 | 47,053 | -80,889 | 15,173 | 0,0003 |
| depth+diss.ox.+veget. | 6 | 46,882 | -80,546 | 15,516 | 0,0002 |
| temp.+veget. | 5 | 45,533 | -80,21 | 15,852 | 0,0002 |
| depth+edge.dist.+temp. | 6 | 46,283 | -79,349 | 16,713 | 0,0001 |
| depth+temp.+veget. | 6 | 46,048 | -78,879 | 17,183 | 9,52E-05 |
| depth+edge.dist.+veget. | 6 | 45,754 | -78,29 | 17,772 | 7,10E-05 |
| diss.ox.+pH+temp. | 6 | 45,398 | -77,578 | 18,484 | 4,97E-05 |
| diss.ox.+edge.dist.+pH | 6 | 45,374 | -77,531 | 18,531 | 4,85E-05 |
| edge.dist.+pH+temp. | 6 | 45,206 | -77,194 | 18,868 | 4,10E-05 |
| diss.ox.+pH+veget. | 6 | 45,01 | -76,803 | 19,259 | 3,37E-05 |
| depth+diss.ox.+pH+temp. | 7 | 46,071 | -76,495 | 19,567 | 2,89E-05 |
| pH+temp.+veget. | 6 | 44,696 | -76,175 | 19,887 | 2,46E-05 |
| edge.dist.+pH+veget. | 6 | 44,688 | -76,158 | 19,904 | 2,44E-05 |
| depth+diss.ox.+edge.dist.+pH | 7 | 45,439 | -75,231 | 20,831 | 1,54E-05 |
| depth+diss.ox.+pH+veget. | 7 | 45,425 | -75,203 | 20,859 | 1,52E-05 |
| diss.ox.+edge.dist.+temp. | 6 | 43,82 | -74,423 | 21,639 | 1,03E-05 |
| depth+edge.dist.+pH+temp. | 7 | 44,979 | -74,31 | 21,751 | 9,70E-06 |
| depth+pH+temp.+veget. | 7 | 44,901 | -74,155 | 21,907 | 8,98E-06 |
| diss.ox.+edge.dist.+veget. | 6 | 43,478 | -73,739 | 22,323 | 7,29E-06 |
| edge.dist.+temp.+veget. | 6 | 43,463 | -73,709 | 22,353 | 7,18E-06 |
| diss.ox.+temp.+veget. | 6 | 43,074 | -72,931 | 23,131 | 4,87E-06 |
| depth+edge.dist.+pH+veget. | 7 | 44,25 | -72,854 | 23,208 | 4,68E-06 |
| depth+diss.ox.+edge.dist.+temp. | 7 | 44,22 | -72,794 | 23,268 | 4,54E-06 |
| depth+diss.ox.+temp.+veget. | 7 | 44,142 | -72,636 | 23,426 | 4,20E-06 |
| depth+diss.ox.+edge.dist.+veget. | 7 | 43,865 | -72,083 | 23,979 | 3,19E-06 |
| depth+edge.dist.+temp.+veget. | 7 | 43,321 | -70,995 | 25,067 | 1,85E-06 |
| diss.ox.+edge.dist.+pH+temp. | 7 | 42,614 | -69,582 | 26,48 | 9,12E-07 |
| diss.ox.+pH+temp.+veget. | 7 | 42,226 | -68,804 | 27,258 | 6,18E-07 |
| diss.ox.+edge.dist.+pH+veget. | 7 | 42,154 | -68,66 | 27,402 | 5,75E-07 |
| edge.dist.+pH+temp.+veget. | 7 | 42,083 | -68,519 | 27,543 | 5,36E-07 |
| depth+diss.ox.+edge.dist.+pH+temp. | 8 | 42,886 | -67,623 | 28,439 | 3,42E-07 |
| depth+diss.ox.+pH+temp.+veget. | 8 | 42,853 | -67,556 | 28,505 | 3,31E-07 |
| depth+diss.ox.+edge.dist.+pH+veget. | 8 | 42,179 | -66,209 | 29,853 | 1,69E-07 |
| diss.ox.+edge.dist.+temp.+veget. | 7 | 40,795 | -65,944 | 30,118 | 1,48E-07 |
| depth+edge.dist.+pH+temp.+veget. | 8 | 41,839 | -65,53 | 30,532 | 1,20E-07 |
| depth+diss.ox.+edge.dist.+temp.+veget. | 8 | 41,218 | -64,287 | 31,775 | 6,46E-08 |
| diss.ox.+pH+edge.dist.+temp.+veget. | 8 | 39,456 | -60,764 | 35,298 | 1,11E-08 |
| depth+diss.ox.+edge.dist.+pH+temp.+veget. | 9 | 39,685 | -58,642 | 37,42 | 3,84E-09 |

**§20:** Summary of adjusted models produced by GLMM analysis relating local descriptors and functional redundancy in tadpole communities in ponds in the Spring (season with low hydric stress). Local descriptors: Depth (water column depth); Diss.ox. (dissolved oxygen); edge.dist. (edge distance); temp (water temperature); veget (percentage of aquatic vegetation cover).

**S20:** Summary of adjusted models produced by GLMM analysis relating local descriptors and functional redundancy in tadpole communities in ponds in the Spring (season with low hydric stress). Local descriptors: Depth (water column depth); Diss.ox. (dissolved oxygen); edge.dist. (edge distance); temp (water temperature); veget (percentage of aquatic vegetation cover).

| Model | df | logLik | AICc | $\Delta$ | weight |
| --- | --- | --- | --- | --- | --- |
| null | 3 | 126,91 | -247,7 | 0 | 0,9205 |
| temp. | 4 | 124,23 | -240,26 | 7,4385 | 0,0223 |
| diss.ox. | 4 | 124,18 | -240,17 | 7,5283 | 0,0213 |
| depth | 4 | 123,31 | -238,42 | 9,2721 | 0,0089 |
| pH | 4 | 123,26 | -238,34 | 9,361 | 0,0085 |
| veget. | 4 | 123,21 | -238,22 | 9,4762 | 0,0081 |
| edge.dist. | 4 | 123,18 | -238,17 | 9,5293 | 0,0078 |
| diss.ox.+temp. | 5 | 121,21 | -232,12 | 15,573 | 0,0004 |
| depth+temp. | 5 | 120,6 | -230,9 | 16,794 | 0,0002 |
| pH+temp. | 5 | 120,58 | -230,86 | 16,837 | 0,0002 |
| depth+diss.ox. | 5 | 120,56 | -230,83 | 16,866 | 0,0002 |
| diss.ox.+pH | 5 | 120,55 | -230,8 | 16,897 | 0,0002 |
| temp.+veget. | 5 | 120,5 | -230,7 | 16,995 | 0,0002 |
| diss.ox.+veget. | 5 | 120,48 | -230,66 | 17,032 | 0,0002 |
| diss.ox.+edge.dist. | 5 | 120,47 | -230,66 | 17,04 | 0,0002 |
| edge.dist.+temp. | 5 | 120,46 | -230,64 | 17,061 | 0,0002 |
| depth+pH | 5 | 119,67 | -229,05 | 18,644 | 8,23E-05 |
| depth+veget. | 5 | 119,61 | -228,92 | 18,776 | 7,71E-05 |
| depth+edge.dist. | 5 | 119,6 | -228,91 | 18,784 | 7,68E-05 |
| edge.dist.+pH | 5 | 119,57 | -228,85 | 18,844 | 7,45E-05 |
| pH+veget. | 5 | 119,57 | -228,84 | 18,854 | 7,41E-05 |
| edge.dist.+veget. | 5 | 119,48 | -228,67 | 19,028 | 6,79E-05 |
| depth+diss.ox.+temp. | 6 | 117,58 | -222,74 | 24,952 | 3,51E-06 |
| diss.ox.+pH+temp. | 6 | 117,56 | -222,71 | 24,984 | 3,46E-06 |
| diss.ox.+temp.+veget. | 6 | 117,48 | -222,55 | 25,143 | 3,19E-06 |
| diss.ox.+edge.dist.+temp. | 6 | 117,47 | -222,52 | 25,174 | 3,14E-06 |
| depth+pH+temp. | 6 | 116,96 | -221,5 | 26,198 | 1,88E-06 |
| depth+diss.ox.+pH | 6 | 116,93 | -221,45 | 26,245 | 1,84E-06 |
| depth+diss.ox.+edge.dist. | 6 | 116,88 | -221,35 | 26,347 | 1,75E-06 |
| diss.ox.+edge.dist.+pH | 6 | 116,88 | -221,35 | 26,348 | 1,75E-06 |
| depth+temp.+veget. | 6 | 116,88 | -221,34 | 26,352 | 1,74E-06 |
| depth+edge.dist.+temp. | 6 | 116,88 | -221,34 | 26,357 | 1,74E-06 |
| depth+diss.ox.+veget. | 6 | 116,86 | -221,31 | 26,389 | 1,71E-06 |
| pH+temp.+veget. | 6 | 116,85 | -221,29 | 26,409 | 1,70E-06 |
| diss.ox.+pH+veget. | 6 | 116,84 | -221,28 | 26,419 | 1,69E-06 |
| edge.dist.+pH+temp. | 6 | 116,84 | -221,26 | 26,433 | 1,68E-06 |
| diss.ox.+edge.dist.+veget. | 6 | 116,77 | -221,13 | 26,568 | 1,57E-06 |
| edge.dist.+temp.+veget. | 6 | 116,74 | -221,06 | 26,636 | 1,51E-06 |
| depth+edge.dist.+pH | 6 | 116,01 | -219,6 | 28,095 | 7,30E-07 |
| depth+pH+veget. | 6 | 115,97 | -219,53 | 28,163 | 7,05E-07 |
| depth+edge.dist.+veget. | 6 | 115,9 | -219,4 | 28,3 | 6,59E-07 |
| edge.dist.+pH+veget. | 6 | 115,87 | -219,34 | 28,362 | 6,39E-07 |
| depth+diss.ox.+pH+temp. | 7 | 113,94 | -213,33 | 34,369 | 3,17E-08 |
| depth+diss.ox.+edge.dist.+temp. | 7 | 113,88 | -213,21 | 34,489 | 2,98E-08 |
| depth+diss.ox.+temp.+veget. | 7 | 113,86 | -213,17 | 34,522 | 2,94E-08 |
| diss.ox.+edge.dist.+pH+temp. | 7 | 113,85 | -213,15 | 34,544 | 2,90E-08 |
| diss.ox.+pH+temp.+veget. | 7 | 113,84 | -213,13 | 34,57 | 2,87E-08 |
| diss.ox.+edge.dist.+temp.+veget. | 7 | 113,74 | -212,93 | 34,764 | 2,60E-08 |
| depth+diss.ox.+edge.dist.+pH | 7 | 113,3 | -212,05 | 35,646 | 1,67E-08 |
| depth+pH+temp.edge.dist. | 7 | 113,27 | -211,98 | 35,717 | 1,61E-08 |
| depth+pH+temp.+veget. | 7 | 113,24 | -211,92 | 35,772 | 1,57E-08 |
| depth+diss.ox.+pH+veget. | 7 | 113,23 | -211,91 | 35,784 | 1,56E-08 |
| depth+diss.ox.+edge.dist.+veget. | 7 | 113,18 | -211,82 | 35,881 | 1,49E-08 |
| diss.ox.+edge.dist.+pH+veget. | 7 | 113,18 | -211,8 | 35,896 | 1,48E-08 |
| depth+edge.dist.+temp.+veget. | 7 | 113,16 | -211,77 | 35,928 | 1,45E-08 |
| edge.dist.+pH+temp.+veget. | 7 | 113,11 | -211,67 | 36,025 | 1,38E-08 |
| depth+edge.dist.+pH+veget. | 7 | 112,31 | -210,07 | 37,625 | 6,22E-09 |
| depth+diss.ox.+edge.dist.+pH+temp. | 8 | 110,28 | -203,85 | 43,845 | 2,77E-10 |
| depth+diss.ox.+pH+temp.+veget. | 8 | 110,23 | -203,74 | 43,955 | 2,63E-10 |
| depth+diss.ox.+edge.dist.+temp.+veget. | 8 | 110,17 | -203,63 | 44,071 | 2,48E-10 |
| diss.ox.+edge.dist.+pH+temp.+veget. | 8 | 110,13 | -203,55 | 44,151 | 2,38E-10 |
| depth+diss.ox.+edge.dist.+pH+veget. | 8 | 109,61 | -202,5 | 45,195 | 1,41E-10 |
| depth+edge.dist.+pH+temp.+veget. | 8 | 109,55 | -202,39 | 45,303 | 1,34E-10 |
| depth+diss.ox.+edge.dist.+pH+temp.+veget. | 9 | 106,58 | -194,25 | 53,442 | 2,29E-12 |

**§21:** Summary of adjusted models produced by GLMM analysis relating local descriptors and functional redundancy in tadpole communities in ponds at Summer season (season with high hydric stress). Local descriptors: Depth (water column depth); Diss.ox. (dissolved oxygen); edge.dist. (edge distance); temp (water temperature); Veget (percentage of aquatic vegetation cover).

**S21:**
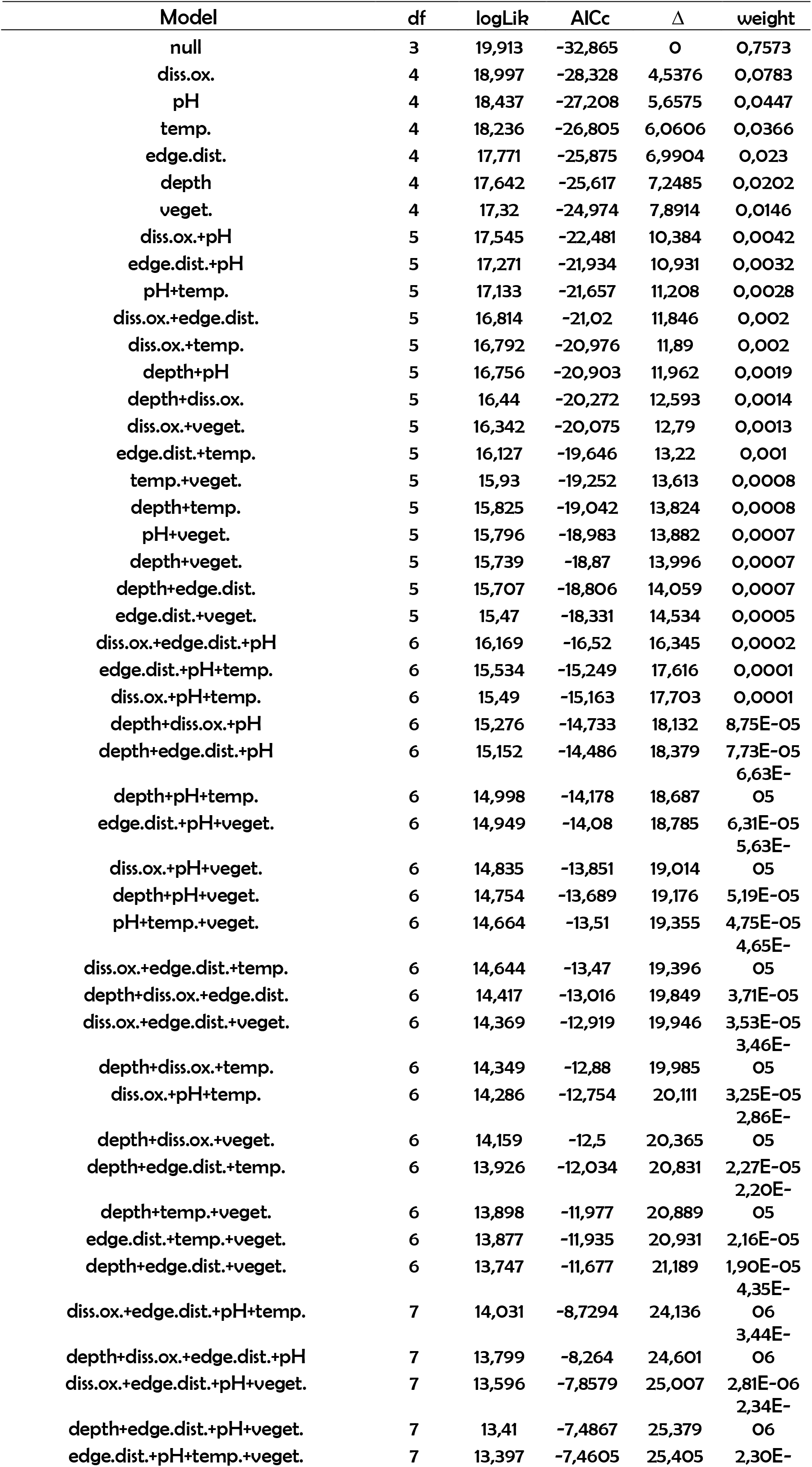
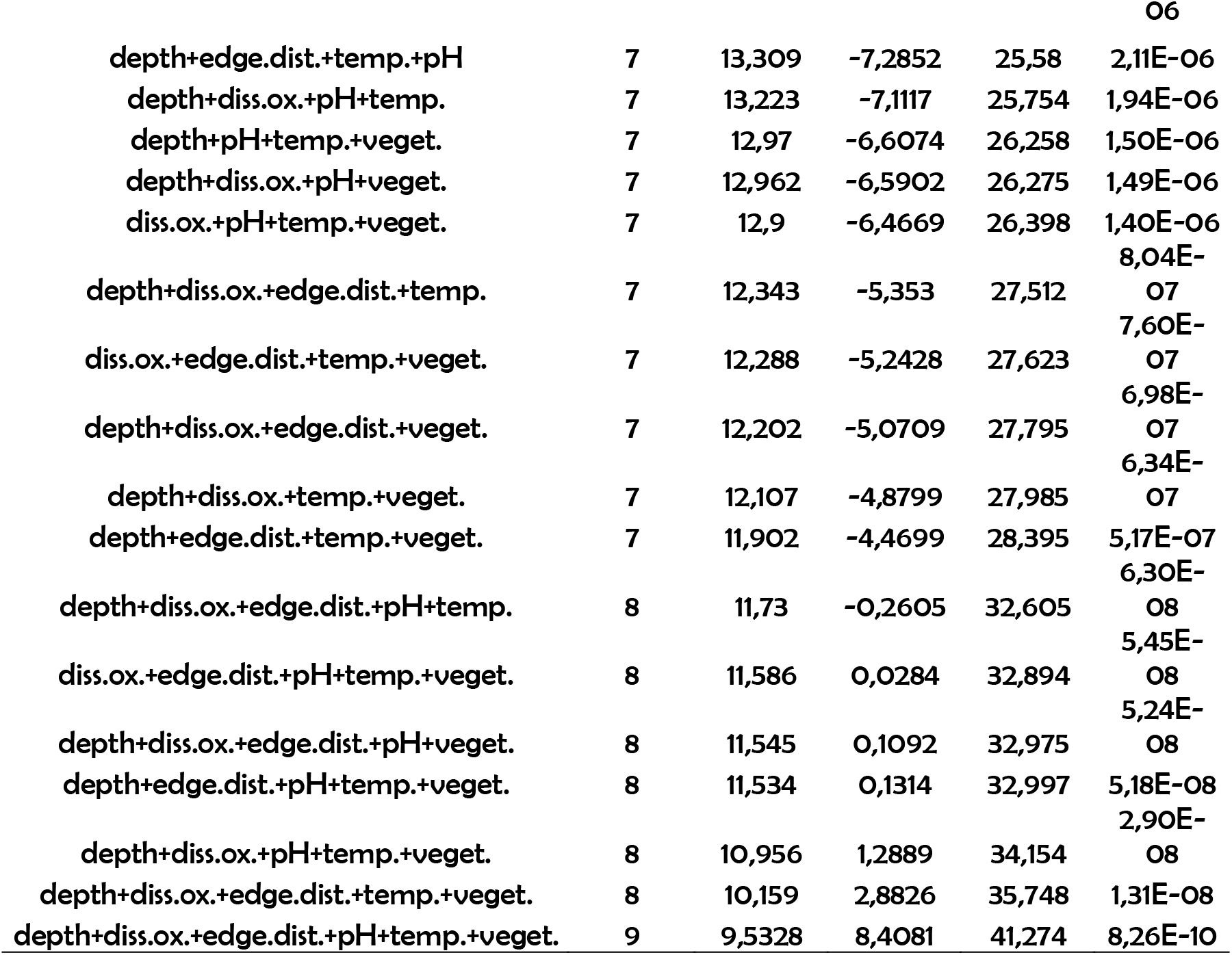
Summary of adjusted models produced by GLMM analysis relating local descriptors and functional redundancy in tadpole communities in ponds at Summer season (season with high hydrical stress). Local descriptors: Depth (water column depth); Diss.ox. (dissolved oxygen); edge.dist. (edge distance); temp (water temperature); veget (percentage of aquatic vegetation cover).

**S22:**
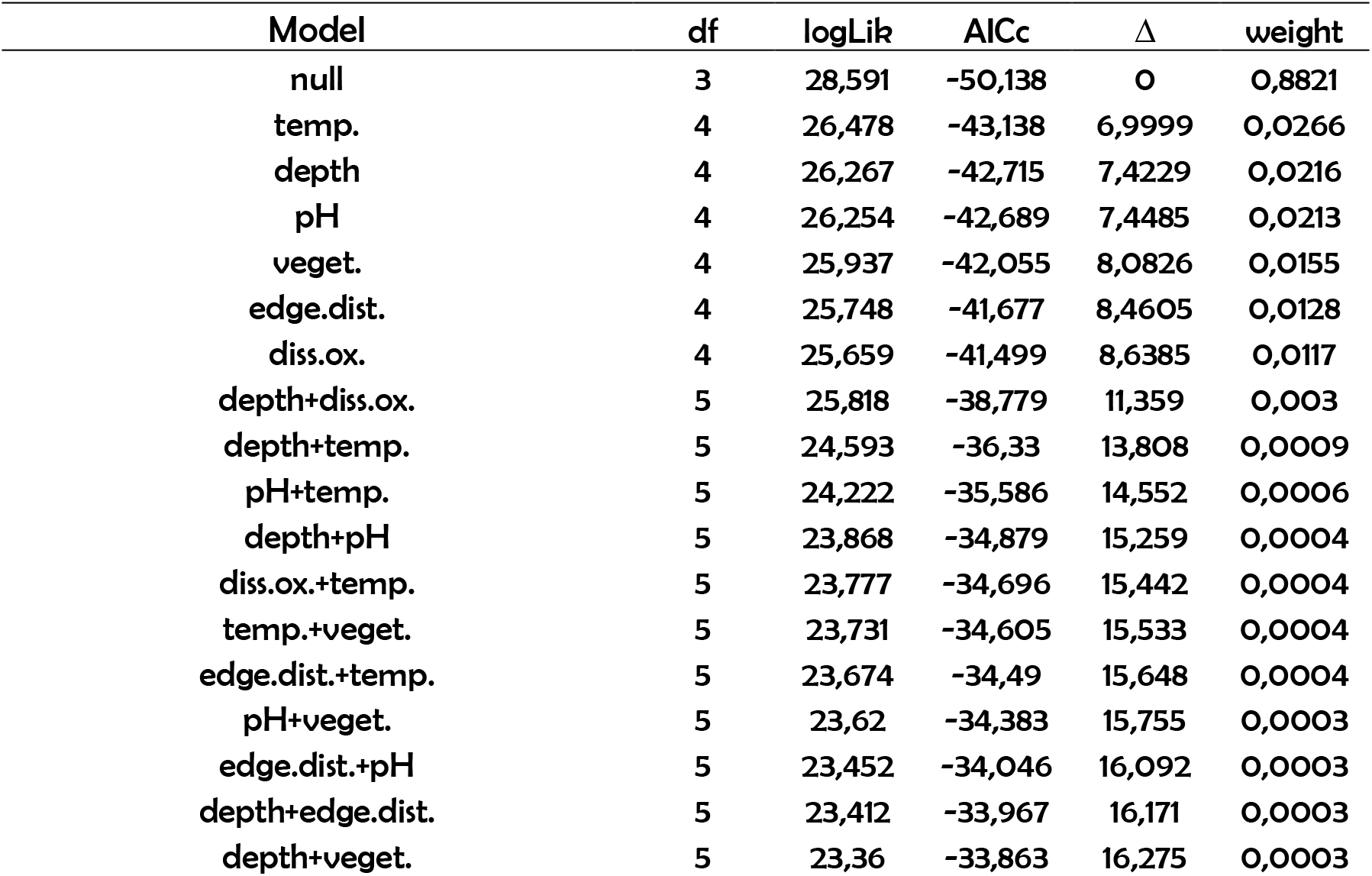
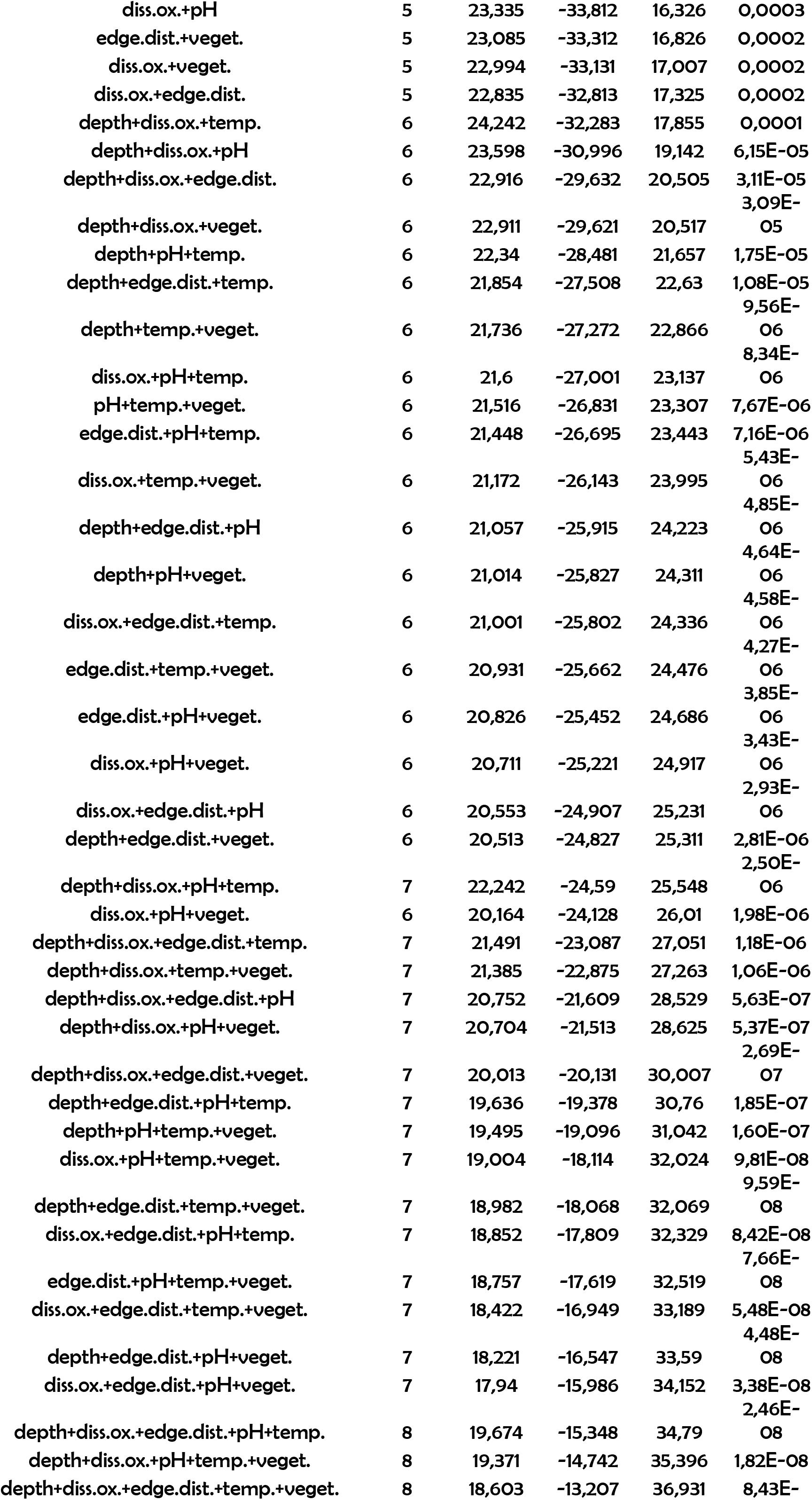
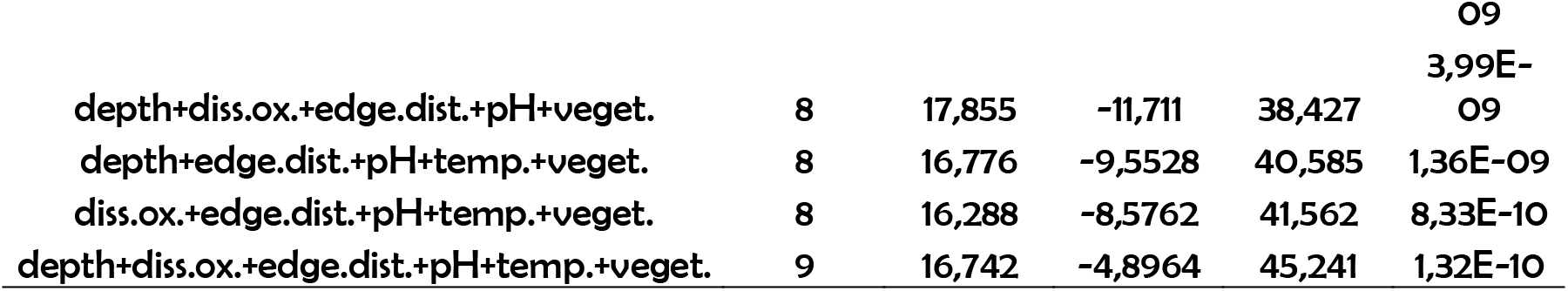
Summary of adjusted models produced by GLMM analysis relating local descriptors and functional redundancy in tadpole communities in ponds in the Autumn (season with high hydric stress). Local descriptors: Depth (water column depth); Diss.ox. (dissolved oxygen); edge.dist. (edge distance); temp (water temperature); veget (percentage of aquatic vegetation cover).

**S23:**
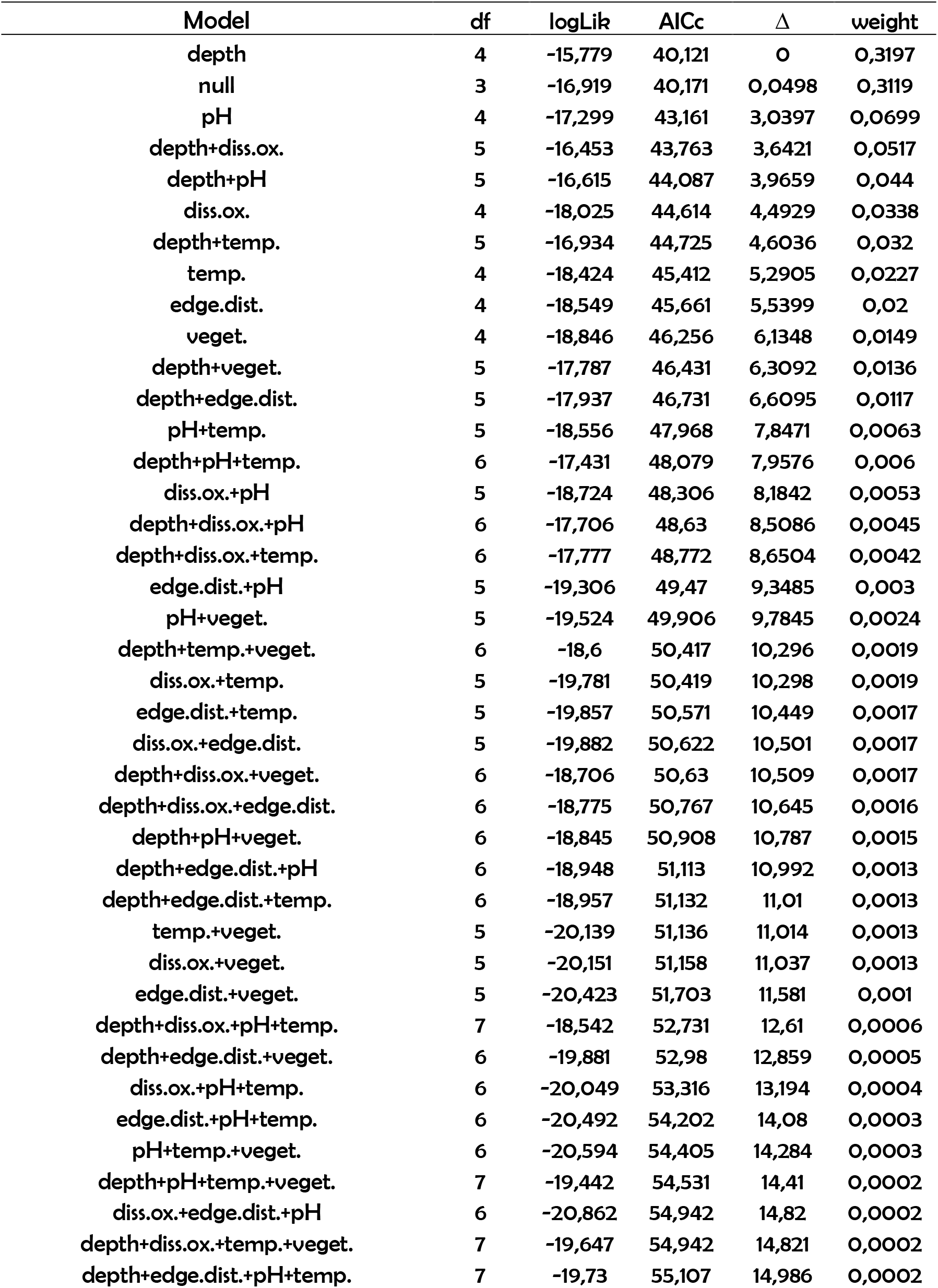
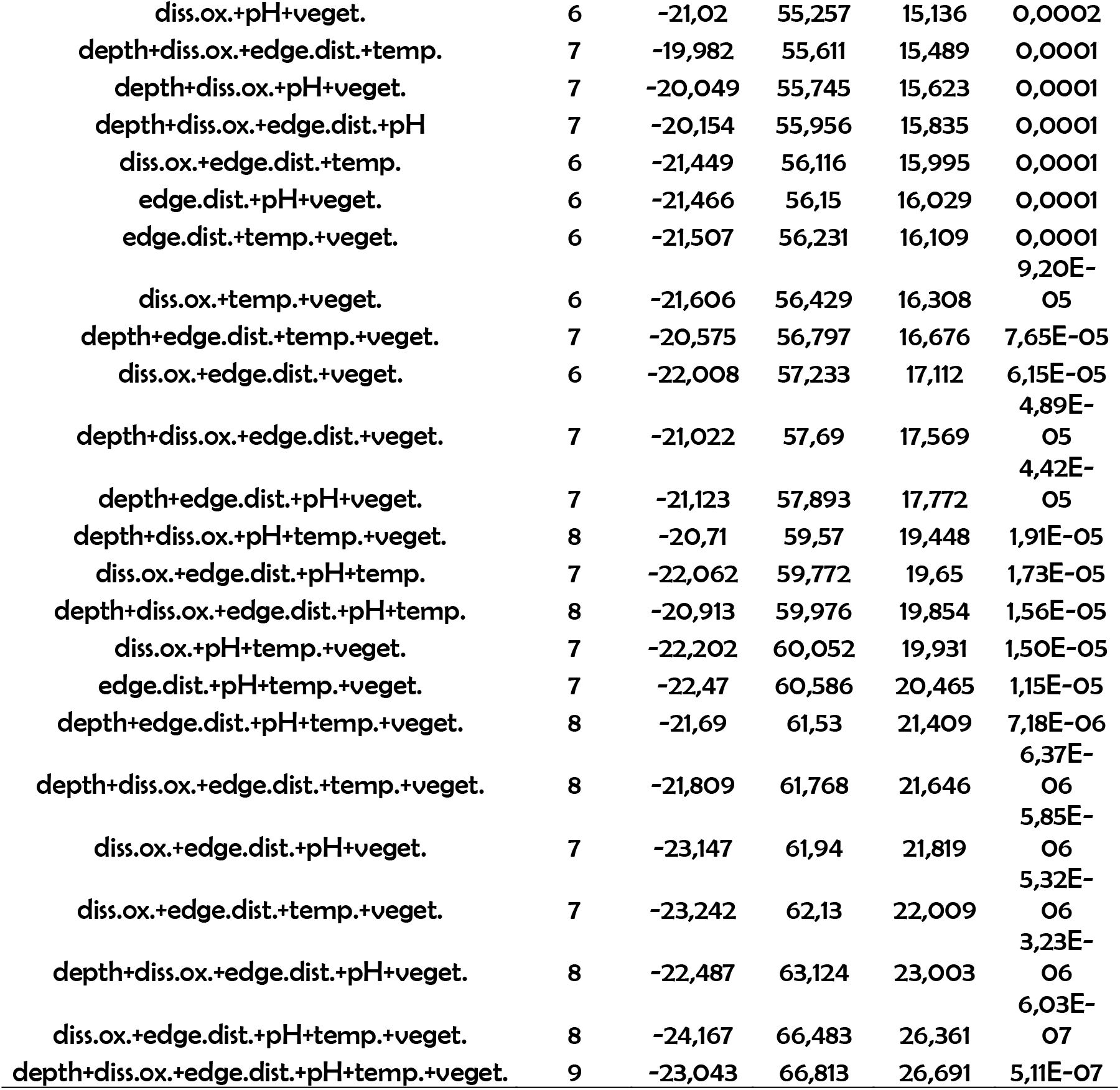
Summary of adjusted models produced by GLMM analysis relating local descriptors and taxonomic diversity in tadpole communities in ponds in the Winter (season with low hydric stress). Local descriptors: Depth (water column depth); Diss.ox. (dissolved oxygen); edge.dist. (edge distance); temp (water temperature); veget (percentage of aquatic vegetation cover).

**S24:**
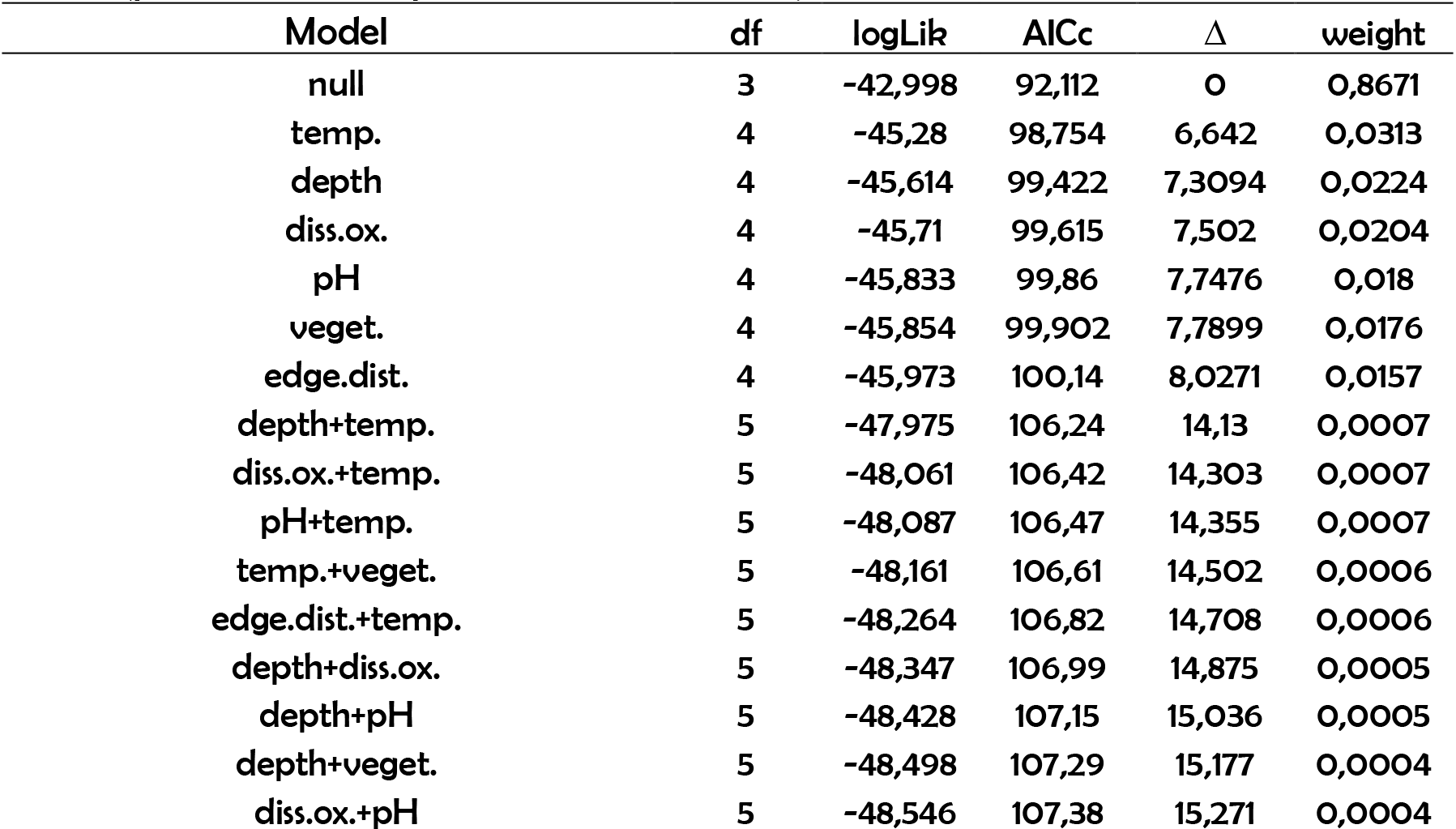
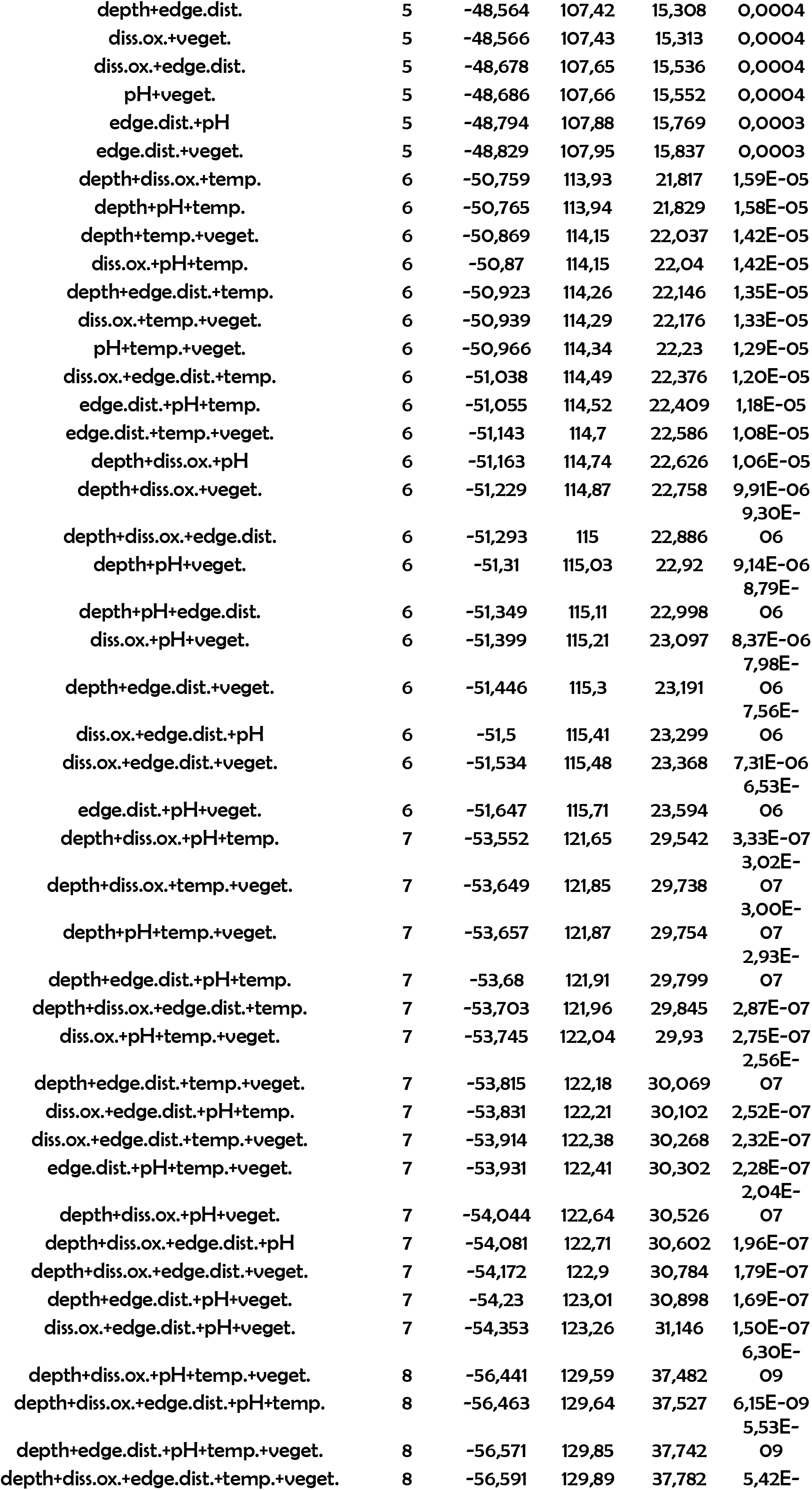
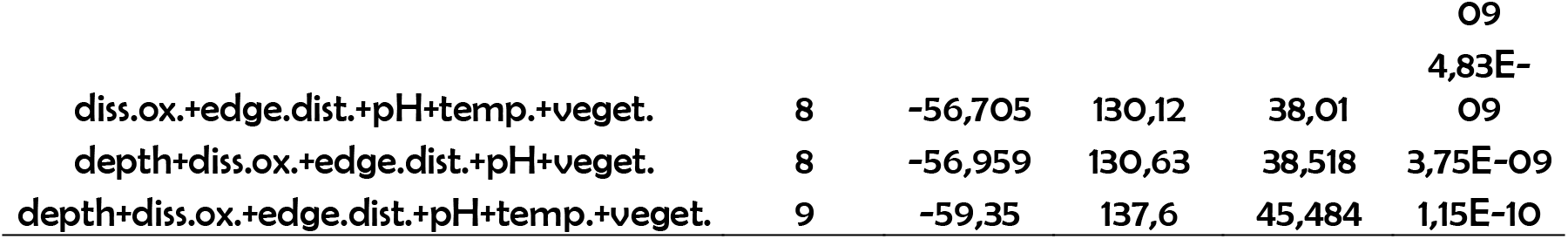
Summary of adjusted models produced by GLMM analysis relating local descriptors and taxonomic diversity in tadpole communities in ponds in the Spring season (season with low hydric stress). Local descriptors: Depth (water column depth); Diss.ox. (dissolved oxygen); edge.dist. (edge distance); temp (water temperature); veget (percentage of aquatic vegetation cover).

**S25:**
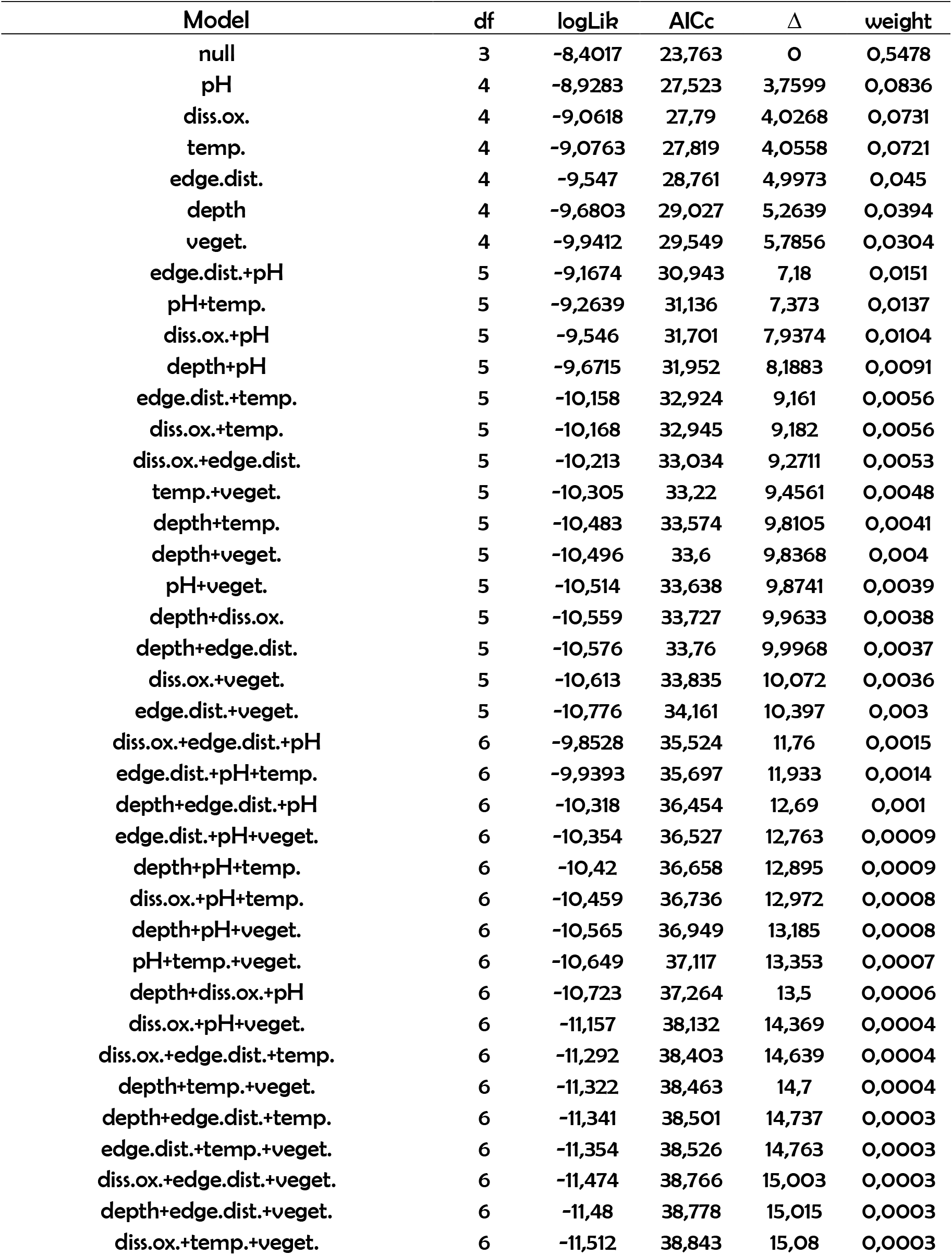
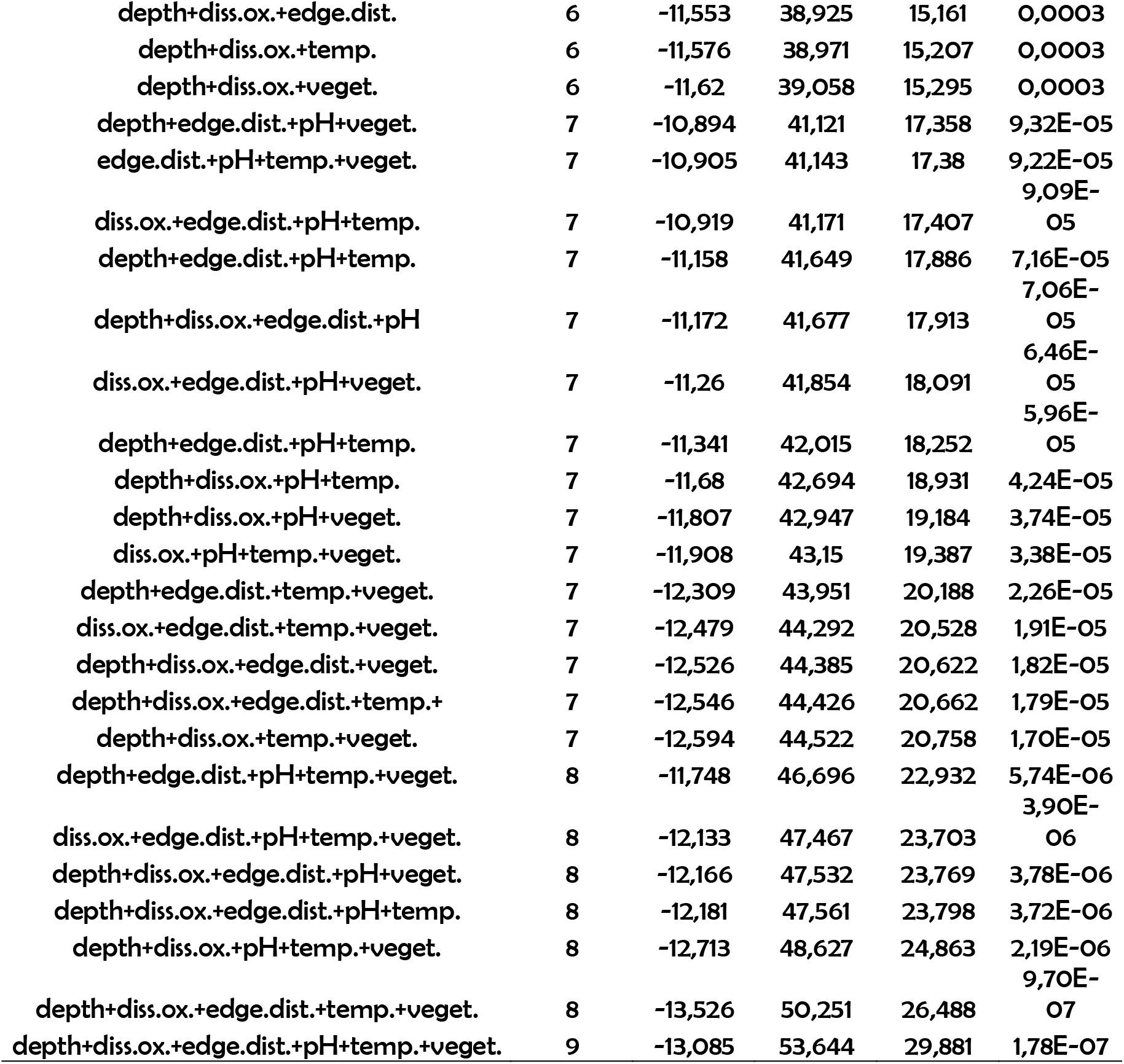
Summary of adjusted models produced by GLMM analysis relating local descriptors and taxonomic diversity in tadpole communities in ponds in the Summer (season with high hydric stress). Local descriptors: Depth (water column depth); Diss.ox. (dissolved oxygen); edge.dist. (edge distance); temp (water temperature); veget (percentage of aquatic vegetation cover).

**S26:** Summary of adjusted models produced by GLMM analysis relating local descriptors and taxonomic diversity in tadpole communities in ponds in the Autumn (season with high hydric stress). Local descriptors: Depth (water column depth); Diss.ox. (dissolved oxygen); edge.dist. (edge distance); temp (water temperature); veget (percentage of aquatic vegetation cover).

| Model | df | logLik | AICc | $\Delta$ | weight |
| --- | --- | --- | --- | --- | --- |
| null | 3 | 2,8556 | 1,3323 | 0 | 0,7088 |
| temp. | 4 | 1,7206 | 6,3771 | 5,0448 | 0,0569 |
| veget. | 4 | 1,6662 | 6,4858 | 5,1535 | 0,0539 |
| pH | 4 | 1,4785 | 6,8611 | 5,5288 | 0,0447 |
| edge.dist. | 4 | 1,3789 | 7,0605 | 5,7281 | 0,0404 |
| depth | 4 | 1,1022 | 7,6138 | 6,2815 | 0,0307 |
| diss.ox. | 4 | 0,894 | 8,0301 | 6,6978 | 0,0249 |
| depth+diss.ox. | 5 | 0,6535 | 11,55 | 10,218 | 0,0043 |
| pH+temp. | 5 | 0,6226 | 11,612 | 10,28 | 0,0042 |
| temp.+veget. | 5 | 0,3968 | 12,063 | 10,731 | 0,0033 |
| pH+veget. | 5 | 0,3355 | 12,186 | 10,854 | 0,0031 |
| edge.dist.+temp. | 5 | 0,1534 | 12,55 | 11,218 | 0,0026 |
| edge.dist.+pH | 5 | 0,1463 | 12,564 | 11,232 | 0,0026 |
| depth+temp. | 5 | 0,1399 | 12,577 | 11,245 | 0,0026 |
| edge.dist.+veget. | 5 | 0,0626 | 12,732 | 11,4 | 0,0024 |
| diss.ox.+temp. | 5 | 0,0363 | 12,785 | 11,452 | 0,0023 |
| depth+pH | 5 | -0,1318 | 13,121 | 11,788 | 0,002 |
| diss.ox.+veget. | 5 | -0,2885 | 13,434 | 12,102 | 0,0017 |
| depth+veget. | 5 | -0,3208 | 13,499 | 12,166 | 0,0016 |
| depth+edge.dist. | 5 | -0,4008 | 13,659 | 12,326 | 0,0015 |
| diss.ox.+pH | 5 | -0,465 | 13,787 | 12,455 | 0,0014 |
| diss.ox.+edge.dist. | 5 | -0,5823 | 14,022 | 12,689 | 0,0012 |
| depth+diss.ox.+pH | 6 | -0,1791 | 16,558 | 15,226 | 0,0004 |
| depth+diss.ox.+temp. | 6 | -0,3363 | 16,873 | 15,54 | 0,0003 |
| pH+temp.+veget. | 6 | -0,7316 | 17,663 | 16,331 | 0,0002 |
| depth+diss.ox.+veget. | 6 | -0,753 | 17,706 | 16,374 | 0,0002 |
| edge.dist.+pH+temp. | 6 | -0,8216 | 17,843 | 16,511 | 0,0002 |
| depth+diss.ox.+edge.dist. | 6 | -0,8493 | 17,899 | 16,566 | 0,0002 |
| depth+pH+temp. | 6 | -0,8859 | 17,972 | 16,64 | 0,0002 |
| diss.ox.+pH+temp. | 6 | -0,8975 | 17,995 | 16,663 | 0,0002 |
| diss.ox.+temp.+veget. | 6 | -1,0761 | 18,352 | 17,02 | 0,0001 |
| edge.dist.+pH+veget. | 6 | -1,1318 | 18,464 | 17,131 | 0,0001 |
| depth+temp.+veget. | 6 | -1,2096 | 18,619 | 17,287 | 0,0001 |
| depth+edge.dist.+temp. | 6 | -1,2224 | 18,645 | 17,312 | 0,0001 |
| edge.dist.+temp.+veget. | 6 | -1,2277 | 18,655 | 17,323 | 0,0001 |
| depth+edge.dist.+pH | 6 | -1,4288 | 19,058 | 17,725 | 0,0001 |
| diss.ox.+edge.dist.+temp. | 6 | -1,5369 | 19,274 | 17,941 | 9,01E-05 |
| depth+pH+veget. | 6 | -1,5397 | 19,279 | 17,947 | 8,98E-05 |
| diss.ox.+pH+veget. | 6 | -1,6119 | 19,424 | 18,091 | 8,36E-05 |
| diss.ox.+edge.dist.+pH | 6 | -1,8036 | 19,807 | 18,475 | 6,90E-05 |
| diss.ox.+edge.dist.+veget. | 6 | -1,8804 | 19,961 | 18,629 | 6,39E-05 |
| depth+edge.dist.+veget. | 6 | -1,9171 | 20,034 | 18,702 | 6,16E-05 |
| depth+diss.ox.+pH+temp. | 7 | -0,9285 | 21,752 | 20,419 | 2,61E-05 |
| depth+diss.ox.+edge.dist.+pH | 7 | -1,282 | 22,459 | 21,126 | 1,83E-05 |
| depth+diss.ox.+pH+veget. | 7 | -1,572 | 23,039 | 21,706 | 1,37E-05 |
| depth+diss.ox.+temp.+veget. | 7 | -1,6667 | 23,228 | 21,896 | 1,25E-05 |
| depth+diss.ox.+edge.dist.+temp. | 7 | -1,6685 | 23,232 | 21,899 | 1,24E-05 |
| depth+edge.dist.+pH+temp. | 7 | -2,013 | 23,921 | 22,588 | 8,82E-06 |
| diss.ox.+pH+temp.+veget. | 7 | -2,073 | 24,041 | 22,709 | 8,31E-06 |
| edge.dist.+pH+temp.+veget. | 7 | -2,2441 | 24,383 | 23,051 | 7,00E-06 |
| depth+pH+temp.+veget. | 7 | -2,2764 | 24,448 | 23,115 | 6,78E-06 |
| depth+diss.ox.+edge.dist.+veget. | 7 | -2,3499 | 24,595 | 23,262 | 6,30E-06 |
| diss.ox.+edge.dist.+pH+temp. | 7 | -2,3811 | 24,657 | 23,325 | 6,10E-06 |
| depth+edge.dist.+temp.+veget. | 7 | -2,6924 | 25,28 | 23,947 | 4,47E-06 |
| diss.ox.+edge.dist.+temp.+veget. | 7 | -2,7505 | 25,396 | 24,063 | 4,22E-06 |
| depth+edge.dist.+pH+veget. | 7 | -2,9661 | 25,827 | 24,495 | 3,40E-06 |
| diss.ox.+edge.dist.+pH+veget. | 7 | -3,0797 | 26,054 | 24,722 | 3,04E-06 |
| depth+diss.ox.+edge.dist.+pH+temp. | 8 | -1,7455 | 27,491 | 26,159 | 1,48E-06 |
| depth+diss.ox.+pH+temp.+veget. | 8 | -2,307 | 28,614 | 27,282 | 8,44E-07 |
| depth+diss.ox.+edge.dist.+pH+veget. | 8 | -2,8342 | 29,668 | 28,336 | 4,98E-07 |
| depth+diss.ox.+edge.dist.+temp.+veget. | 8 | -3,1266 | 30,253 | 28,921 | 3,72E-07 |
| depth+edge.dist.+pH+temp.+veget. | 8 | -3,549 | 31,098 | 29,766 | 2,44E-07 |
| diss.ox.+edge.dist.+pH+temp.+veget. | 8 | -3,666 | 31,332 | 30 | 2,17E-07 |
| depth+diss.ox.+edge.dist.+pH+temp.+veget. | 9 | -3,2986 | 35,185 | 33,853 | 3,16E-08 |

**§27** - Reproductive phenology of adults and temporal distribution of tadpoles recorded in the present study. The spiked bars represent the pattern of adult activity, and numbers represent the abundance of tadpoles in each season of the year. The (\*) represent the presence of tadpoles registered in our study area.

**S27.**
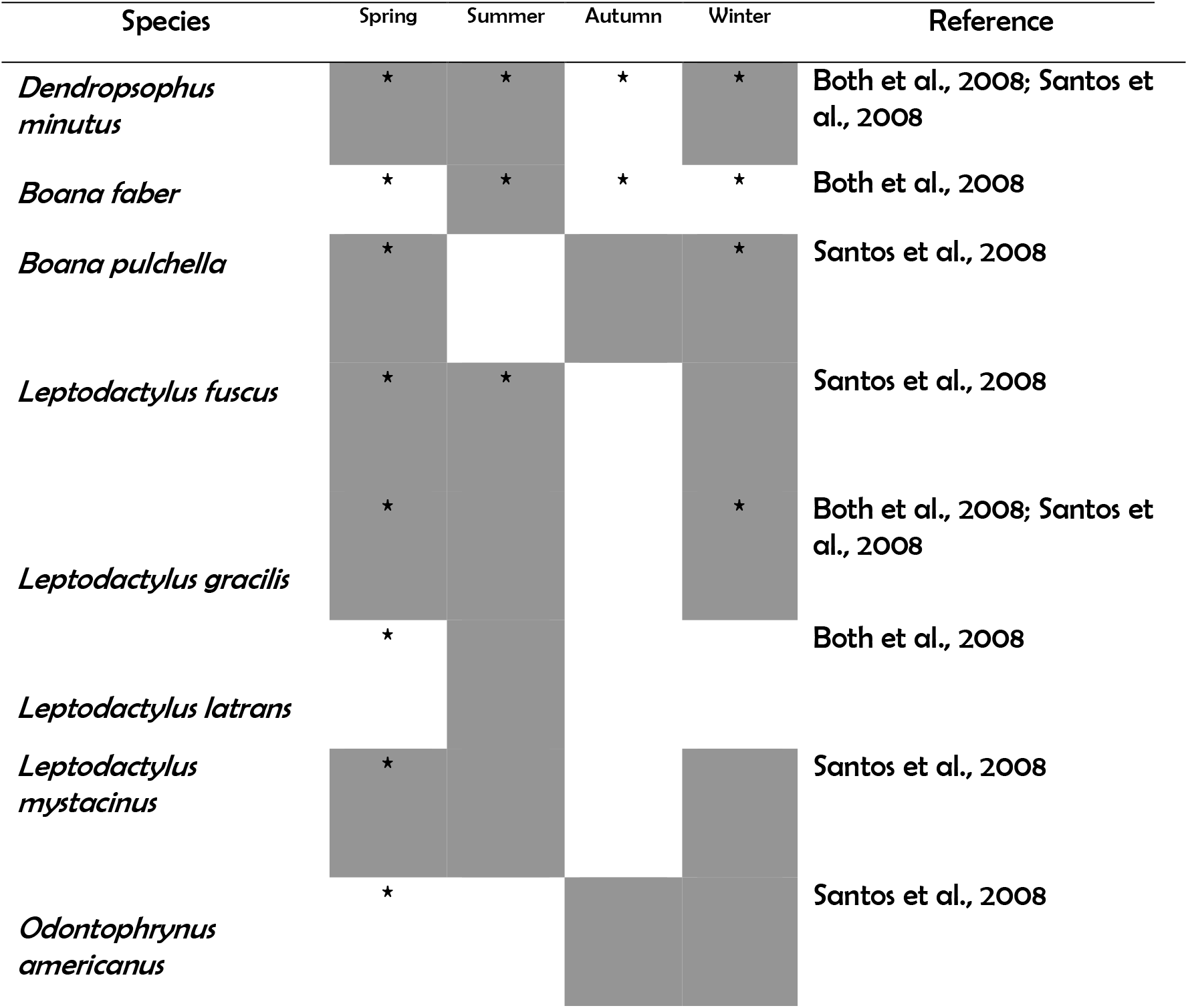
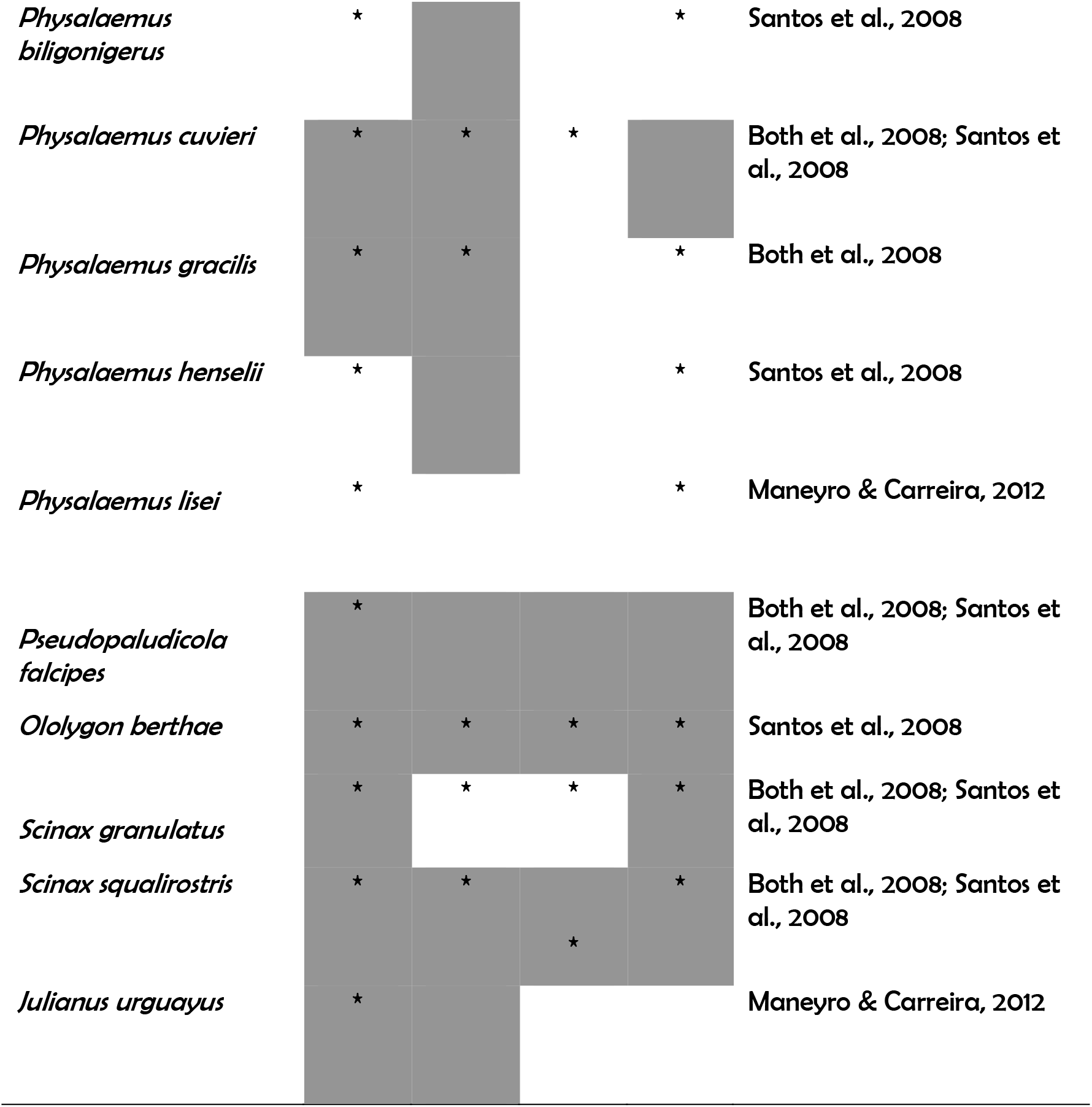
Reproductive phenology of adults and temporal distribution of tadpoles recorded in the present study. The spiked bars represent the pattern of adult activity, and numbers represent the abundance of tadpoles in each season of the year. The (*) represent the presence of tadpoles registered in our study area.

